# Quartet-based species tree methods enable fast and consistent tree of blobs reconstruction under the network multispecies coalescent

**DOI:** 10.1101/2025.11.05.686850

**Authors:** Junyan Dai, Yunheng Han, Erin K. Molloy

## Abstract

Hybridization between species is an important force in evolution, commonly modeled by the network multispecies coalescent. Reconstructing evolutionary histories under this model is computationally challenging, even for level-1 networks where hybridization events are isolated. Divide-and-conquer is a promising path forward, but current methods with statistical guarantees rely on an estimated *tree of blobs* (TOB) for the network, which compresses the non-tree-like parts into single vertices. TOB reconstruction is itself challenging, with the only available method TINNiK having time complexity *O*(*n*^5^ + *n*^4^*k*) for *k* genes and *n* species. Here, we present a new framework for scalable TOB reconstruction with statistical guarantees. Our approach operates by (1) seeking a refinement of the TOB and then (2) contracting edges in it. For step (1), we show that any optimal solution to Weighted Quartet Consensus is a TOB refinement almost surely, as the number of genes goes to infinity, motivating the use of methods, such as ASTRAL or TREE-QMC. For step (2), we show that applying the same hypothesis tests as TINNiK to just *O*(*n*) four-taxon subsets around each edge is sufficient for statistical consistency when the underlying network is level-1. Leveraging TREE-QMC for the first step gives our method time complexity *O*(*n*^3^*k*) and its name: TOB-QMC. On simulated data, TOB-QMC typically matches or exceeds TINNiK in accuracy while being more scalable. TOB-QMC also enables fast exploration of non-tree-like evolution, as demonstrated through re-analysis of three phylogenomic data sets. Lastly, our study clarifies the theoretical utility of quartet-based species tree methods in the context of hybridization, which is critical given the recent result that ASTRAL can be misleading.

## 1 Introduction

Hybridization between species or admixture between populations are important forces in evolution; however, they complicate the reconstruction of evolutionary histories, requiring them to be represented as phylogenetic networks rather than trees. In this context, phylogenetic networks (also called species or population networks) govern the ancestry of unlinked genomic regions (i.e., gene genealogies or gene trees), as modeled by the network multi-species coalescent (NMSC) [39]. Gene trees can differ from the display trees of the network due to *incomplete lineage sorting (ILS)*, which can occur when gene lineages enter the same ancestral population but fail to coalesce, and due to *reticulation events*, which allow gene lineages to trace back to different ancestral populations.

Despite significant progress over the last decade, phylogenetic network reconstruction under the NMSC continues to be a major scientific challenge, even for level-1 networks where hybridization events are isolated from each other. Perhaps the most statistically rigorous methods developed to date are those based on likelihood functions that account for both gene trees evolving within a species network under the NMSC [39, 42] and molecular sequences evolving down each gene tree [13], integrating over all possible gene trees [40]. The latter step can be avoided by estimating gene trees from molecular sequences prior to network reconstruction, but even the simplified likelihood calculation can be prohibitively expensive. This has motivated the development of pseudo-likelihood methods [41, 35], although SNaQ [35], perhaps the most popular pseudolikelihood method for level-1 (semi-directed) networks, is still limited to around 30 species [23].

Given scalability limitations, researchers sometimes add edges representing gene flow to a fixed species tree, typically estimated with ASTRAL [26] (e.g., Fig. 2 in [11]; also see [10]). ASTRAL is a heuristic for the NP-hard Weighted Quartet Consensus (WQC) problem [24], which can be framed as weighting quartets (i.e., unrooted trees on four species) according to their frequencies in the input gene trees and then seeking a species tree to maximize the total weight of the quartets it induces. Dinh and Baños (2025) recently showed that this approach can be *positively misleading* [12] (also see [27]). This negative result impacts numerous prior studies using ASTRAL in the presence of gene flow and suggests scalable phylogenetic network methods are urgently needed.

Divide-and-conquer is a promising path forward [18, 44, 2, 23]. One approach, taken by the recently developed InPhyNet method [23], is to partition species into subsets, estimate a level-1 (semi-directed) network on each subset (e.g., with SNaQ), and then combine the results into a level-1 (semi-directed) network on the complete species set, similar to disjoint tree mergers [28, 29]. InPhyNet [23] achieves a correctness guarantee from partitioning species according to the tree of blobs (TOB) [15] for the network, which compresses the non-treelike parts as single vertices, called blob vertices. This process is reversed in the divide-and-conquer method NANUQ+ [3], which enables statistically consistent expansion of blob vertices in the TOB, producing a level-1 (semi-directed) network. Thus, InPhyNet and NANUQ+ require fast and accurate TOB estimation for empirical performance and consistent TOB estimation for theoretical guarantees.

To our knowledge, the first and only available method for TOB reconstruction under the NMSC is TINNiK [1]. TINNiK is statistically consistent and has time complexity *O*(*n*^4^*k* + *n*^5^), where *k* is the number of input gene trees and *n* is the number of species. The first term comes from computing quartet weights, which are needed to perform hypothesis tests on all subsets of four species, and the second term comes from updating the test results so that encoding them as a distance matrix, applying Neighbor Joining [34], and contracting zero-length edges yields the TOB. However, as TINNiK’s asymptotic runtime would suggest, it cannot scale to large numbers of species. To address this issue, we present a new framework for scalable TOB reconstruction with statistical guarantees, implemented in the software package TOB-QMC.

## 2 Results

### 2.1 TOB-QMC Method Overview

To present our approach, take the phylogenetic network shown in Figure 1A as an example. Internal vertices with in-degree greater than 1 represent hybrid species or admixed populations, whereas internal vertices with in-degree 1 represent speciation events or population splits. Each edge in the network is either a cut edge (meaning its deletion disconnects the network) or a blob edge. The contraction of all blob edges in the network yields the tree of blobs (TOB) (Fig. 1B), after suppressing all degree 2 vertices. A TOB refinement has additional edges, called false positives (FPs), such that contracting them yields the TOB (Fig. 1C).

**Fig. 1:**
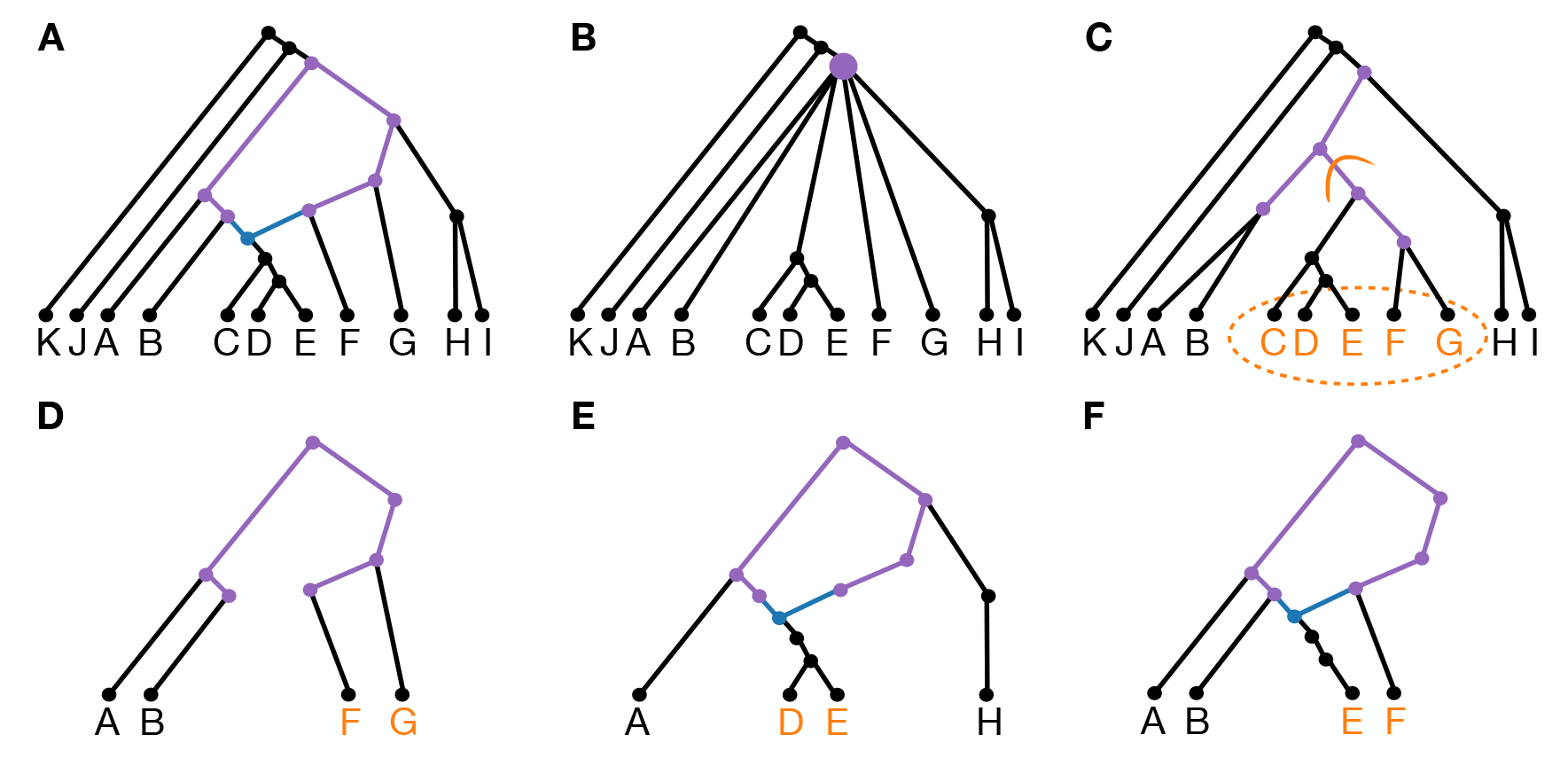
**A)** Directed, binary, level-1 phylogenetic network *N* ^+^. Reticulation edges and vertices are blue; all other edges and vertices in blob ℬ are purple. **B)** Tree of blobs (TOB) *T*_*N*_ for *N* ^+^ with the vertex *v*_*B*_ created by contracting edges in ℬ highlighted purple. **C)** A refinement of *T*_*N*_ that is <u>not</u> displayed by *N* ^+^. Edges that refine *v*_*B*_ are purple. To decide whether to contract the edge marked in orange, an oracle for treelikeness is applied to four taxa, two from each side of its induced bipartition. **D)** Subnetwork on *A, B, F, G*; its TOB is a quartet so the oracle returns treelike (note that the TOB restricted to *A, B, F, G* is a star, called a B-quartet). **E)** Subnetwork on *A, H, D, E*; its TOB is a quartet so the oracle returns treelike (note that the TOB restricted to *A, H, D, E* is a quartet, called a T-quartet). **F)** Subnetwork on *A, B, E, F*; its TOB tree is a star so the oracle returns non-treelike (note that the TOB restricted to *A, B, E, F* is a B-quartet).

**Fig. 2:**
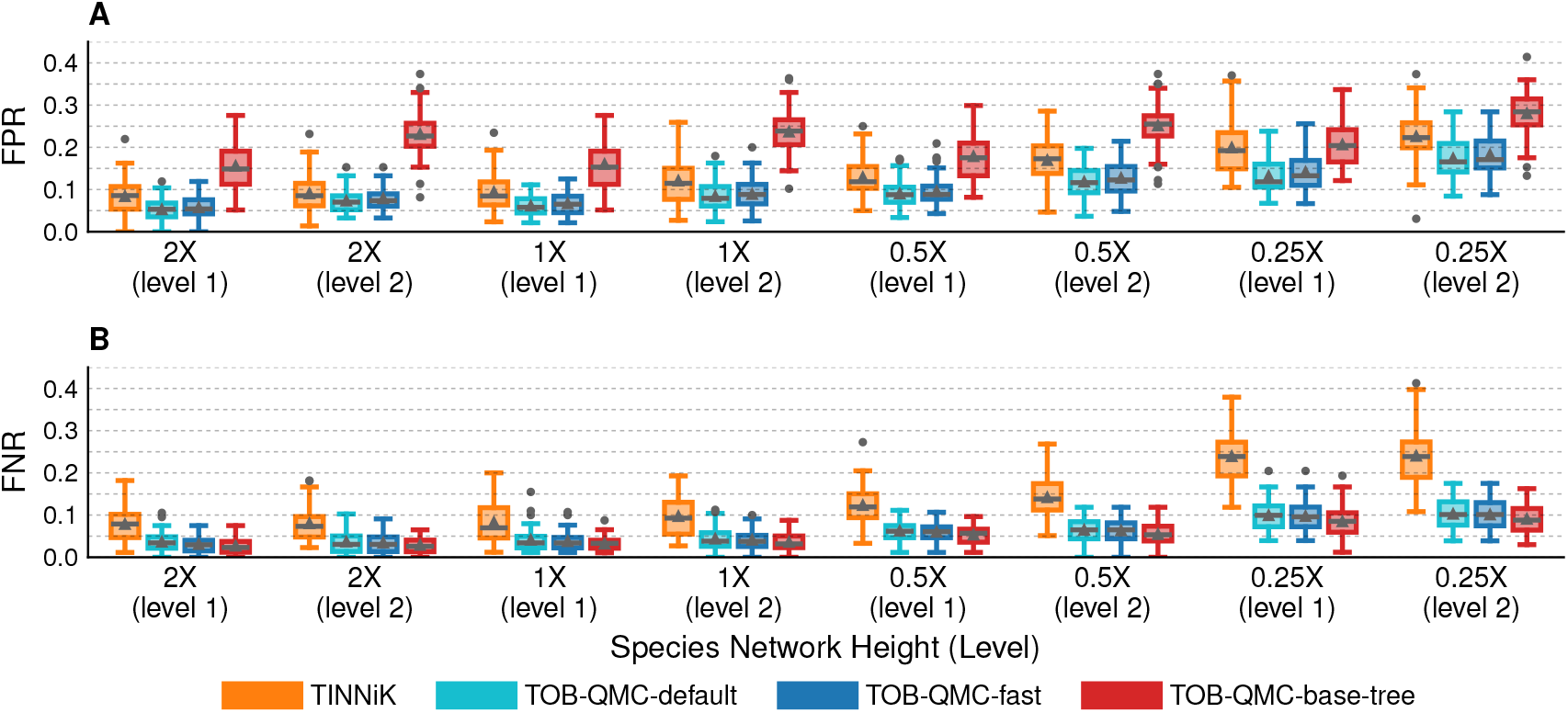
FPR and FNR for TOB reconstruction methods on 100-taxon data sets with estimated gene trees. Model condition on *x*-axis indicates network level and branch scaling factor (lower factor is higher ILS). Triangles indicate means, bars indicate medians, and circles indicate outliers.

Our proposed framework reverses this process first seeking a TOB refinement and second contracting edges in it. For step (1), we seek a solution to the NP-hard Weighted Quartet Consensus (WQC) problem using a fast quartet-based species tree method (e.g., TREE-QMC [16] or ASTRAL). This approach is theoretically grounded because we show that any optimal solution to WQC is a TOB refinement almost surely, as the number of (error-free) gene trees generated under the NMSC goes to infinity (Theorem 1). For step (2), we leverage the same hypothesis testing framework as TINNiK. The input to the testing framework is the number of gene trees inducing each of the three possible quartet trees on four species, called *quartet concordance factors* (qCFs), and the output is whether the evolutionary relationship among the four taxa is treelike, meaning the TOB of their <u>subnetwork</u> is a binary quartet tree rather than an unresolved star tree (see Methods for details). Importantly, even if four taxa are related through blob edges in the network, their relationship in the subnetwork can still be treelike (Figs. 1D–F). We show that applying an oracle for treelikeness to all four-taxon subsets around each edge, taking two taxa from either side, correctly detects FP edges in a TOB refinement for level-1 networks (Corollary 1). Moreover, we design search strategies to reduce the number of four-taxon subsets tested per edge from *O*(*n*^4^) to *O*(*n*) while preserving the correctness guarantee (Theorem 2). Combining our linear-time search algorithm, called 3f1a, with TINNiK’s hypothesis testing framework enables *statistically consistent* TOB reconstruction for level-1 networks under the NMSC, meaning that as the number of gene trees goes to infinity, the probability that our approach returns the true TOB goes to 1. Below we summarize the assumptions about the underlying network made in our proofs of consistency; see Methods for details.

– Standard Assumptions (also required by TINNiK):
  1. Network is binary.
  2. Non-reticulation edges have positive lengths in coalescent units; reticulation edges have lengths greater than or equal to 0.
  3. Inheritance proportions on reticulation edges incident to the same child sum to 1 and take on values between 0 and 1 (non-inclusive); inheritance proportions on non-reticulation edges equal 1.
– Assumptions for refinement tree reconstruction:
  4. Network is T-quartet-nonanomalous (Definition 1).
– Assumptions for FP edge detection:
  5. Network is level-1.
  6. Assumptions inherited from the oracle for treelikeness (i.e., TINNiK’s hypothesis tests):
    6a. Expected qCFs are distinct for every subset of four taxa whose relationship is non-treelike (i.e., the TOB of the subnetwork is an unresolved star tree).
    6b. Network is class 1 quartet-nonanomalous, i.e., for every subset of four taxa whose relationship is treelike, the topology of the highest expected qCF matches the TOB of the subnetwork (T3 option only).

The above assumptions are standard and shared by TINNiK, except condition 5. TINNiK’s consistency guarantee requires condition 6a; otherwise, the null hypothesis of treelikeness may be incorrectly accepted (i.e., the test predicts the relationship is treelike when it is not). Condition 6b regarding anomalous quartets is required by TINNiK when using the T3 test option; otherwise, the null may be incorrectly rejected (i.e., the test predicts the relationship is not treelike when it actually is); our assumptions in refinement tree reconstruction are no stricter (see Definition 1 for details). Anomalous quartets can occur when the subnetwork on four taxa contains a blob with three pendant cut edges and two taxa descending from the hybrid vertex for some numerical parameter settings (Fig. 1E) [5]. These so-called 3_2_-cycles also challenge quartet-based network inference methods; see [35, 6]. Nevertheless, the T3 option is the default in TINNiK due to its strong empirical performance [1]. Unlike TINNiK, our statistical guarantees for FP detection are restricted to level-1 networks; however, this comes with improved scalability: time complexity of *O*(*n*^3^*k*) versus *O*(*n*^4^*k* + *n*^5^).

We implemented our approach on top of the quartet-based species tree method TREE-QMC [16, 17], which is used to build a refinement tree for step (1), giving our method it name: TOB-QMC. Alternatively, a precomputed refinement tree (e.g., built with ASTRAL) can be given as input. **TOB-QMC-exhaustive** tests for treelikeness among all four-taxon subsets around each edge in the refinement tree, storing the minimum *p*-value found and the minimizer four-taxon subset. **TOB-QMC-3f1a** executes the 3f1a search algorithm (see Methods and Algorithm 1 in Supplement). **TOB-QMC-fast** and **TOB-QMC-default** perform an expanded heuristic search, stopping when the maximum number of iterations is reached, either 2*n*^2^ and *n*^2^*/*4, respectively, where *n* is the number of taxa, or a local optima is reached (see Methods and Algorithm 2 in Supplement). After performing the search around each edge, edges are contracted if the null hypothesis of the treelikeness test is rejected (i.e., *p*_*min*_ *< α*). However, the treelikeness test cannot be used when the evolutionary relationship among the four species is a polytomy (i.e., the subnetwork is a star tree). To rule out this scenario, TINNiK’s star test is applied to the minimizer four-taxon subset, and if the null hypothesis is <u>not</u> rejected (i.e., *p*_*min*_ ≥ *β*), the edge is contracted. Thus, higher *α* and lower *β* result in more edges being contracted. The user is required to set the significance thresholds *α, β*, same as TINNiK. However, by storing minimum *p*-values found and taxon minimizers for each edge, the impact of *α, β* can be evaluated offline, avoiding full re-computation of the TOB.

### 2.2 Method Evaluation

We evaluate TOB-QMC and TINNiK using simulations so that the true network and its TOB are known.

#### Data simulation

Data sets were simulated following the protocol from [23]. First, networks with 50, 100, and 200 taxa were simulated via a birth-death-hybridization process using SiPhyNetwork [21]. For each number of taxa, 50 level-1 and 50 level-2 networks were selected (randomly) to produce 50 replicates (see Table S1 for network and TOB summary statistics). Second, 1000 gene trees were simulated for each network under NMSC model with Independent Inheritance using PhyloCoalSimulations [14]. To vary the amount of ILS, we multiplied the branch lengths in the network by scaling factors: 2.0, 1.0, 0.5, and 0.25; this resulted in ILS levels of ~ 45%, 65%, 80%, 90%, respectively (Fig. S2). Higher ILS levels correspond to greater gene tree discordance, due to gene lineages failing to coalesce, and thus are more challenging model conditions. Third, sequences were simulated from each gene tree under the GTR+F+G model. Fourth, gene trees were estimated from sequences using IQ-TREE [30, 22]. The mean gene tree estimation error ranged from 40% to 70% (Figs. S3–S6). Lastly, gene trees were given as input to TOB-QMC and TINNiK. A detailed description of the simulation study, including software versions and commands, is provided in the Supplement.

#### Evaluation metrics

Methods were evaluated based on tree error and runtime. Tree error was computed as the false positive rate (FPR), false negative rate (FNR), and arithmetic mean of FNR and FPR. FNR is the number of branches in the true TOB not in the estimated TOB, divided by the total number of internal branches in the true TOB. FPR is the number of branches in the estimated TOB not in the true TOB, divided by the total number of internal branches in the estimated TOB; this definition should not be confused with the definition of FP in the proof of correctness for the 3f1a algorithm, in which we assume the initial tree is a binary TOB refinement, although this may fail to occur in practice.

#### Hyperparameter tuning

As previously mentioned, both TOB-QMC and TINNiK are based on hypothesis tests applied to subsets of four taxa, specifically the treelikeness test and the star test. Both methods were run using the default T3 option. The significance threshold *α* is used to reject the null hypothesis of treelikeness; low values are typically used to avoid false detection of non-treelike evolution due to gene tree estimation error or small numbers of gene trees. The significance threshold *β* is used to <u>not</u> reject the null hypothesis of starlikeness; higher values are used to avoid false detection of starlike evolutionary relationships. There is currently no formula for selecting *α, β* [1]. To ensure fair settings for both methods, we evaluated the impact of the *α, β* hyperparameters on the 50- and 100-taxon data sets using <u>true</u> gene trees for all combinations of *α* ∈ {10^−5^, 10^−6^, 10^−7^, 10^−8^, 10^−9^} and *β* ∈ {0.90, 0.95, 1.00} (note that these ranges are similar to those used in the original TINNiK study [1] and that setting *β* = 1 ensures the null of the star test is always rejected so branches are never contracted based on it). We observed that setting *β* = 0.95 and *α* = 10^−7^ gave good results for both methods, with error increasing for higher settings of *α* (Figs. S7–S10). We also explored the impact of TOB-QMC’s search algorithms by comparing the final TOB to the initial refinement tree without any edges contracted (Figs. S11–S12). All search algorithms reduced FPR compared to the initial refinement tree with little impact on FNR. TOB-QMC-3f1a had the highest FPR, whereas default and fast modes achieved similar FPR to TOB-QMC-exhaustive. Therefore, we benchmarked TOB-QMC-default and TOB-QMC-fast against TINNiK, both with *β* = 0.95 and *α* = 10^−7^.

### 2.3 Results on Simulated Data Sets with Estimated Gene Trees

#### Impact of ILS level and network level

We first evaluated TOB-QMC and TINNiK on data sets with estimated gene trees, varying the ILS and network levels. For 100 taxa, the TOB-QMC’s initial refinement tree had the best FNR but also the worst FPR of all methods (Fig. 2). FNR was generally low (less than ~5% on average) except for the highest ILS level where FNR increased but was still less than ~10% on average. TOB-QMC-fast and TOB-default had slightly higher FNR compared to the TOB-QMC refinement tree but much lower FPR, suggesting branch contraction based on the minimum *p*-value search was effective. The FPR varied, ranging from ~5% for the easiest model condition (level-1, branch scale factor 2) to ~18% for the hardest (level-2, branch scaling factor 0.25). The performance difference between TOB-QMC and TINNiK increased with model condition difficulty, with TOB-QMC outperforming TINNiK. These trends held on data sets with 50 and 200 taxa (Fig. S13–S14), except that TINNiK could not be run on the largest data sets with 200 taxa.

#### Impact of varying the number of taxa

Next we evaluated TOB-QMC and TINNiK for increasing numbers of taxa (Fig. 3). All methods were fast for 50 taxa. At 100 taxa, TINNiK took over 2 hours on average, while both TOB-QMC versions took less than 0.5 hours. At 200 taxa, TINNiK could not run within our wall clock limit (48 hours), but TOB-QMC-default and TOB-QMC fast took less than 2.5 hours and 1 hour on average, respectively. Tree error of TINNiK increased with the number of taxa, going from ~ 5% for 50 taxa to ~ 10% for 100 taxa, on average for level-2 networks (no branch length scaling). In contrast, the tree error of TOB-QMC was relatively stable (~ 5%). Similar trends were observed for the other model conditions (Figs. S15–S21).

**Fig. 3:**
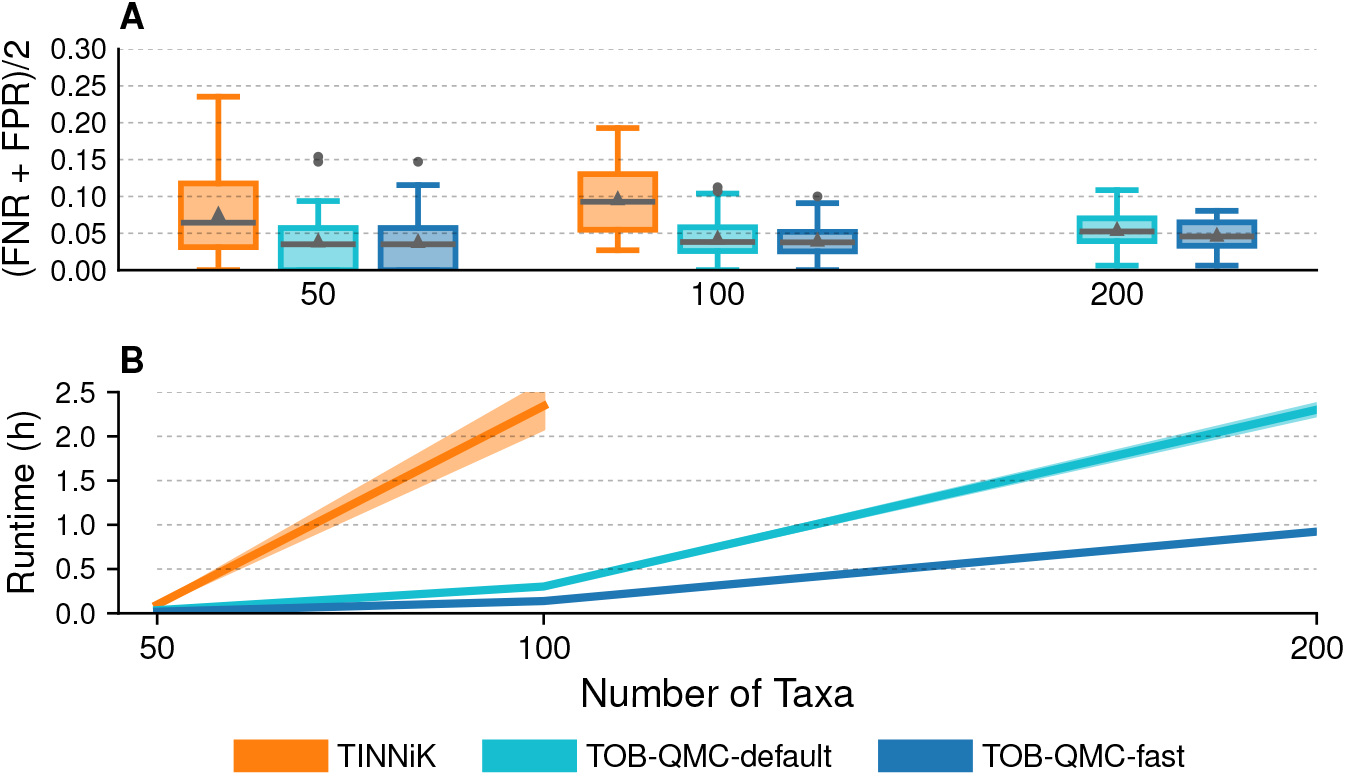
Tree error and runtime for TOB reconstruction methods for level-2 networks with branch scaling factor 1 (ILS level: 65%). Lines and shaded regions in subfigure (b) are means and standard deviations.

#### Impact of violating the quartet-nonanonamalous assumption

Lastly, we evaluated the prevalence of networks that violated the class 1 quartet-nonanomalous assumption and the impact of this model violation on TOB-QMC and TINNiK. To classify replicate networks, we calculated the expected qCFs for all subsets of four taxa, which was computationally intensive, limiting this analysis to networks with 50 and 100 taxa. We found that the fraction of anomalous networks (out of the 50 replicates per model condition) increased with ILS level and network level (Table S2). For 100 taxa, just 6% of networks were anomalous for the easiest model condition (level-1, branch scaling factor 2) but that increased to 50% for the hardest model condition (level −2, branch scaling factor 0.25). Sometimes the FPR and FNR of TOB-QMC and TINNiK were greater for anomalous networks compared to nonanomalous ones for the same model condition (e.g., 100 taxa, level-2, branch scaling factor 1), as might be expected; however, the opposite trend was observed for other model conditions (e.g., 100 taxa, level-1, branch scaling factor 0.25). Importantly, our main findings did not change: TOB-QMC was competitive with TINNiK in terms of tree accuracy, often outperforming it on challenging model conditions, which had the greatest proportion of anomalous networks (Fig. S22–S23).

### 2.4 Application to Phylogenomic Data Sets

Although TOB-QMC-3f1a had slightly higher FPR in simulations, it is fast and simple. FPR depends on setting a fixed *α, β* setting across the entire tree, which is not needed when exploring whether there are signals of non-treelike evolution. To evaluate the utility of TOB-QMC-3f1a for this purpose, we applied it to phylogenomic data sets for the bee subfamily *Nomiinae* (31 taxa, 852 gene trees) [9, 7], the butterfly subfamily *Heliconiinae* (63 taxa, 3393 gene trees) [11], and seed plants (96 taxa, 406 gene trees) [31, 23]. The bee data set was small so we also ran TINNiK and compared TOBs. Both TINNiK and TOB-QMC-3f1a were fast on the bee data set, taking less than 1 minute. The other phylogenomic data sets had larger numbers of taxa, but we were able to analyzed them on a laptop with TREE-QMC-3f1a in just a few minutes.

#### Nomiinae bees

On the bee data set, both TINNiK and TOB-QMC-3f1a returned the same binary TOB when *α* was set to 10^−5^ and 10^−7^, respectively. Increasing *α* to 10^−4^ and 10^−6^, respectively, resulted in a blob for the genus *Stictonomia* (Fig. S24; Table S3). A prior study by [9] reported conflicting species tree topologies on this genus, and gene flow was detected by [7] using several statistical tests for hybridization, including MSCquartets [33], which is similar to TINNiK, as well as HyDe [8] and Patterson’s D statistics [32]. As *α* increased, the TOBs returned by both methods became more unresolved. Notably, TINNiK needed to be re-run for each setting of *α, β*, whereas TOB–3f1a did not (because edges in the initial refinement tree were annotated with minimum *p*-values found as well as taxon names and qCFs for the minimizer subset). This saved time and enabled us to determine why edges were being contracted. For example, TOB-QMC-3f1a contracted the edge associated with *Stictonomia* because the treelikeness test (T3 option) applied to the three taxa in this genus plus *Nomiapis bispinosa* was *p* = 1.4 *×* 10^−7^. Although the associated qCFs of 80/69/21 (0.47/0.41/0.12) are indeed suggestive of gene flow (as the T3 test option evaluates whether two qCFs are equal and *lesser* than the third), there are just 170 gene trees to support this conclusion.

#### Heliconiinae butterflies

The TOB-QMC-3f1a analysis for butterflies had 14 edges with *α <* 10^−15^ and *β* ≥ 0.95; these edges were contracted to get the TOB shown in Fig. 4 (also see Fig. S25 and Table S4). All edges contracted were consistent with gene flow identified by the original study (Fig. 2B in [11]); see the Supplement for details. The original study [11] additionally detected gene flow among 3 species in *Wallacei* clade (branch 43) and in *Eueides* clade (e.g., branches 48 and 50). Branch 43 had *p* = 1.5 *×* 10^−8^ and qCFs of 2710/205/106 (0.90/0.07/0.04), and branch 48 had *p* = 0.0001 and qCFs of 1533/103/55 (0.91/0.06/0.03). These *p*-values are low, and thus suggestive of gene flow, but do not survive the high significance threshold of *α* = 10^−15^ used in the TOB-QMC analysis. Both branch 43 and 48 are characterized by the majority of the gene trees (~90%) supporting the same quartet topology. Of the remaining 10% of gene trees, one of the alternative quartet topologies is displayed 2*×* more than other. Thus, the overall impact on the genome is much low than the contracted branches. For branch 43, we exhaustively performed treelikeness tests (T3 option) on all four-taxon subsets around the edge (specifically its induced quadrapartition; see Methods). The four-taxon subsets included *H. burneyi, H. wallacei*, and *H. egeria*, plus every other taxa. All tests (60 in total) had similar qCFs and *p*-values (Table S5), so the result did not change with more aggressive search.

**Fig. 4:**
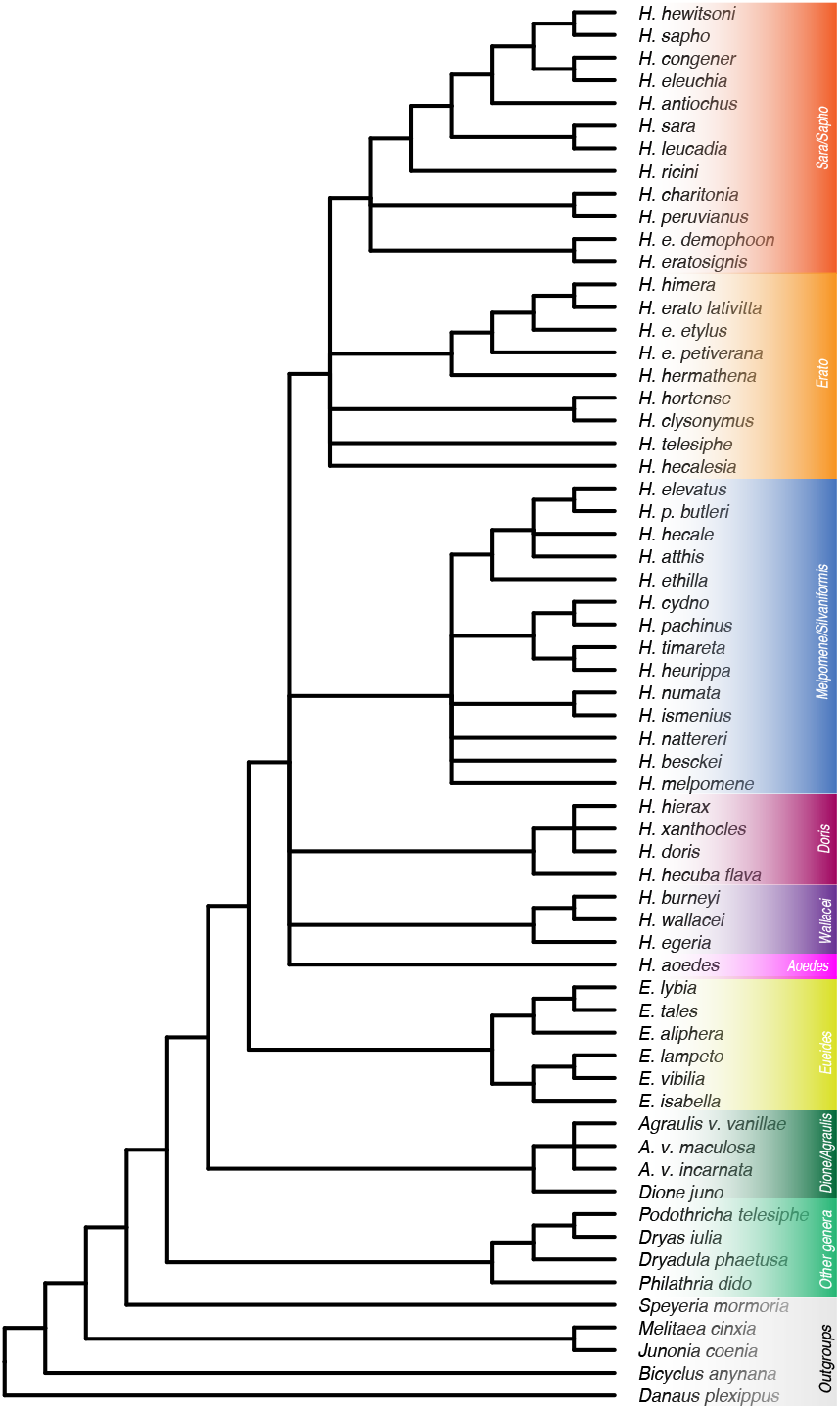
TOB for butterflies computed by TOB-QMC-3f1a setting *α* = 10^−15^ and *β* = 0.95.

#### Seed Plants

Lastly, we applied TOB-QMC to a seed plant data set, previously analyzed by the recently developed divide-and-conquer network method InPhyNet [23]. In the InPhyNet analysis, TINNiK was used to build a TOB for subset decomposition, with *α* = 0.001. The resulting TOB was fairly unresolved, as was the TOB produced in our TOB-QMC-3f1a analysis with *α* = 0.001 (25 branches were contracted; see Fig. S26 and Tables S6–S7). Notably, some branches towards the root had weaker evidence for contraction. For example, branches 6 and 7 had *p*-values less than 10^−5^ and qCFs of 220/27/3 (0.88/0.11/0.01) and 179/26/2 (0.87/0.13/0.01), respectively. Thus, the treelikeness test (T3 option) was based on just 250 and 207 gene trees, respectively, the majority of which (~87%) supported a single topology. Branch 25, in contrast, had a much lower *p*-value (*<* 10^−19^) and qCFs of 149/134/25 (0.48/0.44/0.08), so the test result was supported by a greater number of gene trees (308), the majority of which (~91%) were split between two quartet topologies. To summarize, gene flow around branch 25 had a greater overall impact on genomes compared to branches 6 and 7, although it is worth noting that errors in the identification of single copy orthologs can lead to systematic biases, which may appear like non-treelike evolution.

## 3 Discussion

In this work, we introduced a scalable framework for tree of blobs (TOBs) reconstruction under the network multi-species coalescent (NMSC) that first seeks a TOB refinement and then contracts edges, combining search heuristics with hypothesis testing. Although this approach is simple, we provide theoretical justification. First, we show that any optimal solution to WQC is a TOB refinement almost surely, as the number of gene trees goes to infinity, provided the underlying network is T-quartet-nonanomalous. Second, we show that our 3f1a search algorithm, which samples only a linear number of four-taxon subsets around each branch, correctly detects false positive (FP) edges in a TOB refinement,provided the underlying network is level-1. Together, these results yield a statistically consistent and fast approach for TOB reconstruction. Unlike TOB-QMC, TINNiK’s statistical guarantees are not restricted to level-1 networks; however, this comes with increased time complexity: *O*(*n*^4^*k* + *n*^5^) versus *O*(*n*^3^*k*). This trade-off is important for level-1 (semi-directed) network reconstruction with methods, such as InPhyNet or NANUQ+. Moreover, in our simulation study, TOB-QMC matched or exceeded TINNiK’s accuracy on level-2 networks, suggesting it could be a practical heuristic for higher-level networks.

Apart from network level, TOB-QMC and TINNiK (with T3 option) are subject to the same model assumptions as they were run with the same hypothesis tests and significance thresholds. Therefore, any improvements in accuracy likely stem from TOB-QMC’s two-stage design, which first aggregates signal across all four-taxon subsets and then applies targeted hypothesis tests. In contrast, TINNiK builds the tree largely by applying hypothesis tests to all four-taxon subsets individually. By not leveraging global signal across all subsets, TINNiK may be more sensitive to noisy inputs and model violations. It also is worth noting that TINNiK is deterministic, unlike TOB-QMC, whose output can change depending on the random seeding of the search algorithm (although we expect trends in performance to be unaffected because methods were run on 50 replicates per model condition).

A limitation of both TOB-QMC and TINNiK (T3 option) is the assumption that networks are quartet-nonanomalous. Ané *et al*. proved that networks with generic parameter settings are quartet-nonanomalous under the NMSC with Common Inheritance (*ρ* = 1) [5]; however, such strong positive results do not exist for more general models of inheritance, including Independent Inheritance (*ρ* = 0) used in our simulations. Although Ané *et al*. found anomalous networks with four taxa to be rare in simulations under a birth-death-hybridization model [5], many of the networks in our simulation were quartet-anomalous, especially model conditions with higher network levels and shorter branch lengths. Binning networks based on whether they were quartet-anomalous did not change the conclusions of our study. Perhaps surprisingly, the performance of both TOB-QMC and TINNiK sometimes improved for anomalous networks compared to nonanomalous networks for the same model condition, suggesting there may be correlations with other network properties that make TOB reconstruction easier. However, our study was not designed to evaluate the impact of quartet-anomalous networks on TOB reconstruction, and future work is needed to address this question as well as the extent to which such model violations are biologically realistic.

Another obstacle to TOB reconstruction, with either TOB-QMC or TINNiK, is the burden on users to set significance thresholds: *α, β*. Because TOB-QMC stores the minimum *p*-value associated with each edge, users can efficiently evaluate the impact of different *α* and *β* settings offline without recomputing the refinement tree. We recommend first examining the distribution of *p*-values to understand the strength of evidence across the tree, and then exploring a grid of *α, β* values to assess the stability of the inferred TOB across these settings. Selecting appropriate values of *α* and *β* could be challenging in practice, as the best choice may depend on data characteristics, such as ILS and gene tree estimation error levels, as well as on the search algorithm (e.g. 3f1a versus versus exhaustive search), and thus no single setting may be universally optimal. However, edges with low *p*-values provide strong evidence of non-treelike signal and will be contracted across a wide range of *α* values, whereas edges with *p*-values near the decision threshold will be sensitive to hyperparameter choices. Accordingly, edges that are contracted only under extreme parameter values (e.g., a very large *α*, leading to aggressive over-contraction) should be interpreted cautiously, whereas edges that are consistently contracted across a broad range of settings provide stronger evidence of a non-treelike signal. Future work should evaluate the effectiveness of hypothesis testing, both the tests developed by TINNiK as well as other approaches (e.g. [32]), and develop procedures for automatically setting significance thresholds to relieve the burden on users.

Alternatively, biologists may use TOB-QMC for exploring whether there are signals of non-treelike evolution in their data, which does not require fixed settings of *α, β*. Our analysis of phylogenomic data sets suggests that TOB-QMC-3f1a is fast and effective for this purpose. Moreover, the TOB-QMC framework is closely connected to how researchers analyze large data sets, first building a species tree with ASTRAL and then annotating it with gene flow or introgression events. This approach to network reconstruction can be positively misleading, as recently shown by [12]. Our study provides theoretical justification for such analyses, provided the results are interpreted carefully. Specifically, the ASTRAL tree should be interpreted as a TOB refinement and signals of non-treelike evolution around edges (i.e., low *p*-values) should be taken as an indication that those branches may not be trustworthy and that network reconstruction may be needed. During the time frame of our study, a new network reconstruction method, called Squirrel, was released [19]. Squirrel creates candidate TOBs, through edge contractions, which is similar to TOB-QMC, and then expands the blob vertices to produce a network (note that it contracts branches based on split support from quarnets estimated from DNA sequences). However, unlike TINNiK and TOB-QMC, Squirrel is not designed for coalescent-based species network reconstruction and has no statistical guarantees under the NMSC. However, other methods for level-1 network reconstruction targeting the NMSC model, such as NANUQ^+^ and InPhyNet, depend on TOB reconstruction. We therefore anticipate that TOB-QMC, or its algorithmic techniques, will greatly enable scalable level-1 network reconstruction under the NMSC; future work should explore this application.

## 4 Methods

To present TOB-QMC, we first review terminology, largely from [5], and prior results.

### 4.1 Terminology and Background

#### Phylogenetic networks

A *directed* phylogenetic network *N* ^+^ is a directed acyclic finite graph with the property that there exists a directed path from the root vertex (a unique vertex with in-degree 0) to all other vertices. Leaf vertices, which have out-degree 0, are bijectively labeled by a set *S* of species. There are two types of non-root, non-leaf vertices: *tree vertices*, which have in-degree 1, and *reticulation vertices*, which have in-degree *>* 1. We say that *N* ^+^ is *binary* if the root has degree 2, tree and reticulation vertices have degree 3, and leaves have degree 1. A tree edge is an arc whose child is a speciation or a leaf vertex; all other arcs are *reticulation edges*. A *wedge* path *P* from *v* to *v*^*′*^ in *N* ^+^ is a sequence of distinct vertices *v* = *u*_0_, *u*_1_ …, *u*_*k*−1_, *u*_*k*_ = *v*^*′*^ such that there exists some *u*_*i*_ that “roots” the path so (*u*_*i*_, *u*_*i*−1_), …, (*u*_1_, *v*) and (*u*_*i*_, *u*_*i*+1_), …, (*u*_*k*−1_, *v*^*′*^) are directed paths [20] (note that wedge paths are also called up-down paths in [5]). A directed network *N* ^+^ on species set *S* can be converted into a *semi-directed* network, denoted *N*^−^, by removing all edges and vertices above the lowest stable ancestor *s* of *S* (i.e., the lowest vertex that lies on all directed paths from the root to leaves in *S*; see [36], p.263), undirecting all tree edges, and suppressing *s* if it has degree 2 (Fig. 1 in [5]; also see [35, 6]). The wedge path terminology is used for semi-directed networks because there is a bijection between wedge paths in *N* ^+^ and paths in *N*^−^ that do not contain *v-structure* from traversing two reticulation edges incident to the same target. A **subnetwork** of *N*^−^ on *Y* ⊆ *S* is the subgraph induced by the union of vertices and edges on all wedge paths between all pairs of taxa in *Y* (Fig. 1). The **restriction** of *N*^−^ to *Y*, denoted *N*^−^|_*Y*_, is formed by computing the subnetwork on *Y* and then repeatedly suppressing degree 2 vertices, parallel arcs, and degree 2 blobs [20] (see Fig. S1 as an example). A phylogenetic tree is simply a phylogenetic network without any reticulation vertices. A tree is **displayed** by *N*^−^ if it can be obtained from *N*^−^ by deleting all but one of the reticulation edges with the same target vertex and then deleting edges and vertices not contained in any wedge path between any two leaves [5] (note that it is often convenient to suppress degree 2 vertices).

A bridge is an edge whose deletion increases the number of connected components of the graph. A *blob* in a network is a maximal bridgeless subgraph when treating all edges as undirected. A blob is trivial if it consists of a single vertex. The level of a blob is the number of reticulation edges in it minus the number of reticulation vertices in it (note that a network is level-*k* if *k* is the maximal level of its blobs, or equivalently, it can be transformed into a tree by removing at most *k* edges from each of its blobs). The degree of a blob is the number of bridges adjacent to it (an *m*-blob is a blob with degree *m*). Edges in blobs are **blob edges**, and all other edges (i.e., bridges) are **cut edges**. The **tree of blobs (TOB)** [15] can be computed from a network by (1) un-directing all edges, (2) contracting all blob edges, and then (3) suppressing all vertices with degree two (Fig. 1A–B). The contraction step results in each blob being collapsed into a single vertex, which may have degree greater than 3, referred to as a polytomy. A tree *T* refines the TOB if the TOB can be obtained by contracting edges in *T* [38]. Any tree displayed by a network refines the TOB (as all cut edges are preserved); however, a TOB refinement need not be a display tree of the network (Fig. 1C).

The deletion of a vertex *v* with degree *m* in a phylogenetic tree produces *m* connected subgraphs with leaves labeled by taxon subsets *P*_1_, *P*_2_, …, *P*_*m*_; thus, *v* induces an *m***-partition** of the species set *S*, denoted *Part*(*v*) = *P*_1_|*P*_2_| … |*P*_*m*_; the same holds for the deletion of an *m*-blob in a phylogenetic network. Likewise, the deletion of edge *e* in a tree (or the deletion of a cut edge *e* in a network) induces a **bipartition**, denoted *Bip*(*e*) = *A*|*B*. A bipartition is trivial if either |*A*| = 1 or |*B*| = 1; otherwise, it is non-trivial. We say that four taxa are **around** an edge *e* in a tree *T* (or around a cut edge in a network) if exactly two taxa are drawn from each block of the bipartition induced by *e*. If both endpoints of *e* have degree 3, deleting *e* and its endpoints also induces a **quadrapartition** *Quad*(*e*) = *A*_1_|*A*_2_|*B*_1_|*B*_2_, where *A*_1_ ∪ *A*_2_ = *A* and *B*_1_ ∪ *B*_2_ = *B* (or vice versa). A **quartet** *q* is an unrooted tree on four species. There are three possible quartet topologies denoted by their non-trivial bipartition: *a, b*|*c, d* and *a, c*|*b, d* and *a, d*|*b, c*, where *S* = {*a, b, c, d*}. We say that quartet *q* on *Y* ⊆ *S* is induced by an unrooted tree *T* on *S* if *T*|_*Y*_ is isomorphic to *q*.

A semi-directed, binary network on four taxa contains a 4-blob or not. Not having a 4-blob, which we refer to as being *treelike*, is equivalent to having a cut edge that induces a non-trivial bipartition or having a TOB that is a binary quartet tree [1, 20]. Conversely, having a 4-blob, which we refer to as being *non-treelike*, is equivalent to having cut edges that only induce trivial bipartitions or having a TOB that is an unresolved star tree. There is also a connection to the number of display trees.

##### Lemma 1

**(Lemma 1 in [5])**. *A semi-directed, binary network on four taxa displays exactly one binary quartet tree if, and only if, it does not contain a 4-blob*.

For more than four species, the above results hold after restricting the network to four species; however, the relationship among the four species can change when looking at the TOB for the subnetwork versus the larger network. Take Figure 1 as an example. Restricting *N* ^+^ to *A, B, F, G* and then computing the TOB of the subnetwork yields the quartet tree *A, B*|*F, G*, so an oracle would say their relationship is treelike; however, computing the TOB for *N* ^+^ and then restricting it to *A, B, F, G* yields an unresolved star tree, called a *B-quartet* by [1]. Fortunately, there is a nice relationship between cut edges of a network and its four-taxon subnetworks, specifically:

##### Lemma 2

**(Forward Direction of Theorem 5.1 in [20])**. *Let N*^−^ *be a semi-directed, binary network on species set S. If A*|*B is a bipartition induced by a cut edge of N*^−^, *then either A*|*B is trivial or for all pairwise distinct elements a*_1_, *a*_2_ ∈ *A and b*_1_, *b*_2_ ∈ *B, a*_1_, *a*_2_|*b*_1_, *b*_2_ *is induced by a cut edge of* 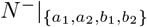.

This result follows from the definition of cut edge and the construction of subnetworks. It is noteworthy that the converse also holds (Theorem 5.1 in [20]), although it is unnecessary for our results.

#### Network Multispecies Coalescent (NMSC)

The NMSC is parameterized by a species network 𝒩 ^+^ = (*N* ^+^, *Θ*), where *N* ^+^ is a directed phylogenetic network and *Θ* is a set of numerical parameters: branch lengths in coalescent units, inheritance proportions, and the inheritance correlation *ρ* [14]. A gene genealogy can be generated under the NMSC by sampling one gene per species and tracing ancestry backward in time until all gene lineages coalesce (i.e., trace back to a common ancestor); see the Supplement for details. We let ℙ _*𝒩*_+ (*T*) denote the probability of an unrooted gene genealogy *T* on species set *Y* ⊆ *S* under the NMSC given 𝒩 ^+^. 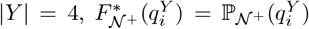 for *i* = {1, 2, 3} are the <u>expected</u> **quartet concordance factors (qCFs)** where *i* indexes the three possible quartets on *Y*. Expected qCFs are fully determined by the semi-directed version of 𝒩 ^+^ (Proposition 3 in [5]); thus, we use 𝒩^−^ going forward. The <u>observed</u> qCFs are computed by counting the number of gene trees (out of *k*) that induce 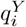, denoted 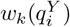, and then dividing by *k* (or normalizing so that the qCFs for *Y* sum to 1 if gene trees are missing taxa).

We make standard assumptions about networks (i.e., conditions 1-3 in TOB-QMC Method Overview). Our first result further assumes networks are **T-quartet-nonanamolous**.

##### Definition 1

**(Quartet-nonanomalous).** *Let* 𝒩^−^ = (*N*^−^, *Θ*) *be a semi-directed species network on species set S that satisfies standard assumptions. We say that* 𝒩^−^ *is* quartet-nonanomalous *if for every subset Y* ⊆ *S of four taxa*,

1. *N*^−^|_*Y*_ *displays more than one quartet tree <u>or</u>*
2. *N*^−^|_*Y*_ *displays one quartet tree* 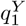 *and the following inequalities hold:* 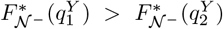 *and* 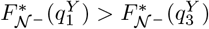 *where* 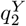 *and* 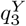 *denote the other two quartets on Y*.

*The definition above only puts restrictions on subnetworks that display one quartet tree, called class 1 (sometimes restrictions are also put on subnetworks that display two quartet trees, called class 2; see Definition 5 in [5]). Our first result only requires the inequalities above to hold on subsets Y for which the TOB of N*^−^ *restricted to Y is a quartet tree, called a* T-quartet *by [1]; thus, we refer to this property as* T-quartet-nonanomalous. *This assumption is no more restrictive than the one made by TINNiK (T3 option)*.

Although one might think that quartet-based species tree methods, such as ASTRAL, would be guaranteed to produce a display tree of the network if it is quartet-nonanonmalous, Dinh and Baños recently showed this is not the case [12].

#### Weighted Quartet Consensus (WQC)

The popular species tree method ASTRAL-III [43] is a constrained dynamic programming for the NP-hard WQC problem [24]. The input to WQC is *k* unrooted gene trees, whose leaves are labeled by a set *S* of *n* taxa. The output is an unrooted binary tree *T* on *S* that maximizes the quartet score: 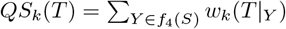, where *f*_4_(*S*) denotes all four-taxon subsets of *S*, that is *f*_4_(*S*) = {{*a, b, c, d*} : *a, b, c, d* ∈ *S, a*≠ *b c*≠ *d*}. It is worth noting that ASTRAL does not explicitly compute quartet weights.

### 4.2 WQC is a TOB refinement almost surely as the number of gene trees goes to infinity

Our framework for TOB reconstruction begins by seeking a refinement of the TOB with a quartet-based species tree method. To motivate this approach, we show that any optimal solution to WQC is a TOB refinement almost surely, as the number of gene trees goes to infinity (Theorem 1); our proof relies heavily on the next result.

#### Lemma 3.

*Let* 𝒩^−^ = (*N*^−^, *Θ*) *be a species network on species set S that satisfies standard assumptions and is T-quartet-nonanomalous. Assume that the error in all qCFs is zero (i*.*e*., *they equal their expected values under the NMSC given* 𝒩^−^*). Then, there exists a binary TOB refinement* 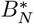 *with higher quartet score than any binary tree T on S that does <u>not</u> refine the TOB. Note that we assume the TOB for N*^−^ *is not an unresolved star tree; otherwise, all binary trees on S refine it so the result is trivial*.

*Proof*. Observe that restricting the TOB *T*_*N*_ to any 4-tuple of species *Y* ∈ *f*_4_(*S*), denoted *T*_*N*_ |_*Y*_, yields either a binary quartet tree or an unresolved star tree (note that these are called T- and B-quartets by [4]). Let 𝒮_*tree*_(*T*_𝒩_) and 𝒮_*star*_(*T*_𝒩_) denote the four-taxon subsets in *f*_4_(*S*) partitioned based on those two cases, respectively. Now the quartet score of an unrooted binary tree *t* can be written as

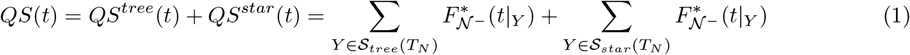

where *QS*^*tree*^(·) is the “tree part” of quartet score and *QS*^*star*^(·) is the “star part” of the quartet score (see Fig. 5 for an example). Note that we omit the subscript *k* from *QS* to indicate that it is computed from expected qCFs rather than observed qCFs by our assumption.

**Fig. 5:**
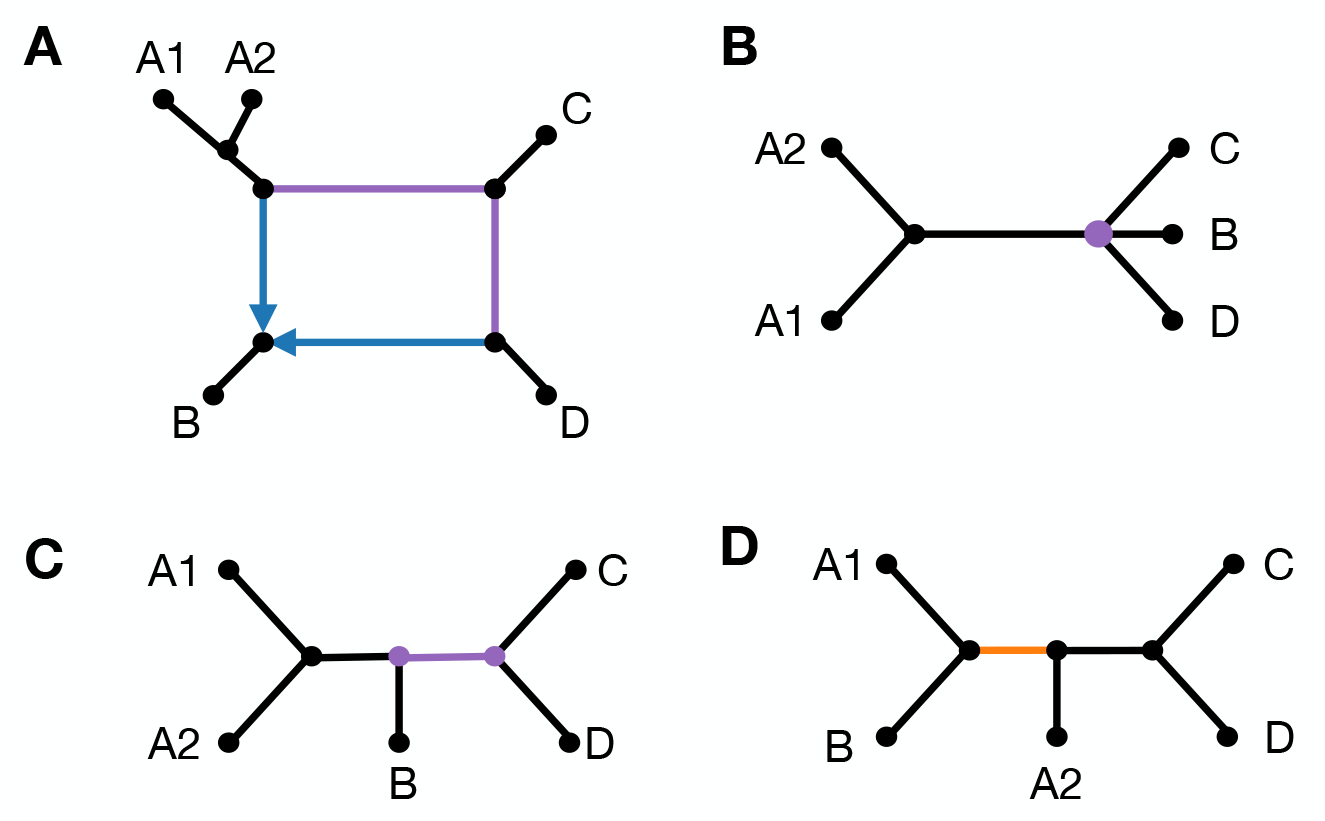
**A)** A 5-taxon semi-directed network *N*^−^. **B)** TOB for *N*^−^ has one polytomy *v* that induces partition *Part*(*v*) = {*P*_1_ = {*A*_1_, *A*_2_}, *P*_2_ = {*B*}, *P*_3_ = {*C*}, *P*_4_ = {*D*}}. Thus, the “tree part” of the quartet score *QS*^*tree*^(·) is computed from 4-tuples: *A*_1_, *A*_2_, *B, C* and *A*_1_, *A*_2_, *B, D* and *A*_1_, *A*_2_, *C, D*. The “star part” of the quartet score *QS*^*star*^(·) is computed from 4-tuples: *A*_1_, *B, C, D* and *A*_2_, *B, C, D* (note that these two subsets have the same expected qCFs because *A*_1_ and *A*_2_ are exchangeable). **C)** A TOB refinement for *N*. TOB refinements maximize the tree part of the quartet score. Assume that this TOB refinement also maximizes the star part of the quartet score, i.e., 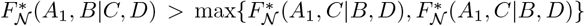. **D)** A TOB non-refinement that also maximizes that star part of the quartet score because it induces the same quartets on *A*_1_, *B, C, D* and *A*_2_, *B, C, D* as subfigure (c). However, TOB non-refinements do not maximize the tree part of the quartet score.

#### Tree part of quartet score

We first consider maximizing the tree part of the quartet score. Observe that for any binary tree *T* on *S* that is <u>not</u> a refinement of *T*_*N*_,

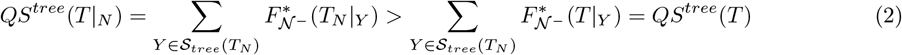

by our assumption that 𝒩^−^ is T-quartet-nonanomalous. Thus, the tree part of the quartet score is uniquely maximized by *T*_*N*_.

#### Star part of quartet score

We now consider maximizing the star part of the quartet score, partitioning 𝒮_*star*_(*T*_*N*_) based on the observation that for any *Y* = {*a, b, c, d*} ∈ 𝒮_*star*_(*T*_*N*_), there exists a unique vertex *v* in *T*_*N*_ with degree *m >* 3 whose deletion induces an *m*-partition *Part*(*v*) = *P*_1_|*P*_2_| … |*P*_*m*_ such that *a* ∈ *P*_*i*_, *b* ∈ *P*_*j*_, *c* ∈ *P*_*s*_, *d* ∈ *P*_*t*_ with *P*_*i*_≠ *P*_*j*_≠ *P*_*s*_≠ *P*_*t*_. This gives

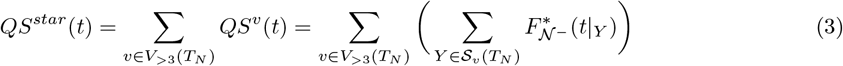

where *V*_*>*3_(*T*_*N*_) is the set of vertices in *T*_*N*_ with degree greater than 3 and 𝒮_*v*_(*T*_*N*_) is the set of all four-taxon subsets “around” vertex *v* (i.e., all ways of drawing four taxa from distinct blocks of the partition induced by *v*). More precisely, let *f*_4_(*Part*(*v*)) be the set made from all ways of selecting four distinct blocks from *Part*(*v*), that is *f*_4_(*Part*(*v*)) = {*P*_*i*_, *P*_*j*_, *P*_*k*_, *P*_*l*_ : *P*_*i*_, *P*_*j*_, *P*_*k*_, *P*_*l*_ ∈ *Part*(*v*), *P*_*i*_,≠ *P*_*j*_≠ *P*_*k*_≠ *P*_*l*_}, and let *f*_*cover*_({*P*_*i*_, *P*_*j*_, *P*_*k*_, *P*_*l*_}) be the set made from all ways of selecting one taxa from each of the four input blocks, that is, *f*_*cover*_({*P*_*i*_, *P*_*j*_, *P*_*k*_, *P*_*l*_}) = {{*a, b, c, d*} : *a* ∈ *P*_*i*_, *b* ∈ *P*_*j*_, *c* ∈ *P*_*k*_, *d* ∈ *P*_*l*_}. Then, 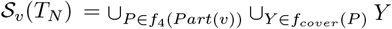.

We now focus on optimizing each “blob *v* part” of the quartet score *QS*^*v*^(·), associated with a single vertex *v* ∈ *V*_*>*3_(*T*_*N*_) created by contracting a blob in *N*^−^ into a polytomy. We ***claim*** that there exists a tree 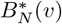 that (1) maximizes *QS*^*v*^(·) and (2) does not break apart the blocks of *Part*(*v*), meaning that each bipartition in the set {*P*_*i*_|*S* ∈ *P*_*i*_ : *P*_*i*_ ∈ *Part*(*v*)} is induced by an edge of 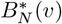. Our proof of this claim is based on the following observation: any 4-tuple of species covering {*P*_*i*_, *P*_*j*_, *P*_*k*_, *P*_*l*_} have the same expected qCFs because any lineage sampled from a species in *P*_*i*_ enters the blob in 𝒩^−^, associated with *v* in *T*_*N*_, from the same cut edge (and similarly for *P*_*j*_, *P*_*k*_, *P*_*l*_), and only after the four sampled lineages enter this blob can they coalesce. Thus, the expected qCFs are fully dictated by the blob and do not depend on which particular taxon is selected from a given block (i.e., taxa in the same block are exchangeable in this setting); see [1, 5].

We are now ready to prove our claim. For the sake of contradiction, suppose that all trees that maximize *QS*^*v*^(·) violate property (2). Let *T*^∗^ be one such tree. Then, there exists at least one block *P*_*j*_ in *Part*(*v*) such that no edge in *T*^∗^ induces bipartition *P*_*j*_|*S* ∈ *P*_*j*_. Let *QS*^*v*(*x*)^(·) denote the part of the quartet score from blob vertex *v* that is associated with taxon *x*, specifically:

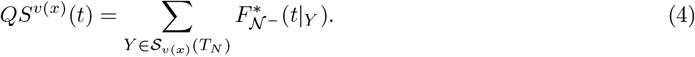

where 𝒮_*v*(*x*)_(*T*_*N*_) = {*Y* ∈ 𝒮_*v*_(*T*_*N*_) : *Y* ∩ {*x*} = ∅}. Next, select taxon *a* ∈ *P*_*j*_ such that *QS*^*v*(*a*)^(*T*^∗^) ≥ *QS*^*v*(*b*)^(*T*^∗^) for all *b* ∈ *P*_*j*_ ∈ \{*a*}. For each selection of *b*, there are two possible cases: *QS*^*v*(*a*)^(*T*^∗^) = *QS*^*v*(*b*)^(*T*^∗^) <u>or</u> *QS*^*v*(*a*)^(*T*^∗^) *> QS*^*v*(*b*)^(*T*^∗^). Now, detach *b* and reattach it so that it is sister to *a*. The resulting tree has the same blob *v* score as *T*^∗^ (i.e., *QS*^*v*^(*T*^∗^)−*QS*^*v*(*b*)^(*T*^∗^)+*QS*^*v*(*a*)^(*T*^∗^) = *QS*^*v*^(*T*^∗^)) because (i) there is a bijection between four-taxon subsets in 𝒮_*v*(*b*)_(*T*_*N*_) and four-taxon subsets in 𝒮_*v*(*a*)_(*T*_*N*_) and (ii) for any four-taxon subset {*b, x, y, z*} ∈ 𝒮_*v*(*b*)_(*T*_*N*_), the matched four-taxon subset {*a, x, y, z*} ∈ 𝒮_*v*(*a*)_(*T*_*N*_) has the same expected qCFs (i.e., 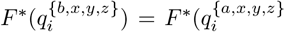 for all *i* ∈ {1, 2, 3}); this follows from the construction of 𝒮_*v*_(*T*_*N*_) and taxon exchangeability, as previously discussed. For the second case, the same actions are taken, but now the resulting tree has a blob *v* score greater than *T*^∗^ (i.e., *QS*^*v*^(*T*^∗^) − *QS*^*v*(*b*)^(*T*^∗^) + *QS*^*v*(*a*)^(*T*^∗^) *> QS*^*v*^(*T*^∗^)). Repeating for all *b* ∈ *P*_*j*_ |{*a*} and all *P*_*j*_ ∈ *Part*(*v*) yields a tree that has blob *v* score greater than or equal to *T*^∗^ and satisfies property (2), which is a contradiction. Thus, 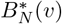 exists, although it is not necessarily a unique maximizer of *QS*^*v*^(·), and moreover, trees violating property (2) can maximize *QS*^*v*^(·).

We now make some observations about properties of 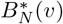. First, the bipartitions preserved in property (2) are induced by the edges of *T*_*N*_ that are incident to *v*. Second, 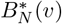 must be binary after removing all but one taxon from each block of *Part*(*v*). This follows by contradiction as any refinement of this modified tree increases *QS*^*v*^(·) (because all expected qCFs are positive under standard assumptions) and the score increase would be amplified after reattaching the removed taxa. Third, we can take a version of 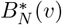 going forward that contracts all edges that induce bipartitions of the form *Z*|*W* where *Z* ⊂ *P*_*i*_ and *W* ∩ *P*_*i*_≠ ∅ (or vice versa). Any four-taxon subset around such an edge must draw two taxa from *P*_*i*_ and thus cannot be in 𝒮_*v*_(*T*_*N*_) by construction, so these edge contractions do not impact the score *QS*^*v*^(·). To summarize, 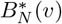 is a binary refinement of vertex *v* in *T*_*N*_ that maximizes *QS*^*v*^(·).

#### Putting tree and star parts together

By bipartition compatibility, there exists a compatibility supertree 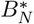 for 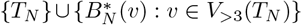 (Chapters 3, 7 in [37]). 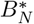 is a binary refinement of the TOB that maximizes each of the blob *v* parts of the quartet score, by our construction of each 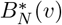, as well as the tree part of the quartet score. It follows that 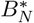 has a greater quartet score than any binary tree *T* that does <u>not</u> refine the TOB because even if the two trees achieve the same score on the star part, 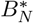 achieves a greater score on the tree part, which is why we assume the TOB is not an unresolved star tree.

Following Lemma 3, we show that 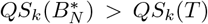 if the absolute error in qCFs is bounded by δ^∗^ = *π/*2|*f*_4_(*S*)|, where the *π* is minimum absolute difference between the qCFs of T-quartets (Lemma 4 in the Supplement). By the strong law of large numbers, observed qCFs converge almost surely to their expectation simultaneously, as *k* → ∞ (this assumption is standard in the phylgoenetic literature, e.g. [25]). This means that for any *ϵ >* 0 and any δ *>* 0, there exists an integer *K >* 0 such that with probability at least 1 − *ϵ*, the inequality 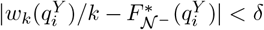 holds for all *i* ∈ {1, 2, 3}, all *Y* ∈ *f*_4_(*S*) and all *k* ≥ *K*.

This gives us our result:

##### Theorem 1

*Let* 𝒩^−^ = (*N*^−^, *Θ*) *be a species network on S that satisfies standard assumptions and is T-quartet-nonanomalous. Let G*_1_, *G*_2_, …, *G*_*k*_ *be a sequence of k independent and identically distributed unrooted gene trees on S sampled from the distribution induced by NMSC given* 𝒩^−^. *Then, for any binary tree T that does <u>not</u> refine the TOB and any ϵ >* 0, *there exists a binary TOB refinement* 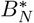 *and an integer K >* 0 *such that* 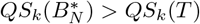 *for all k* ≥ *K with probability at least* 1 − *ϵ. Put simply, any optimal solution to WQC is a refinement of the TOB for N*^−^ *almost surely, as k* → ∞. *Note that we assume the TOB for N*^−^ *is not an unresolved star tree; otherwise all binary trees on S refine the TOB so the result is trivial*.

A corollary of Theorem 1 is that ASTRAL (version 3 or earlier) returns a TOB refinement almost surely, as the number of gene trees goes to infinity. This is because ASTRAL solves WQC within a constrained search space, defined by the bipartitions induced by the input gene trees, and all bipartitions have probability strictly greater than zero under the NMSC and thus will appear as the number of gene trees goes to infinity, as noted by [26]. This motivates the use of ASTRAL, and other WQC heuristics, in seeking a TOB refinement; our implementation uses TREE-QMC.

### 4.3 False positives in a TOB refinement can be identified with a linear number of tests for level-1 networks

To obtain a consistent estimator of the TOB, we need to correctly detect all edges in the TOB refinement that are not in the TOB, called false positives (FPs). Our approach is based on applying an oracle for treelikeness to four-taxon subsets sampled around each edge. The following lemma is useful for proving our second result.

#### Lemma 4.

*Let N*^−^ *be a semi-directed, binary, level-1 species network on taxon set S. Let T*_*N*_ *be the TOB of N*^−^, *and let B*_*N*_ *be a binary refinement of T*_*N*_. *Let e be an arbitrary edge in B*_*N*_ *that is <u>not</u> in T*_*N*_, *so it refines a vertex v*_*B*_ *in T*_*N*_ *created by contracting the edges of some non-trivial blob* ℬ *in N*^−^. *Then, for any four-taxon subset Y around e, N*^−^|_*Y*_ *contains a 4-blob if and only if Y satisfies two conditions:*

1. *Each taxon in Y is selected from a distinct block of the m-partition induced by v*_ℬ_, *denoted Part*(*v*_ℬ_).
2. *Exactly one taxon in Y is a descendant of the unique reticulation vertex v*_*h*_ *in* ℬ.

*Proof*. Let *Z*|*Y* be the bipartition induced by *e*, and for *Y* = {*a, b, c, d*}, let *a, b* ∈ *Z* and *c, d* ∈ *W* by our assumption that *Y* is around *e*. Because edge *e* in *B*_*N*_ is a refinement of vertex *v*_ℬ_ in *T*_*N*_, the blocks of *Part*(*v*_*B*_) = *P*_1_|*P*_2_| … |*P*_*m*_ are distributed across *Z*|*W* (i.e., each *P*_*i*_ is contained in either *Z* or *W*). Any taxa drawn from the same block attach to ℬ via the same cut edge, and conversely, any taxa drawn from distinct blocks attach to ℬ via distinct cut edges. Let *B*(*x*) denote the vertex in ℬ incident to the cut edge whose deletion separates taxon *x* from the subgraph containing ℬ.

⇐ Assume that *Y* meets the two conditions above. By condition (1), *B*(*a*), *B*(*b*), *B*(*c*), *B*(*d*) are distinct. By condition (2), assume w.l.o.g that *d* is the only taxon in *Y* that descends from the unique reticulation vertex *v*_*h*_ in ℬ. Then, because ℬ is level-1, there are exactly two wedge paths from *v*_*h*_ = *B*(*d*) to *B*(*a*), and the union of their vertices and edges is ℬ. Adding the vertices and edges on the wedge paths from *x* to *B*(*x*) for all *x* ∈ *Y* and then suppressing all degree 2 vertices, parallel edges, and degree 2 blobs yields *N*^−^|_*Y*_. Thus, these conditions are sufficient for *N*^−^|_*Y*_ to contain a 4-blob.

⇒ Now, we just need to show that these conditions are necessary for *N*^−^|_*Y*_ to contain a 4-blob. Based on our assumptions, either *a, b, c, d* are drawn from distinct blocks of *Part*(*v*_ℬ_) or else at least one pair *a, b* or *c, d* are drawn from the same block. The latter results in at least one cut edge in *N*^−^ whose deletion splits *a, b* from *c, d*; thus, *N*^−^|_*Y*_ contains a cut edge inducing *a, b*|*c, d* by Lemma 2. Thus, *a, b, c, d* must come from distinct blocks. Now suppose that none are descendants of the unique hybrid vertex *v*_*h*_ in ℬ. Then, because ℬ is level-1, there is only one wedge path between each pair of distinct vertices in {*B*(*a*), *B*(*b*), *B*(*c*), *B*(*d*)} (the only other path crosses *p*_1_, *v*_*h*_, *p*_2_ where (*p*_1_, *v*_*h*_) and (*p*_2_, *v*_*h*_) are the two reticulation edges in ℬ so it is not wedge). As no wedge paths include *v*_*h*_, there is a single path connecting *B*(*a*), *B*(*b*), *B*(*c*), *B*(*d*) in the formation of *N*^−^|_*Y*_, and their order defines the non-trivial bipartition induced by a cut edge in *N*^−^|_*Y*_; thus, it cannot contain a 4-blob. Thus, both conditions are necessary for *N*^−^|_*Y*_ to contain a 4-blob.

#### Corollary 1.

*Let N*^−^ *be a semi-directed, binary, level-1 species network on taxon set S. Let T*_*N*_ *be the TOB of N*^−^, *and let B*_*N*_ *be a binary refinement of T*_*N*_. *A correct labeling of branches in B*_*N*_ *is obtained by applying an oracle* 𝒪 *for treelikeness to all four-taxon subsets around each edge e and labeling e as FP if* 𝒪 *returns non-treelike for one or more subsets (otherwise e is labeled as TP)*.

*Proof*. By Lemma 2, all TPs will never get incorrectly labeled as FPs by this approach. Therefore, to conclude the proof, it is only left to show that if *e* is an FP, there exists at least one four-taxon subset *Y* ^∗^ around *e* such that *N*^−^|_*Y*_ ∗ contains a 4-blob. By the setting, any FP edge *e* in *B*_*N*_ refines a vertex *v* _ℬ_ in *T*_*N*_ created by contracting the edges of some non-trivial blob ℬ in *N*^−^, so the blocks of *Part*(*v*_ℬ_) are distributed across the bipartition induced by *e*. Observe that (1) ℬ has a unique hybrid vertex *v*_*h*_ by our assumption that it is level-1, so the deletion of the cut edge incident to *v*_*h*_ corresponds to the block of *Part*(*v*) that contains all descendants of *v*_*h*_, (2) |*Part*(*v*_ℬ_)| ≥ 4 (otherwise there is no way to refine vertex *v*_ℬ_), and (3) each side of the bipartition induced by *e* contains at least two blocks of *Part*(*v*_ℬ_) (otherwise ℬ_*N*_ is not binary). Thus, there is at least one four-taxon subset *Y* ^∗^ around *e* that meets the conditions of Lemma 4, giving us our result.

Corollary 1 gives a brute force approach for FP detection, which is Ω(*n*^4^) per edge and thus Ω(*n*^5^) in total, which is no better than TINNiK. Our goal is to design efficient algorithms that are guaranteed to find at least one four-taxon subset that meets the conditions of Lemma 4. This leads us to define a disjoint cover.

#### Definition 2

**(Disjoint cover)**. *Let T be a phylogenetic tree on species set S, and let Q*(*e*) *be a set of four-taxon subsets around e in T. We say that Q*(*e*) *is a* disjoint cover *if two conditions are satisfied: (1) every four-taxon subset in Q*(*e*) *draws one taxon from each block of the quadrapartition induced by e, and (2) every taxon in the complete species set S is included in at least one four-taxon subset in Q*(*e*).

#### Theorem 2.

*Let N*^−^ *be a semi-directed, binary, level-1 network on S, let T*_*N*_ *be the TOB of N*^−^, *and let B*_*N*_ *be a refinement of T*_*N*_. *Let* 𝒜 *be an algorithm that applies an oracle* 𝒪 *for treelikeness to a disjoint cover of four-taxon subsets around each e in B*_*N*_, *labeling e as FP if* 𝒪 *returns non-treelike for at least one subset tested (otherwise e is labeled as TP). Then, applying* 𝒜 *to* ℬ_*N*_ *followed by contracting all branches labeled FP returns T*_*N*_.

*Proof*. Following the proof of Corollary 1, it suffices to show that if *e* is an FP edge, then there is at least one subset *Y* ^∗^ in the disjoint cover that satisfies the conditions of Lemma 4. By the setting, the blocks of *Part*(*v* _ℬ_) = *P*_1_|*P*_2_| … |*P*_*m*_ are distributed across the quadrapartition of *e* (i.e., each *P*_*i*_ is contained in a single block of the quadrapartition of *e*). Condition 1 of the disjoint cover requires one taxon to be drawn from each block of the quadrapartition induced by *e*, so in turn, one taxon is drawn from a distinct block of *Part*(*v*_ℬ_); thus, all subsets in *Q*(*e*) satisfy condition 1 of Lemma 4. Condition 2 of the disjoint cover requires every taxon in *S* to be included in at least one of the subset in *Q*(*e*), so at least one subset in *Q*(*e*) contains a taxon descending from the unique reticulation vertex in ℬ; such a subset also satisfies condition 2 of Lemma 4.

Theorem 2 leads us to propose an algorithm that selects one taxon from each block of the quadrapartition, fixes the selection for three of the blocks, and then tests all ways of selecting a taxon from the fourth block, and then iterating so that the “alter” step is performed for every block of the quadrapartition. We refer to this quadraparition search algorithm as **3-fix, 1-alter (3f1a)** (see Algorithm 1 in the Supplement). It is easy to see that 3f1a tests a disjoint cover of four-taxon subsets and thus has a correctness guarantee when the underlying network is level-1. In practice, we do not have an oracle and instead apply TINNiK’s hypothesis tests to qCFs computed from gene trees, with fixed significance thresholds *α, β*. In this setting, the consistency our approach for level-1 networks follows from combining Theorem 1, Theorem 2, and Theorem 4 in [1], which states that there exists a sequence *α*_*k*_ such that TINNiK’s hypothesis testing procedure applied to *k* gene trees returns the correct answer for all four-taxon subsets of *S* with probability going to 1, as the number *k* of gene trees goes to infinity (note that any setting of *β >* 0 is allowed and that *α*_*k*_ → 0).

Unlike TINNiK, which tests all four-taxon subsets, our 3f1a algorithm tests just *O*(*n*) 4-taxon subsets around each edge, so *O*(*n*^2^) in total. The qCFs for any four-taxon subset can be computed from gene trees on the fly in *O*(*k*) time using the well-known binary lifting algorithm to retrieve the LCA of any pair of leaves in a gene tree in *O*(1) time. This requires a precomputation step with time and storage complexity of *O*(*n* log(*n*)*k*), meaning the total time complexity of applying the 3f1a algorithm to every branch is *O*(*n*^2^*k* + *n* log(*n*)*k*).

We found that the 3f1a search algorithm was effective for biological data, where it was not necessary to set a fixed *α* value, but it was less effective for simulated data, even for level-1 networks (Figs. S13–S14 in the Supplement), potentially because setting a single *α* value that is effective across the tree presents challenges. Notably, there can be more than one four-taxon subset around an edge whose relationship is non-treelike and some may yield lower *p*-values than others due to noise, model violations, or differences in their expected qCFs if they draw taxa from different blocks of *Part*(*v*_ℬ_). Searching more aggressively to find a lower *p*-value could increase robustness to the choice of *α*. Additionally, relaxing the requirement of drawing one taxa from each block of the induced quadrapartition allows greater exploration of the space, although it can come at the cost of testing some subsets that do not satisfy condition (1) of Lemma 4, which is a necessary condition even for higher level networks (as this part of the proof does not rely on the network level).

This motivated us to implement a 3f1a-like algorithm that tests taxa around the induced bipartition (i.e., drawing two taxa from either block) until a predetermined number of tests have been performed (see Algorithm 2 in the Supplement). By default, we bound the number of tests conducted by *O*(*n*^2^) so that the time complexity of detecting FPs *O*(*n*^3^*k* + *n* log(*n*)*k*) does not exceed the time complexity of seeking a TOB refinement (note that TREE-QMC has time complexity *O*(*n*^3^*k*) with some assumptions and ASTRAL-III has time complexity *O*(*n*^1.726^*k*^1.726^*D*), where *D* = *O*(*nk*) is the sum of degrees of all unique vertices in the input gene trees). Lastly, we note that combining the bipartition search heuristic with 3f1a yields a consistent method for level-1 networks and does not increase time complexity.

## Acknowledgments

We are grateful to Dr. Cécile Ané, Dr. Elizabeth Allman, Dr. Hector Baños, Dr. John Rhodes, as well as anonymous RECOMB reviewers, for feedback that improved the paper. This material is based upon work supported by the U. S. National Science Foundation (NSF) under Grant No. 2441458 (to EKM) as well as the State of Maryland. The authors thank Francesco Cicconardi for sharing gene trees for the analysis of *Heliconiinae* butterflies and helpful discussions. The authors thank Dr. Claudia Solís-Lemus and Nathan Kolbow for motivation for studying trees of blobs and discussions regarding phylogenetic network simulations, which were initiated at the NSF-funded

Institute for Computational and Experimental Research in Mathematics. Any opinions, findings, and conclusions or recommendations expressed in this material are those of the author(s) and do not necessarily reflect the views of the sponsors.

## Code and Data Availability

TOB-QMC is implemented within TREE-QMC. The source code is publicly available on Github at https://github.com/molloy-lab/TREE-QMC.

