## Supplement for "Quartet-based species tree methods enable fast and consistent tree of blobs reconstruction under the network multispecies coalescent"

May 14, 2026

**References** 41

### 1 Supplemental Methods

#### 1.1 Preliminaries

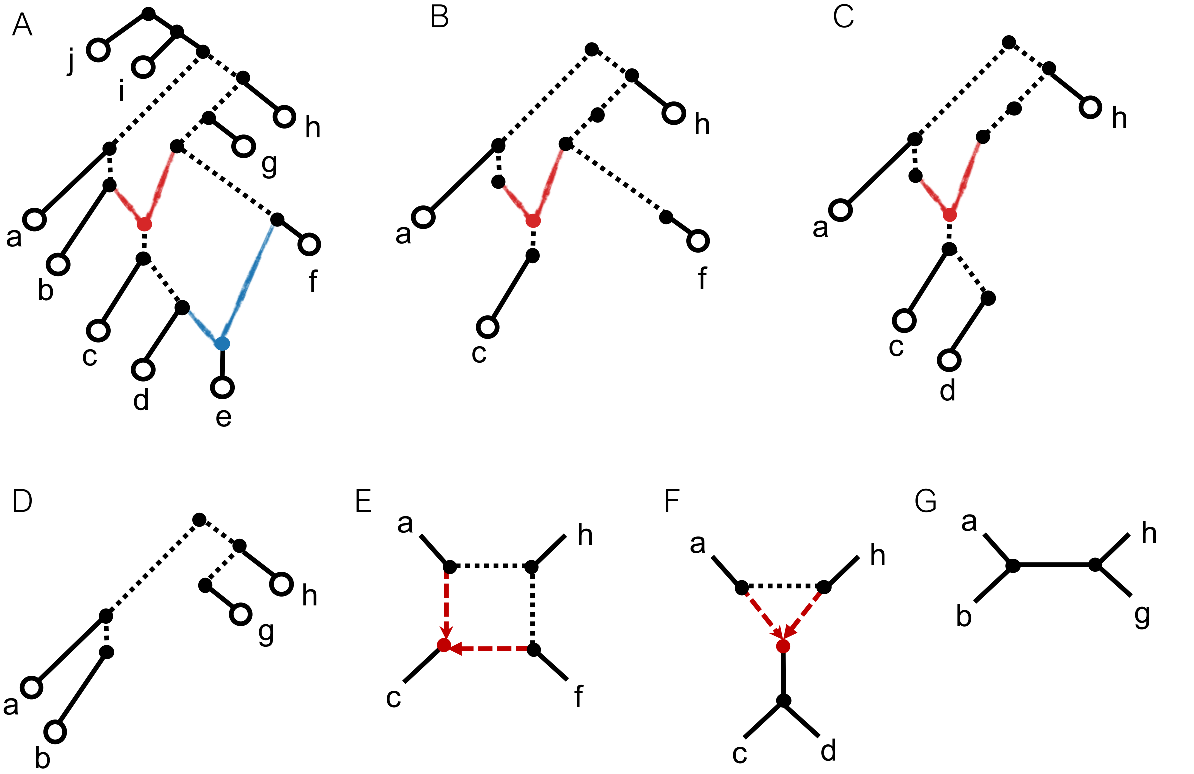

Figure S1: **Restricted and induced subnetworks.** **A)** A level-2 directed phylogenetic network $N^+$. Tree edges are black, and reticulation edges are red or blue to match the color of their shared child vertex. Reticulation edges are part of the blob, as are the dashed edges. **B)** Subnetwork on $\{a, c, f, h\}$; see [3]. **C)** Subnetwork on $\{a, c, d, h\}$. **D)** Subnetwork on to $\{a, b, g, h\}$. **E)** Restriction of $N^+$ to $\{a, c, f, h\}$; see [12]; it contains a 4-blob. **F)** Restriction of $N^+$ to $\{a, c, d, h\}$. **G)** Restriction of $N^+$ to $\{a, b, h, g\}$.

#### 1.2 Network Multispecies Coalescent Model

Our results assume that data (gene trees) are generated under the network multispecies coalescent model, with correlated inheritance. We include a description of the model below for completeness, although it is not necessary for understanding our results.

**Network multispecies coalescent (NMSC).** The *network multispecies coalescent*, with inheritance correlation, was recently described by Fogg *et al.* [8]. The model is parameterized by the inheritance correlation $\rho$, which is used to set the concentration parameter $\alpha = \frac{\rho}{1-\rho}$, as well as a directed phylogenetic network $N^+$ on species/population set $S$ and a set $\theta$ of numeric values associated with the edges of $N^+$:

- Branch lengths $\ell(e) = t(e)/2N_e(e)$ in coalescent units for each arc $e$, where
  - $N_e(e)$ is the effective population size and
  - $t(e)$ is the number of generations
- Inheritance probability (proportion) $\gamma(e)$

A species network $\mathcal{N}^+ = (N^+, \Theta)$ is *metric* if each edge $e$ has length $\ell(e) \geq 0$ and is assigned an inheritance probability $\gamma(e) \in [0,1]$ such that its sum across all reticulation edges with the same child vertex equals 1. Put simply, each vertex $v$ has an associated inheritance probability distribution $\gamma(v) = \{\gamma(u \mapsto v) : u \in \textit{parents}(v)\}$. We also typically assume that $0 < \gamma(e) < 1$ if $e$ reticulation vertex (otherwise $\mathcal{N}^+$ can be simplified) and that all tree edges have positive lengths.

Given a species network $\mathcal{N}^+$, a *gene genealogy* $T^+$ can be generated under the NMSC($\rho$) by sampling lineages, typically one lineage per species is sampled for the gene, and then tracing the ancestry of these lineages backward in time with following the procedure.

- At a **tree node** $v$, all lineages from descendants entering $v$ (or sampled if $v$ is a leaf) can coalesce on the incoming edge $e = p \mapsto v$ with population size $N_e(e)$ and generation time $t(e)$ following the standard the multi-species coalescent (MSC) [15, 32, 23, 25].
- At a **reticulation node** $v$ with $k$ parent nodes, each lineage entering the node is randomly assigned a parent as follows.
    1. Arbitrarily order the lineages entering $v$, denoted $l_1, l_2, \ldots$.
    2. Randomly assign a parent to lineage $l_1$ according to the inheritance probability distribution $\gamma(v)$. Set $i = 1$.
    3. Randomly assign a parent to the next lineage $l_{i+1}$ according to a distribution where the parents assigned to lineages $l_1, \ldots, l_i$ have probability $\frac{1}{\alpha+i}$, and the NULL parent has probability $\frac{\alpha}{\alpha+i}$. If the NULL parent is assigned, randomly assign a parent to $l_{i+1}$ according to the inheritance probability distribution $\gamma(v)$.
    4. Repeat step 3 until all lineages have been assigned a parent.

  This process allows each lineage to have increased preference for having the same ancestry as the previously processed lineages. The correlation between the parent assignment for any pair of lineages $l_i$ and $l_j$ is $\rho = 1/(1+\alpha)$ [8]. All lineages assigned to the same parent $u$ can coalesce on the incoming edge $u \mapsto v$, if $\ell(u \mapsto v) > 0$, following the standard MSC.

Thus, the resulting gene genealogy $T^+$ is a rooted, binary phylogenetic tree with each leaf vertex labeled by the species from which the gene lineage was sampled.

The NMSC with inheritance correlation was introduced by Fogg *et al.* [8] to generalize and unify the two commonly used traditional network coalescent models: the Common Inheritance model and the Independent Inheritance model, which we now define.

**Common Inheritance model.** The *Common Inheritance model* is the NMSC($\rho$) model with $\rho = 1$, which is equivalent to setting $\alpha = 0$. When $\alpha = 0$, all lineages must be assigned to the same parent as lineage $l_1$ because $\frac{\alpha}{\alpha+i} = 0$ for any $i$. Put simply, all lineages sampled for the same gene must evolve within the same display tree.

**Independent Inheritance model.** The *Independent Inheritance model* is the NMSC($\rho$) model with $\rho = 0$, which is equivalent to setting $\alpha = \infty$. When $\alpha = \infty$, the NULL parent is assigned with probability $\frac{\alpha}{\alpha+i} = \frac{1}{1+\frac{i}{\alpha}} = 1$, asymptotically, for any $i$. Thus, all lineages are independently assigned a parent according to the inheritance probability distribution, so there is no ancestry preference among lineages sampled for the same gene.

**Model Comparison.** The Independent Inheritance model is more flexible and thus is the most commonly-used model for species network inference [2, 29, 34, 24, 18, 5]. The Common Inheritance model can be viewed as strong selection acting either for or against gene variants associated with donor populations, which results in all of lineages sampled at a particular locus/gene tracing their ancestry back through the same display tree. As a consequence of this strong assumption, the Common Inheritance model has nice statistical properties [9, 33, 17].

#### 1.3 Any optimal solution to WQC is a TOB refinement almost surely as number of gene trees goes to infinity

**Lemma 4.** *Let $\mathcal{N}^- = (N^-, \Theta)$ be a species network on species set $S$ that satisfies standard assumptions and is T-quartet-nonanomalous. Let $T_N$ denote the TOB of $N^-$, and let $\pi$ be the minimum absolute difference between the qCFs of T-quartets, specifically:*

$$\pi = \min_{Y \in \mathcal{S}_{tree}(T_N)} \min\{|F^*_{\mathcal{N}^-}(q_1^Y) - F^*_{\mathcal{N}^-}(q_2^Y)|, |F^*_{\mathcal{N}^-}(q_1^Y) - F^*_{\mathcal{N}^-}(q_3^Y)|\}. \tag{1}$$

*where $q_1^Y$ is isomorphic to the T-quartet $T_N|_Y$ and $q_2^Y, q_3^Y$ are the other possible binary quartets on $Y$. Let $G_1, G_2, \ldots, G_k$ be a sequence of $k$ independent and identically distributed unrooted gene trees on $S$ sampled from the distribution induced by NMSC given $\mathcal{N}^-$. Then, when the error in the observed qCFs is bounded by $\delta^*$, specifically:*

$$\left| \frac{w_k(q_i^Y)}{k} - F^*_{\mathcal{N}^-}(q_i^Y) \right| < \delta^* = \frac{\pi}{2|f_4(S)|}$$

*for all $i \in \{1, 2, 3\}$ and all $Y \in f_4(S)$, there exists a binary tree $B_N^*$ on $S$ that refines $T_N$ such that $QS_k(B_N^*) > QS_k(T)$ for any binary tree $T$ on $S$ that does not refine $T_N$. Note that we assume the TOB $T_N$ is not an unresolved star tree; otherwise all binary trees on $S$ refine $T_N$ so the result is trivial.*

*Proof.* To begin, observe that $\pi > 0$ by our assumption that the network $\mathcal{N}^-$ has positive branch lengths on its tree edges and is defined because the TOB $T_N$ is not an unresolved star tree, so there is at least one T-quartet. Following upon the proof of Lemma 3, we partition the quartet score of a tree $t$ on $S$ computed from *expected qCFs* into two parts: the **tree part** and the **star part**, specifically:

$$QS(t) = QS^{tree}(t) + QS^{star}(t) = \sum_{Y \in \mathcal{S}_{tree}(T_N)} F^*_{\mathcal{N}^-}(t|_Y) + \sum_{Y \in \mathcal{S}_{star}(T_N)} F^*_{\mathcal{N}^-}(t|_Y) \tag{2}$$

where $\mathcal{S}_{tree}(T_N)$ and $\mathcal{S}_{star}(T_N)$ denote the four-taxon subsets on which $T_N|_Y$ is a binary quartet tree (T-quartet) or an unresolved star tree (B-quartet), respectively. Note that we omit the subscript $k$ from $QS$ to indicate that it is computed from expected qCFs rather than observed qCFs. Next, we partition the quartet score computed from *observed qCFs*, specifically:

$$QS_k(t) = QS_k^{tree}(t) + QS_k^{star}(t) = \sum_{Y \in \mathcal{S}_{tree}(T_N)} \frac{w_k(t|_Y)}{k} + \sum_{Y \in \mathcal{S}_{star}(T_N)} \frac{w_k(t|_Y)}{k}. \tag{3}$$

Now, let $B_N^*$ be some tree on $S$ that maximizes $QS(\cdot)$. By Lemma 3, $B_N^*$ is a refinement of the TOB $T_N$. Thus, we just need to show that when $|w_k(q_i^Y)/k - F^*_{\mathcal{N}^-}(q_i^Y)| < \delta$,

$$QS_k^{star}(T) - QS_k^{star}(B_N^*) < QS_k^{tree}(B_N^*) - QS_k^{tree}(T) \tag{4}$$

as this implies $QS_k(B_N^*) > QS_k(T)$.

***Right side.*** To begin, we give a lower bound for the right side of Equation 4. We need only consider the four-taxon subsets in $\mathcal{S}_{tree}(T_N)$ for which $T|_Y \neq B_N^*|_Y$, denoted $\mathcal{Y}_{tree}$. Then,

$$\begin{aligned} QS_k^{tree}(B_N^*) - QS_k^{tree}(T) &= \sum_{Y \in \mathcal{Y}_{tree}} \left( \frac{w_k(B_N^*|_Y)}{k} - \frac{w_k(T|_Y)}{k} \right) \\ &\geq \sum_{Y \in \mathcal{Y}_{tree}} \big( (F^*_{\mathcal{N}^-}(B_N^*|_Y) - \delta) - (F^*_{\mathcal{N}^-}(T|_Y) + \delta) \big) \\ &\geq |\mathcal{Y}_{tree}|\pi - |\mathcal{Y}_{tree}| 2\delta \end{aligned} \tag{5}$$

by the assumption that $\mathcal{N}^-$ is T-quartet-nonanomalous. The idea is that $T$ is lucky (i.e., errors always increase its score), whereas $B_N^*$ is unlucky (i.e., errors always decrease its score and the gain from preserving each T-quartet in $T_N$ is always just $\pi$).

***Left side.*** Next, we give an upper bound for the left side of the Equation 4. We need only consider the four-taxon subsets in $\mathcal{S}_{star}(T_N)$ for which $T|_Y \neq B_N^*|_Y$, denoted $\mathcal{Y}_{star}$. First, observe that

$$Q_k^{star}(B_N^*) = \sum_{Y \in \mathcal{Y}_{star}} \frac{w_k(B_N^*|_Y)}{k} \geq \sum_{Y \in \mathcal{Y}_{star}} (F_{\mathcal{N}^-}^*(B_N^*|_Y) - \delta) = QS^{star}(B_N^*) - |\mathcal{Y}_{star}|\delta. \tag{6}$$

Second, observe that

$$QS_k^{star}(T) = \sum_{Y \in \mathcal{Y}_{star}} \frac{w_k(T|_Y)}{k} \leq \sum_{Y \in \mathcal{Y}_{star}} (F_{\mathcal{N}^-}^*(T|_Y) + \delta) = QS^{star}(T) + |\mathcal{Y}_{star}|\delta. \tag{7}$$

Again, the idea is that $T$ is lucky (i.e., errors always increase its score), whereas $B_N^*$ is unlucky (i.e., errors always decrease its score). Putting Equations 6 and 7 together gives us an upper bound:

$$QS_k^{star}(T) - QS_k^{star}(B_N^*) \leq \left(QS^{star}(T) + |\mathcal{Y}_{star}|\delta\right) - \left(QS^{star}(B_N^*) - |\mathcal{Y}_{star}|\delta\right) \leq 2|\mathcal{Y}_{star}|\delta \tag{8}$$

noting that $QS^{star}(B_N^*) \geq QS^{star}(T)$ by the proof of Lemma 3.

***Combining bounds together.*** Lastly, we combine Equations 5 and 8 to get

$$QS_k^{star}(T) - QS_k^{star}(B_N^*) \leq 2|\mathcal{Y}_{star}|\delta < |\mathcal{Y}_{tree}|\pi - |\mathcal{Y}_{tree}|2\delta \leq QS_k^{tree}(B_N^*) - QS_k^{tree}(T)$$

where the inner inequality holds when

$$2|\mathcal{Y}_{star}|\delta < |\mathcal{Y}_{tree}|\pi - |\mathcal{Y}_{tree}|2\delta$$
$$\delta < \frac{|\mathcal{Y}_{tree}|}{|\mathcal{Y}_{tree}| + |\mathcal{Y}_{star}|} \frac{\pi}{2}$$

giving us $QS_k(B_N^*) > QS_k(T)$. Because $|\mathcal{Y}_{tree}| \geq 1$ (otherwise $T$ refines the TOB $T_N$) and $|\mathcal{Y}_{tree}| + |\mathcal{Y}_{star}| \leq |\mathcal{S}_{tree}(T_N)| + |\mathcal{S}_{star}(T_N)| = |f_4(S)|$, the inequality holds when $\delta < \delta^* = \pi/2|f_4(S)|$; this is our result. $\square$

#### 1.4 Testing 4-taxon subsets around each branch

**Algorithm 1: Quadrapartition 3 fix, 1 alter (3f1a) search.**

**input :** Set $\mathcal{T}$ of gene trees, Quadrapartition induced by edge $e$, denoted $Quad(e)$
**output:** Minimum $p$-value found during search

$A, B, C, D \leftarrow Quad(e)$;
$a_* \in A$ `// select taxon from` $A$ `uniformly at random`
$b_* \in B$ `// select taxon from` $B$ `uniformly at random`
$c_* \in C$ `// select taxon from` $C$ `uniformly at random`
$d_* \in D$ `// select taxon from` $D$ `uniformly at random`
$p_{min} \leftarrow 1$; $x_{min} \leftarrow \{\}$;
**for** $a \in A$ **do**
  $p \leftarrow$ `quartetTreeTest(qCF(`$\mathcal{T}, \{a, b_*, c_*, d_*\}$`))`;
  **if** $p < p_{min}$ **then** $p_{min} \leftarrow p$; $x_{min} \leftarrow \{a, b_*, c_*, d_*\}$ ;
**for** $b \in B$ **do**
  $p \leftarrow$ `quartetTreeTest(qCF(`$\mathcal{T}, \{a_*, b, c_*, d_*\}$`))`;
  **if** $p < p_{min}$ **then** $p_{min} \leftarrow p$; $x_{min} \leftarrow \{a_*, b, c_*, d_*\}$ ;
**for** $c \in C$ **do**
  $p \leftarrow$ `quartetTreeTest(qCF(`$\mathcal{T}, \{a_*, b_*, c, d_*\}$`))`;
  **if** $p < p_{min}$ **then** $p_{min} \leftarrow p$; $x_{min} \leftarrow \{a_*, b_*, c, d_*\}$ ;
**for** $d \in D$ **do**
  $p \leftarrow$ `quartetTreeTest(qCF(`$\mathcal{T}, \{a_*, b_*, c_*, d\}$`))`;
  **if** $p < p_{min}$ **then** $p_{min} \leftarrow p$; $x_{min} \leftarrow \{a_*, b_*, c_*, d\}$ ;
**return** $p_{min}, x_{min}$

**Algorithm 2: Bipartition heuristic search.** Note that the number of tests is $O(n^2)$ by default to ensure that the time complexity of FP detection does not exceed the time of seeking a TOB refinement with TREE-QMC.

**input :** set $\mathcal{T}$ of gene trees, bipartition induced by edge $e$, denoted $Bip(e)$, and an iteration limit
**output:** minimum $p$-value found during search, sampling 4-taxon subsets around the bipartition induced by $e$

$A, B \leftarrow Bip(e)$;
$q^* \leftarrow ()$;
$i = 0$;
**while** $i <$ *iteration limit* **do**
  $q_{min} \leftarrow (a_1, a_2, b_1, b_2)$ `// randomly pick` $a_1, a_2$ `in` $A$ `and` $b_1, b_2$ `in` $B$
  **repeat**
    $\mathbf{N} \leftarrow \emptyset$;
    **for** $a' \in A \setminus \{a_1, a_2\}$ **do** $\mathbf{N} \leftarrow \mathbf{N} \cup \{(a', a_1, b_1, b_2), (a', a_2, b_1, b_2)\}$ ;
    **for** $b' \in B \setminus \{b_1, b_2\}$ **do** $\mathbf{N} \leftarrow \mathbf{N} \cup \{(a_1, a_2, b', b_1), (a_1, a_2, b', b_2)\}$ ;
    $q_{new} \leftarrow \mathbf{argmin}_{q \in \mathbf{N}}$ `quartetTreeTest(qCF(`$\mathcal{T}, q$`))`;
    $i \leftarrow i + |\mathbf{N}|$;
    **if** *p-value of* $q_{new}$ *is smaller than* $q_{min}$ **then** $q_{min} \leftarrow q_{new}$ ;
  **until** $q_{min}$ *is not updated* ***or*** $i >$ *iteration limit*;
  **if** *p-value of* $q_{min}$ *is smaller than* $q^*$ **then** $q^* \leftarrow q_{min}$ ;
**return** *p-value of* $q^*$

#### 1.5 Simulation Study

##### 1.5.1 Species Network Simulation under Birth-Death-Hybridization model

Species networks were simulated under a birth–death-hybridization process with the R package SiPhyNetwork (v1.1.0) [13], based on the tutorial provided by the package.[1] Specifically, we used the following command:

```
library(ape)
library(SiPhyNetwork)
set.seed(42)
inheritance.fxn <- make.beta.draw(10, 10)
hybrid_proportions <- c(0.5, 0.25, 0.25)
gsa_nets <- sim.bdh.taxa.gsa(m = <upper bound number of taxa>,
                             n = <desired number of taxa>,
                             numbsim = <number of replicates>,
                             lambda = 1,
                             mu = 0.01,
                             nu = <hybridization rate>,
                             hybprops = hybrid_proportions,
                             hyb.inher.fxn = inheritance.fxn)
```

The command above calls the general sampling approach (GSA) function simulates $m$ taxa under the simple sampling approach (SSA) and then samples from time periods where the desired number $n$ of taxa are present. In the command above, $m$ is an upper bound on the number of taxa (otherwise the network is discarded), $n$ is the desired number of taxa, *numbsim* is the number of networks to simulate, $\lambda$ is the speciation/birth rate, $\mu$ is the death/extinction rate, and $\nu$ is the hybridization rate.

We simulated three model conditions based on the number of taxa. For the **50-taxon model condition**, we simulated 10,000 replicates, setting $m = 100$, $n = 50$, and $\nu = 0.02$; this yielded 35 level-0 networks, 447 level-1 networks, and 754 level 2-networks (the remaining networks were higher level). For the **100-taxon model condition**, we simulated 20,000 replicates, setting $m = 150$, $n = 100$, and $\nu = 0.002$; this yielded 95 level-0 networks, 771 level 1-networks, and 1,179 level 2-networks (the remaining networks were higher level). For the **200-taxon model condition**, we simulated 10,000 replicates, setting $m = 500$, $n = 200$, and $\nu = 0.0005$; this yielded 68 level-0 networks, 735 level 1-networks, and 1159 level 2-networks (the remaining networks were higher level). Importantly, we decreased the hybridization rate $\nu$ to ensure that hybridization events were recent (i.e., close to tips), as such events are often of interest to evolutionary biologists (future work should explore deep hybridization events). The other parameters were set based on the tutorial vignette, except that we lowered the death rate to 0.01 to speed up the simulation, which it still took several hours per model condition.

The desired network level cannot be specified explicitly in SiPhyNetwork. Instead, we performed a large number of simulation runs so that we are able to select the networks with desired levels. For 50, 100, and 200 taxon model conditions, we selected 50 networks that were level 1 and level 2; we selected 35, 50, and 25 level-0 networks, respectively, as a control to evaluate the level of incomplete lineage sorting (ILS). Network level was evaluated with the following command:

```
level <- SiPhyNetwork::getNetworkLevel(net)
```

After this selection phase, the true tree of blobs was computed for each network using the algorithm introduced in[11] implemented in PhyloNetworks [30]; specifically, we executed the following command:

```
net = readnewick(input_network_file)
tob = treeofblobs(net)
writenewick(tob, output_tree_of_blobs_file)
```

The summary statistics for the networks and tree of blobs are reported in Table S1. We also evaluated whether the 50-taxon and 100-taxon networks **empirically violated** the quartet-nonanomalous assumption by computing the expected qCFs; the calculations were implemented with the Julia packages

[1] https://cran.r-project.org/web/packages/SiPhyNetwork/vignettes/introduction.html

PhyloNetworks and QuartetNetworkGoodnessFit. If at least one four-taxon subset failed to meet the conditions of Definition 1 in the main text, the entire network was classified as (class 1) quartet-anomalous. The fraction of quartet-anomalous networks are shown in Table S2.

| # of Taxa | Level | # of Reticulations | Inheritance Proportion | # of Nontrivial Blobs | Mean Blob Size | # of internal branches in TOB | Mean # leaves below reticulations |
|---|---|---|---|---|---|---|---|
| 50 | 1 | $1.76 \pm 0.87$ | $0.42 \pm 0.05$ | $1.58 \pm 0.78$ | $8.46 \pm 2.78$ | $39.94 \pm 4.62$ | $1.75 \pm 1.79$ |
| 50 | 2 | $2.70 \pm 0.91$ | $0.42 \pm 0.04$ | $1.56 \pm 0.73$ | $13.18 \pm 3.99$ | $33.60 \pm 4.79$ | $1.94 \pm 1.96$ |
| 100 | 1 | $1.64 \pm 0.75$ | $0.42 \pm 0.06$ | $1.62 \pm 0.73$ | $11.74 \pm 3.45$ | $85.68 \pm 5.25$ | $1.74 \pm 1.39$ |
| 100 | 2 | $2.54 \pm 0.71$ | $0.42 \pm 0.04$ | $1.52 \pm 0.68$ | $18.11 \pm 5.55$ | $78.04 \pm 5.30$ | $1.66 \pm 0.82$ |
| 200 | 1 | $1.98 \pm 0.87$ | $0.41 \pm 0.06$ | $1.96 \pm 0.86$ | $13.53 \pm 3.42$ | $180.28 \pm 8.60$ | $2.29 \pm 3.65$ |
| 200 | 2 | $2.94 \pm 0.87$ | $0.41 \pm 0.04$ | $1.88 \pm 0.80$ | $19.47 \pm 5.23$ | $171.42 \pm 7.87$ | $1.86 \pm 1.05$ |

Table S1: Statistics (mean $\pm$ standard deviation) for species networks simulated under each model condition. Note that the lower of the two inheritance proportions is taken.

| # of taxa | Network Level | Branch Scale Factor (ILS) | # Replicates | Fraction of Class 1 Quartet Anomalous |
|---|---|---|---|---|
| 50 | 1 | 0.25x | 50 | 0.320 |
| 50 | 1 | 0.5x | 50 | 0.220 |
| 50 | 1 | 1.0x | 50 | 0.180 |
| 50 | 1 | 2.0x | 50 | 0.060 |
| 50 | 2 | 0.25x | 50 | 0.340 |
| 50 | 2 | 0.5x | 50 | 0.320 |
| 50 | 2 | 1.0x | 50 | 0.220 |
| 50 | 2 | 2.0x | 50 | 0.160 |
| 100 | 1 | 0.25x | 47 | 0.319 |
| 100 | 1 | 0.5x | 50 | 0.160 |
| 100 | 1 | 1.0x | 50 | 0.120 |
| 100 | 1 | 2.0x | 50 | 0.060 |
| 100 | 2 | 0.25x | 50 | 0.500 |
| 100 | 2 | 0.5x | 50 | 0.400 |
| 100 | 2 | 1.0x | 50 | 0.260 |
| 100 | 2 | 2.0x | 50 | 0.100 |

Table S2: The fraction of class 1 quartet-anomalous out of the replicate networks simulated under each model condition. Note that 3 replicates of model condition of 100 taxa, 0.25x ILS level and network level of 1, we ran out 256GB memory limit for computing expected qCFs and thus are not included in this analysis.

##### 1.5.2 Gene Tree Simulation under Network Multispecies Coalescent

Gene trees were simulated from species networks under NMSC (Independent Inheritance) with the Julia package PhyloCoalSimulations (v1.0.0) [8] based on the tutorial provided by the package.[2] Specifically we used the following command:

```
using PhyloNetworks
using PhyloCoalSimulations
using PhyloPlots
foreach(readdir(<directory>))do f
    println("Object: ", f)
    net = readTopology(readline(<directory> * f))
    trees = simulatecoalescent(<species network>, <number of gene trees>,
        <number of individuals per species>)
    writeMultiTopology(trees, <directory> * f * ".truegenetrees")
end
```

setting the number of individuals sampled per species to 1 and the number of gene trees to 1000.

To vary the amount of ILS, we multiplied branch lengths in the species tree by 2.0, 1.0, 0.5, and 0.25 prior to simulate species trees; these model conditions are also referred to as very low, low, medium, and high ILS, respectively. The amount of ILS can be evaluated for level-0 networks, as the normalized Robinson-Foulds (RF) distance [27] between gene trees and the level-0 networks (species trees), because there is no gene flow. The average gene tree discordance due to ILS ranged from 35% to 95% (Fig. S2), with very low, low, medium, and high ILS model conditions have mean discordance of 40–50%, 60–70%, ∼80%, and ∼90%, respectively.

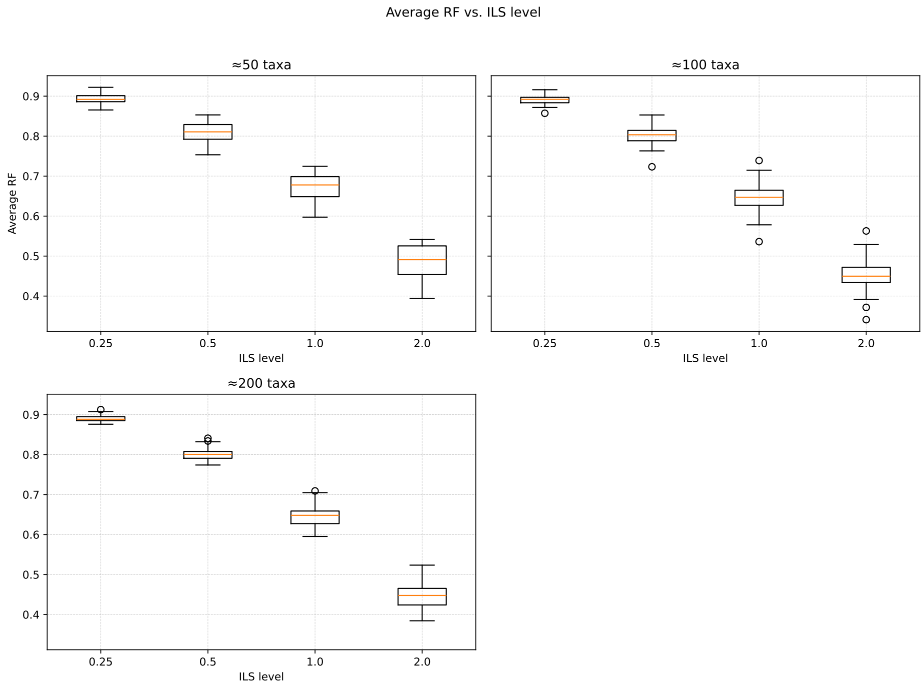

Figure S2: **Empirical ILS Level.** The ILS level was evaluated empirically using level-0 species networks, which are just species trees. The $x$-axis is the species tree height scaling factor. The $y$-axis is the mean gene tree discordance (i.e., the normalized Robinson-Foulds distance between the true species tree and each true gene tree, averaged across all 1000 true gene trees). Box plots are based on empirical ILS level for different replicates.

[2] https://juliaphylo.github.io/PhyloCoalSimulations.jl/stable/man/getting_started

##### 1.5.3 Tree of Blob Reconstruction and Evaluation

**TINNiK.** We ran TINNiK [1], as implemented in MSCquartets v3.2 [26] based on the tutorial provided by the package.[3] Specifically, we used the following command:

```
args=commandArgs(trailingOnly=TRUE)
library('MSCquartets')
gts = read.tree(file=args[1])
output = TINNIK(gts, alpha=as.numeric(args[3]),
              beta=as.numeric(args[4]), plot=FALSE)
write.tree(output$ToB, file=args[2])
```

where the first argument is the input gene trees, the second argument is the output tree of blobs, the third argument is the $\alpha$ hyperparameter for the tree-test, and the fourth argument is the $\beta$ hyperparameter for the star-test. We ran TINNiK on each data set with varying the $\alpha$ and $\beta$ values, as described in the next section.

**TOB-QMC.** We implemented TOB-QMC, within TREE-QMC (https://github.com/molloy-lab/TREE-QMC). Two commands are required to run TOB-QMC.

The first command reconstructs a base tree with TREE-QMC [10] and then annotates each branch with the $p$-value for the star-test given the average qCFs around the branch as well as the minimum $p$-value found for the tree-test.

```
./tree-qmc \
    -i <input gene trees> \
    -o <output tree with p-values for each branch> \
    --iter_limit_blob <iteration limit> --store_pvalue
```

The star-tests and tree-tests are performed by invoking the same R code as TINNiK. The minimum $p$-value is found by testing different 4-taxon subsets around the branch, searching for a subset that yields the lowest $p$-value. Depending on the experiment, we ran TOB-QMC using one of four search modes:

- **exhaustive**: Tests all subsets taking 2 taxa from each side of the bipartition (invoked by setting the iteration limit to 0)
- **default**: Tests $2n^2$ subsets according to bipartition search algorithm (Algorithm 2) (invoked by setting the iteration limit 5000, 20000, and 80000 for 50, 100, and 200 taxa, respectively)
- **fast**: Tests $n^2/4$ subsets according to bipartition search algorithm (Algorithm 2) (invoked by setting the iteration limit to 625, 2500, and 10,000 for 50, 100, and 200 taxa, respectively)
- **3f1a** Tests $n$ subsets according to the 3f1a search algorithm (Algorithm 1) (invoked by using the flag `--3f1a`)

The second command (below) contracts branches in the output of command 1 based on the user-provided hypothesis testing hyperparameters: $\alpha$ and $\beta$.

```
./tree-qmc \
    -i <input tree with p-values for each branch> \
    -o <output tree of blobs> \
    --blob \
    --alpha <alpha threshold> \
    --beta <beta threshold> \
    --load_pvalue
```

[3] https://cran.r-project.org/web/packages/MSCquartets/vignettes/TINNIK.html

We ran this second step of TOB-QMC on each data set varying the $\alpha$ and $\beta$ values, as described in the next section. In the version of TOB-QMC tests, the star-test is applied to average qCFs around the branch, as described by [28]; however, it is faster, and more principled, to apply them to the same qCFs that give the minimum $p$-value found to rule out singularities. Future versions of TOB will make this change, we do not anticipate that it would impact our results. The runtime reported for TOB-QMC only includes the time for command 1 because command 2 is very fast. See the TOB-QMC tutorial for details.

**Evaluation Metrics.** We evaluated estimated TOBs by computing average of the false negative branch rate (FNR) and the false positive branch rate (FPR). FNR is the number of branches from the true TOB that are missing from the estimated TOB divided by the number of internal branches in the true TOB. FPR is the number of branches in the estimated TOB that are missing from the true TOB divided by the number of internal branches in the estimated TOB. In most phylogenetic studies, the true and estimated trees are binary, so FPR equals FNR, so only one of them needs to be reported (they are also equivalent to the normalized RF error). Because TOBs are non-binary, we report FPR and FNR separately (although we use their average for tuning hyperparameters).

##### 1.5.4 Sequence Simulation under the GTR+F+G model

Sequences, specifically multiple sequences alignments (MSAs), were simulated from each true gene tree under GTR+F+G model using AliSim [19]. The simulation procedure mostly followed the ASTRAL-II study [21]; specifically, base frequences were drawn from Dirichlet(36,26,28,32), GTR substitution parameters were drawn from a Dirichlet(16,3,5,5,6,15), and rates-across-sites shape parameters were drawn from an exponential distribution with rate 1.2, with values below 0.1 discarded. In contrast to the ASTRAL-II study, we fixed the alignment length to be 100 sites rather than drawn it from a log normal distribution.

##### 1.5.5 Gene Tree Estimation

Gene trees were estimated from MSAs as follows. First, we ran Modelfinder [14] to find the best nucleotide substitution model. The GTR+F+ASC+G4 was the most frequently selected model in our data (Fig. S3, S4, S5). After selecting the best fit model, gene trees were estimated using IQ-TREE (v3.0.1) [20, 22]. Mean gene tree estimation error (GTEE) ranged from 40% to 70% (Fig. S6), with very low, low, medium, and high ILS model conditions have mean discordance of ∼45%, ∼48%, ∼53%, and ∼60%, respectively. IQ-TREE sometimes failed to run (see an example error message below); however, all data sets included at least 900 estimated gene trees, so we did not explore the issue further.

```
ERROR: modelfactory.cpp:1131: virtual bool ModelFactory::initFromNestedModel(std::map<std:
ERROR: STACK TRACE FOR DEBUGGING:
ERROR: 1   funcAbort()
ERROR: 2   ()
ERROR: 3   gsignal()
ERROR: 4   abort()
ERROR: 5   ()
ERROR: 6   ModelFactory::initFromNestedModel(std::map<std::__cxx11::basic_string<char, std
ERROR: 7   CandidateModel::evaluate[abi:cxx11](Params&, ModelCheckpoint&, ModelCheckpoint&,
ERROR: 8   CandidateModelSet::test(Params&, PhyloTree*, ModelCheckpoint&, ModelsBlock*, int
ERROR: 9   runModelFinder(Params&, IQTree&, ModelCheckpoint&, std::__cxx11::basic_string<ch
ERROR: 10   startTreeReconstruction(Params&, IQTree*&, ModelCheckpoint&)
ERROR: 11   runPhyloAnalysis(Params&, Checkpoint*, IQTree*&, Alignment*&)
ERROR: 12   runPhyloAnalysis(Params&, Checkpoint*)
ERROR: 13   main()
ERROR: 14   __libc_start_main()
ERROR: 15   ()
ERROR:
ERROR: *** IQ-TREE CRASHES WITH SIGNAL ABORTED
```

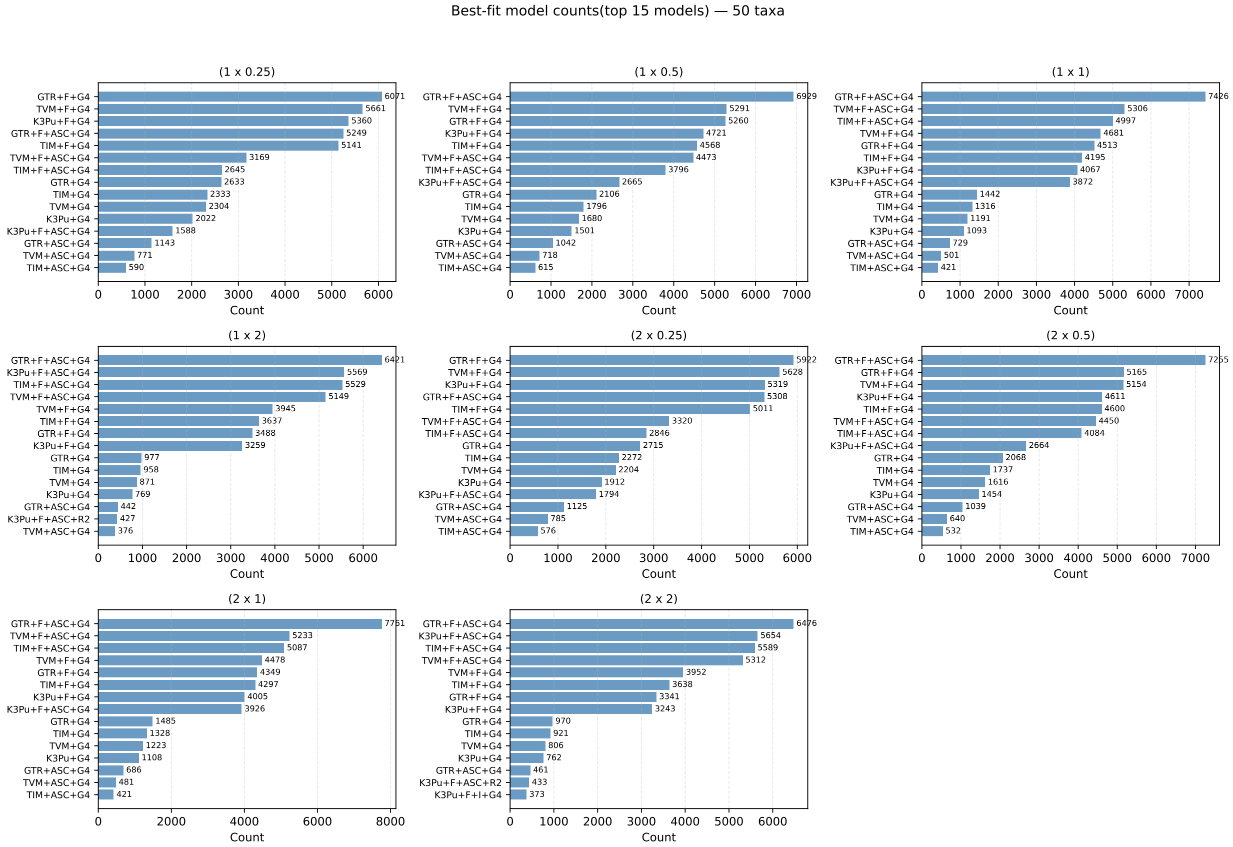

Figure S3: **Gene tree estimation models for 50-taxon data sets.**

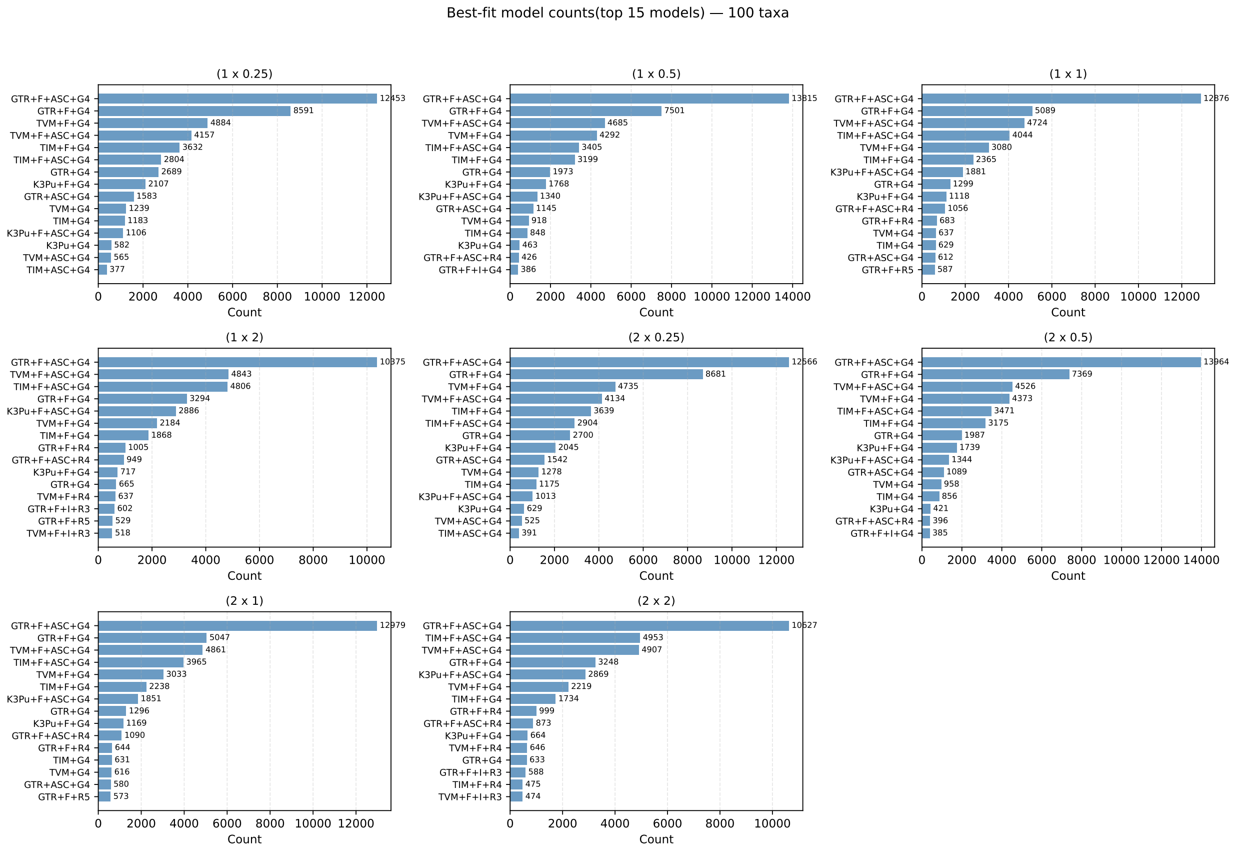

Figure S4: **Gene tree estimation models for 100-taxon data sets.**

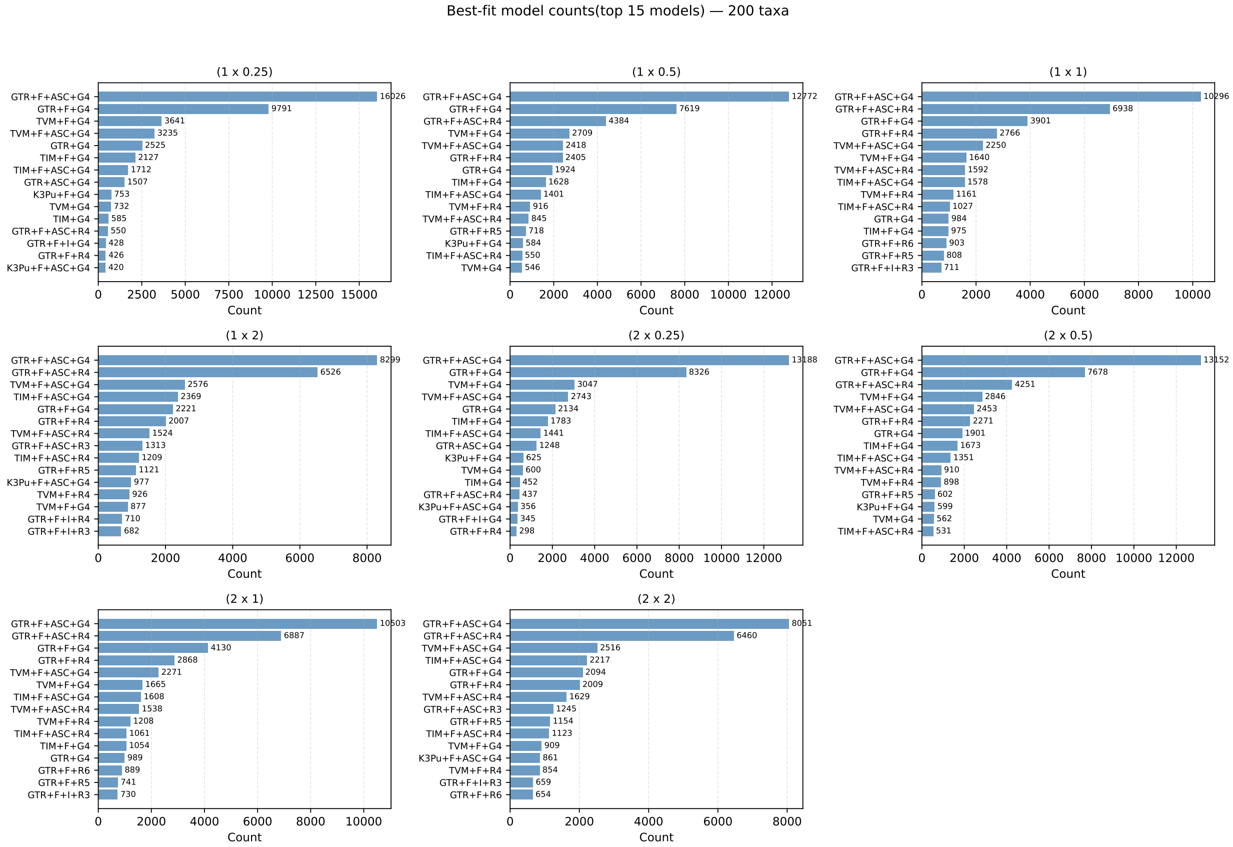

Figure S5: **Gene tree estimation models for 200-taxon data sets.**

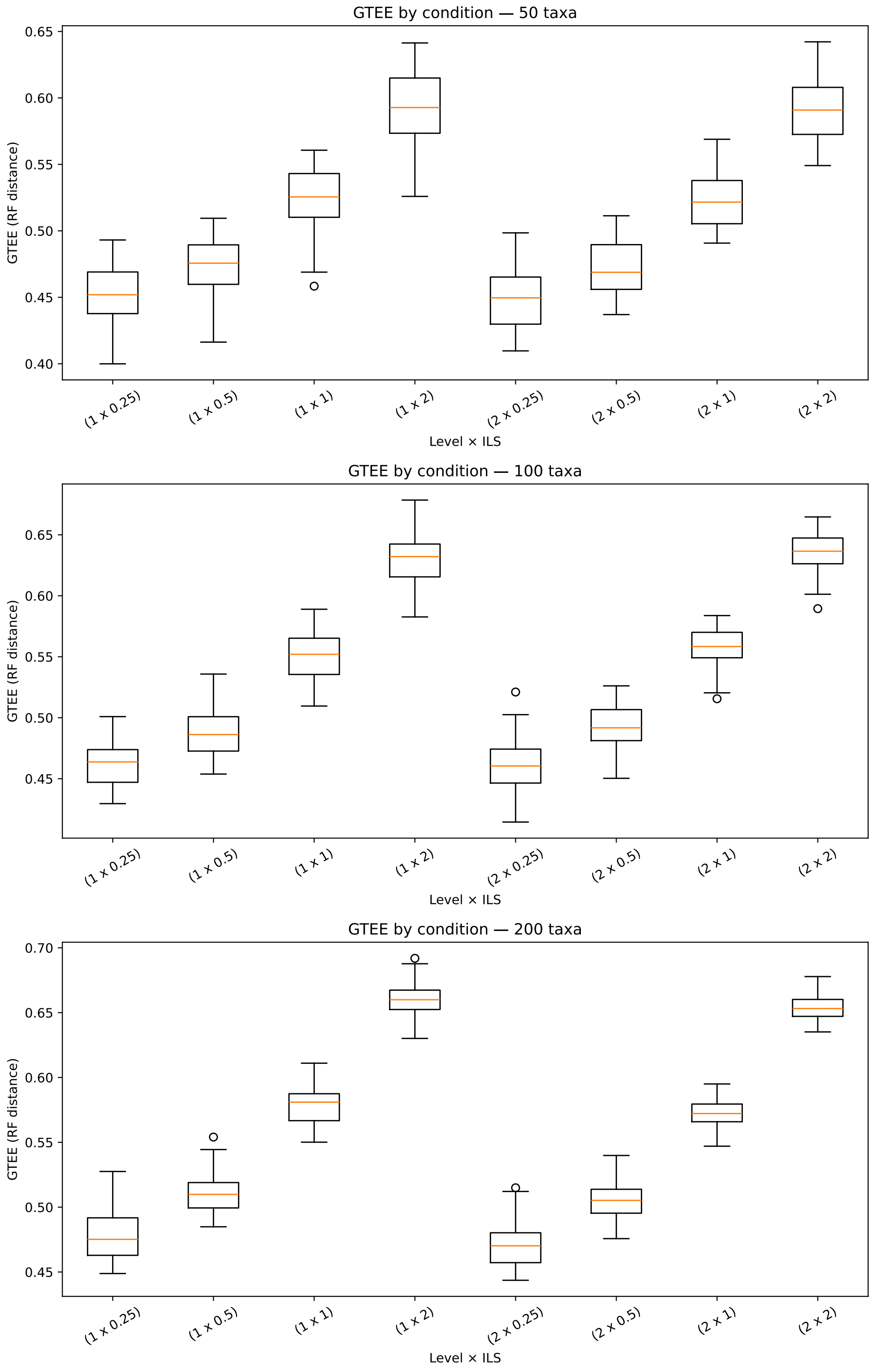

Figure S6: **Empirical GTEE Level.** The $x$-axis is the network level and species tree height scaling factor. The $y$-axis is the mean gene tree estimation error (i.e., the normalized Robinson-Foulds distance between the true and estimated tree, averaged across all 1000 gene trees). Box plots are based on empirical mean GTEE level for different replicates.

##### 1.5.6 Runtime Evaluation

To further evaluate the scalability of TINNiK and TOB-QMC, we benchmarked them on 50-, 100-, 200-taxon datasets with estimated gene trees. Runtime experiments were performed on compute nodes outfitted with AMD EPYC-7313 cores and 2TB of RAM. The AMD EPYC-7313 cores are configured as a dual-socket with 4 NUMA nodes of 256GB per socket. TINNiK and TOB-QMC were run with 1 thread, 256 GB of memory, and a maximum wall clock time of 48 hours. After our experiments we learned that TINNiK can leverage multiple cores to compute the quartet concordance factors. Similar parallelism could be employed by TOB-QMC, as hypothesis testing can be conducted on each branch in parallel and 4-taxon subsets around a given branch can be evaluated in parallel. That being said, testing in single-threaded mode is most inline with our time complexity analysis. When collecting runtime data, we requested exclusive access of these compute nodes and local memory binding via slurm scheduler to reduce potential detrimental effects on runtime measurements.

Ultimately, we were unable to run TINNiK on 200-taxon data sets because either it did not complete within the 48 wall-clock time limit

```
    Loading required package: ape
Loading required package: phangorn
Analyzing 200 taxa: l100, l101, l102, l103, l104, l105, l106, l107, l108, l109, l110, l111, l112,
     l113, l114, l115, l116, l117, l118, l119, l120, l121, l122, l123, l124,...(see output table
    for full list)
Counting occurrences of displayed quartets for 64684950 four-taxon subsets of 200 taxa across
    1000 gene trees.
Applying hypothesis test for model T3 to 64684950 quartets.
Applying hypothesis test for star tree model to 64684950 quartets.
slurmstepd: error: *** JOB 5533599 ON cbcb14 CANCELLED AT 2025-10-12T16:24:12 DUE TO TIME LIMIT
    ***
```

or it ran out of memory (256GB).

```
Loading required package: ape
Loading required package: phangorn
Analyzing 200 taxa: l100, l101, l102, l103, l104, l105, l106, l107, l108, l109, l110, l111, l112,
     l113, l114, l115, l116, l117, l118, l119, l120, l121, l122, l123, l124,...(see output table
    for full list)
Counting occurrences of displayed quartets for 64684950 four-taxon subsets of 200 taxa across 995
     gene trees.
Applying hypothesis test for model T3 to 64684950 quartets.
Applying hypothesis test for star tree model to 64684950 quartets.
slurmstepd: error: Detected 1 oom_kill event in StepId=5533600.batch. Some of the step tasks have
     been OOM Killed.
```

Given limited compute resources, we also did not run TOB-QMC-exhaustive on 200-taxon datasets, as it took around 30 hours.

##### 1.5.7 Hyperparameter Tuning – Hypothesis Testing Thresholds

An issue with running both TINNiK and TOB-QMC is that testing thresholds need to be set for the star-test and the tree-test, denoted $\beta$ and $\alpha$, respectively. In [1], TINNiK was evaluated on a species network with 23 species, for which the tree of blobs had 11 internal branches; the authors found that setting $\beta = 1$ and $\alpha$ between $1e-7$ and $1e-3$ yielded the correct TOB, but setting $\alpha$ higher gave an over-resolved TOB and setting it lower gave an under-resolved TOB. More recently, Kolbow et al. [16] ran TINNiK on data sets with 30 to 200 species, setting $\alpha = 0.01$ and $\beta = 0.99$. To evaluate the impact of $\alpha$ and $\beta$, we ran TINNiK and TOB-exhaustive on 50- and 100-taxon data sets (with true gene trees), setting $\beta \in \{0.9, 0.95, 1.0\}$ $\alpha \in \{1e-9, 1e-8, 1e-7, 1e-6, 1e-5\}$ (Figs. S7, S8, S9, S10). Overall, the impact of different settings of $\beta$ had a minor impact on method performance. The impact of $\alpha$ was greater with $1e-5$ resulting in noticeably worse performance. Setting $\beta = 0.95$ and $\alpha = 1e-7$ gave reasonable results for both methods and thus were used in further experiments.

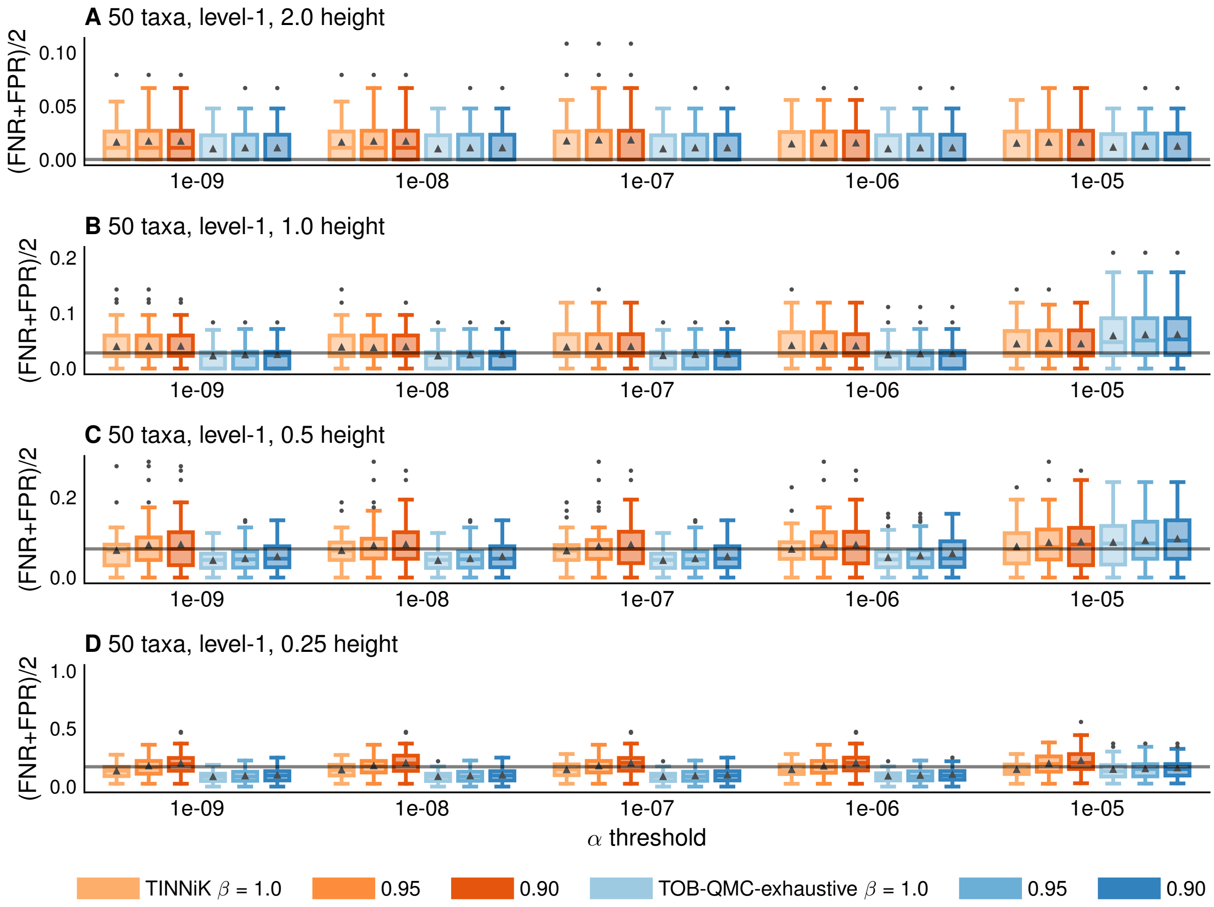

Figure S7: **Impact of $\alpha$ and $\beta$ thresholds on 50-taxon, level-1 data sets.** The solid line is the error for TINNiK setting $\beta = 0.95$ and $\alpha = 1e-7$.

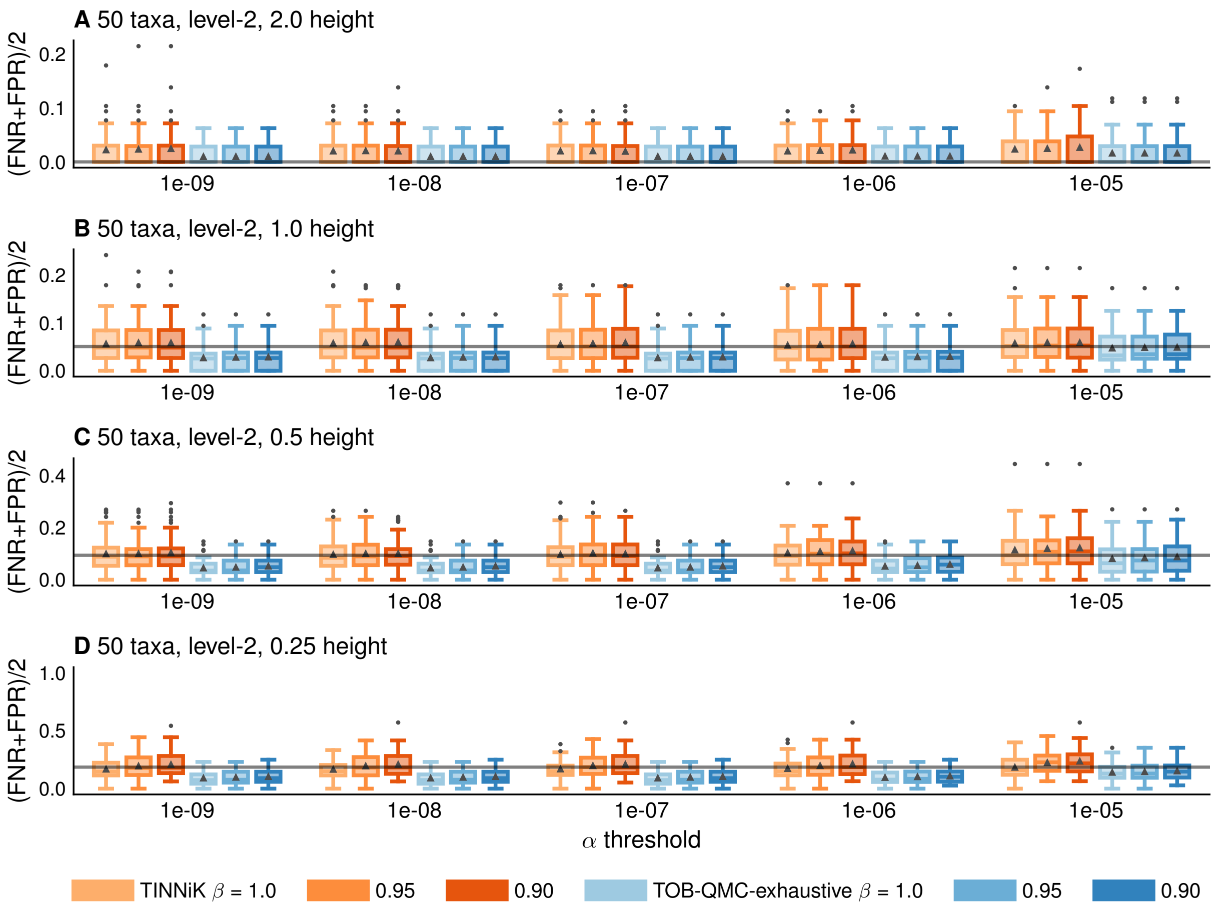

Figure S8: **Impact of $\alpha$ and $\beta$ thresholds on 50-taxon, level-2 data sets.** The solid line is the error for TINNiK setting $\beta = 0.95$ and $\alpha = 1e-7$.

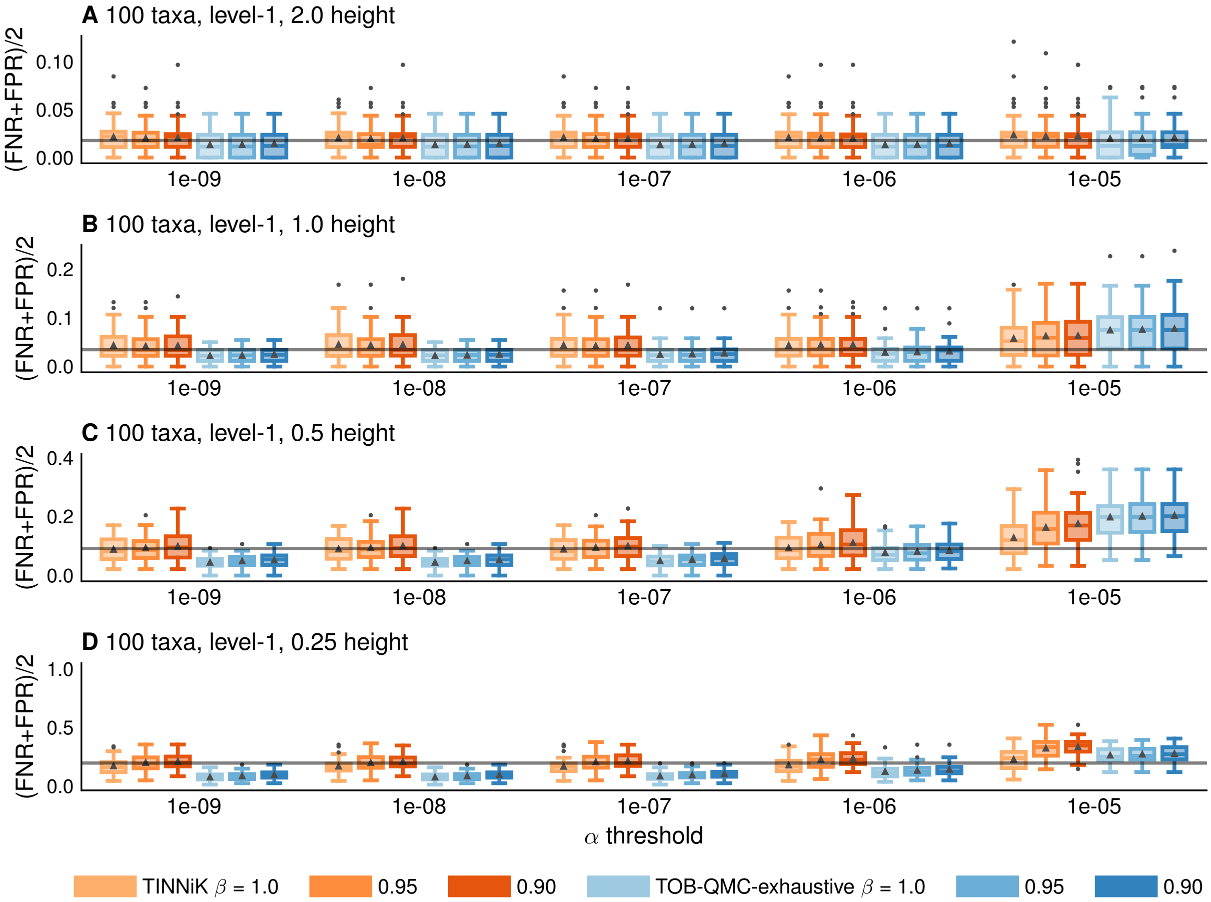

Figure S9: **Impact of $\alpha$ and $\beta$ thresholds on 100-taxon, level-1 data sets.** The solid line is the error for TINNiK setting $\beta = 0.95$ and $\alpha = 1e - 7$.

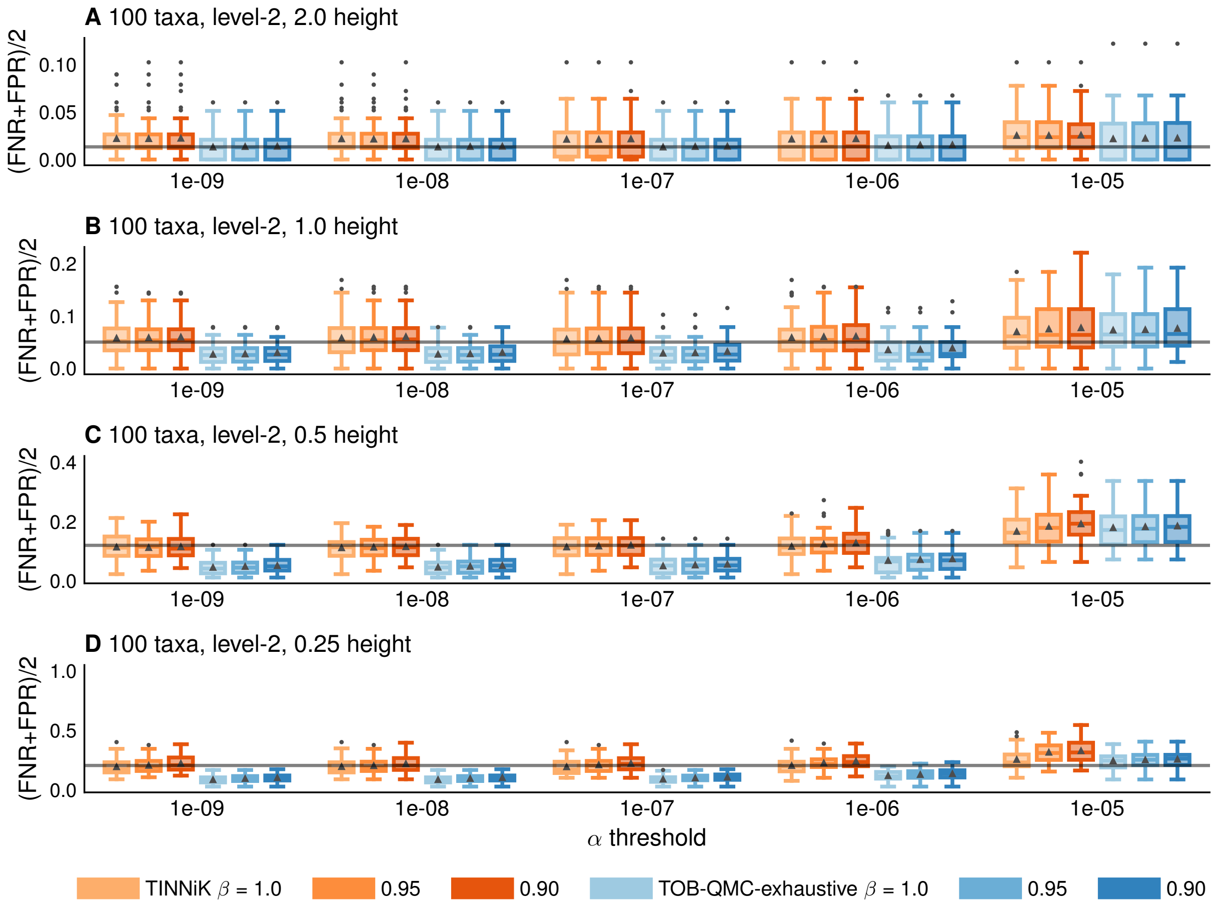

Figure S10: **Impact of $\alpha$ and $\beta$ thresholds on 100-taxon, level-2 data sets.** The solid line is the error for TINNiK setting $\beta = 0.95$ and $\alpha = 1e-7$.

##### 1.5.8 Hyperparameter Tuning – TOB-QMC's Search Algorithm

We also evaluated the impact of TOB-QMC's search algorithm on 50- and 100-taxon data sets (true gene trees only). All TOB-QMC search algorithms achieve similar FN rates and outperform TINNiK; however, they differ in terms of FP rates. The 3f1a search, although consistent, has the highest FP rate. In contrast, bipartition search with quadratic iteration limit ($2n^2$) achieves similar FP rate to exhaustive search. In our evaluation study, TOB-QMC is run with bipartition search and quadratic iteration limit, as these default and fast modes are much faster than exhaustive search and more accurate than 3f1a search.

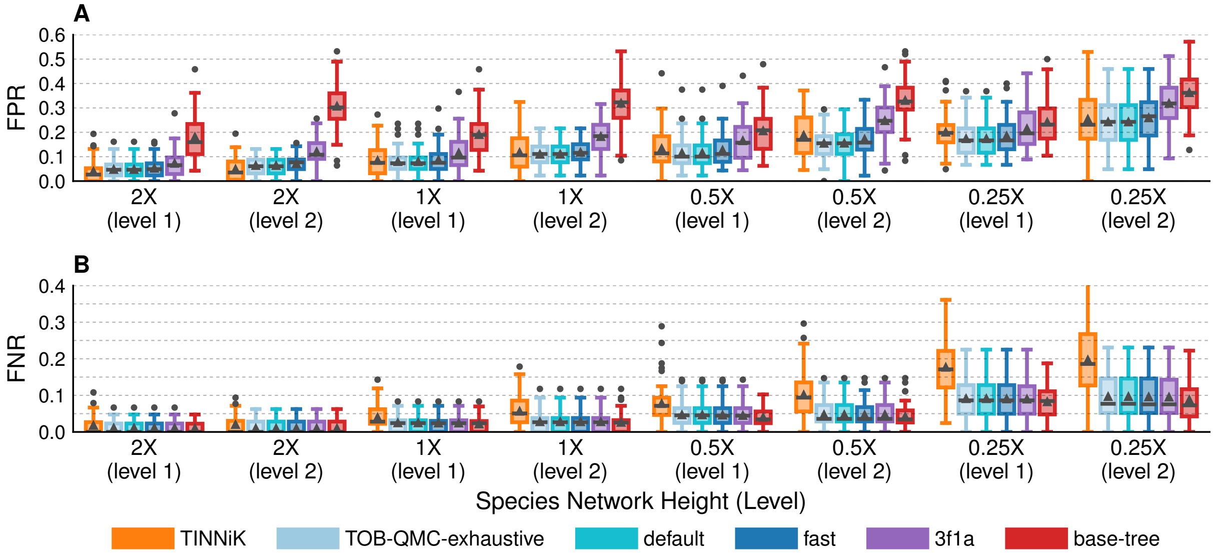

Figure S11: **Impact of TOB's search heuristic on 50-taxon data sets.**

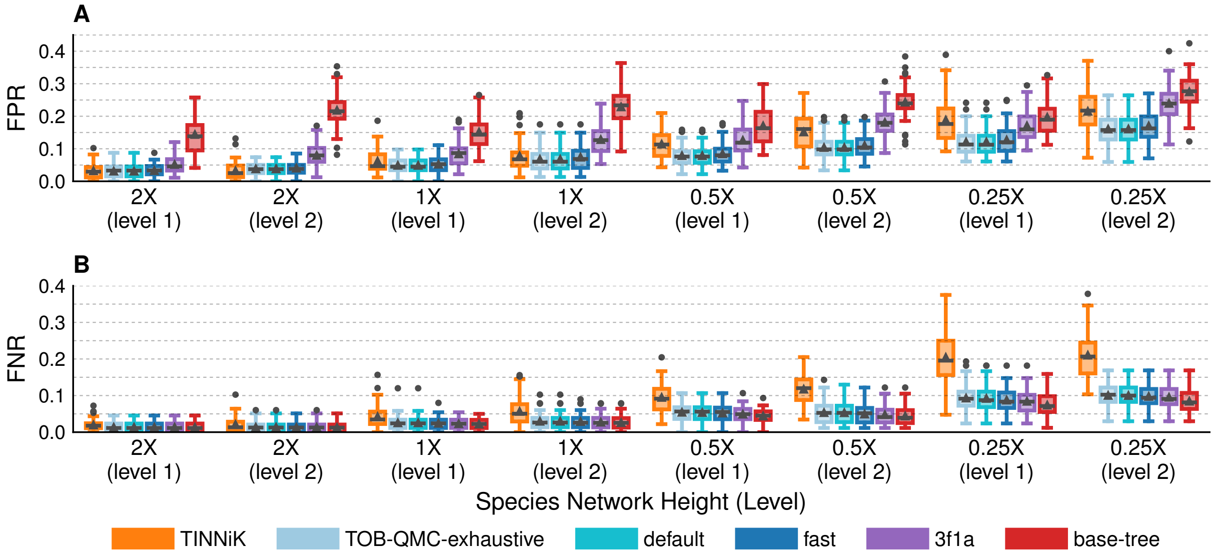

Figure S12: **Impact of TOB's search heuristic on 100-taxon data sets.**

##### 1.5.9 R environments

We downloaded `R (version 4.5.1)` by the following command:

```
wget https://cran.r-project.org/src/base/R-4/R-4.5.1.tar.gz
```

We installed `Rcpp, Rinside, TINNiK` R packages by the following commands:

```
Rscript -e "install.packages('Rcpp', repos='https://cloud.r-project.org/')"
Rscript -e "install.packages('RInside', repos='https://cloud.r-project.org/')"
Rscript -e "install.packages('MSCquartets', repos='https://cloud.r-project.org/')"
```

### 2 Supplemental Results

#### 2.1 Simulation study varying network level and ILS level

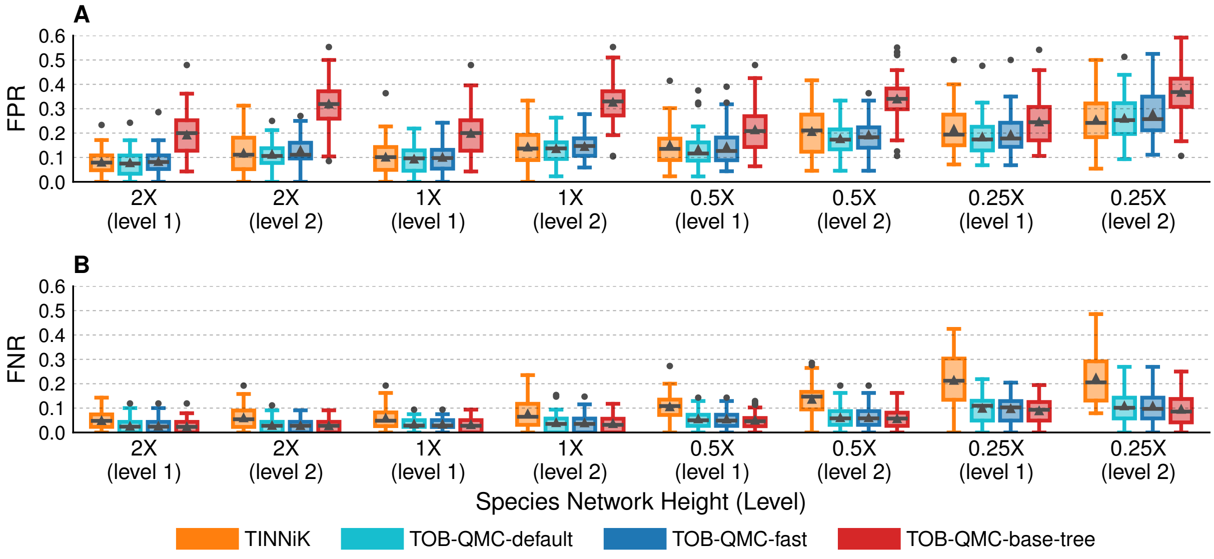

Figure S13: **Impact of network level and ILS level on 50-taxon data sets.**

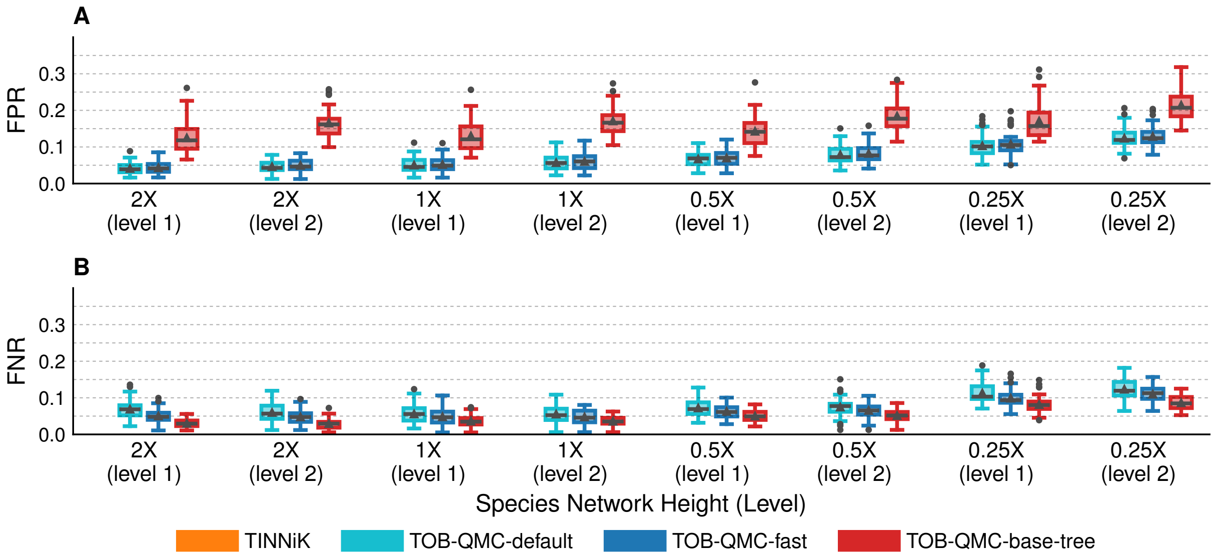

Figure S14: **Impact of network level and ILS level on 200-taxon data sets.** Note that TINNiK could not be run on 200-taxon data sets due to exceeding the wall clock time or memory.

#### 2.2 Simulation study varying number of taxa

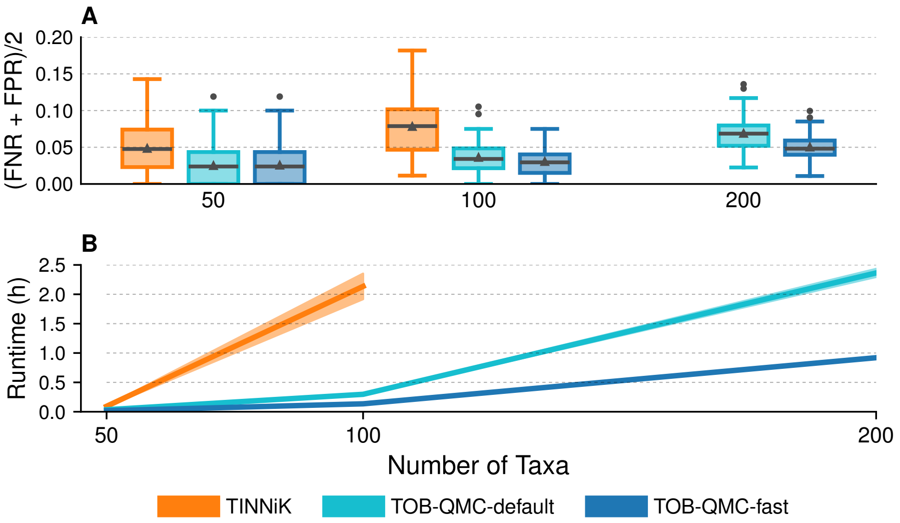

Figure S15: **Impact of number of taxa on level-1 network data sets with scale factor 2.0 (ILS: ∼45%) and estimated gene trees.** Note that TINNiK could not be run on 200-taxon data sets due to exceeding the wall clock time or memory.

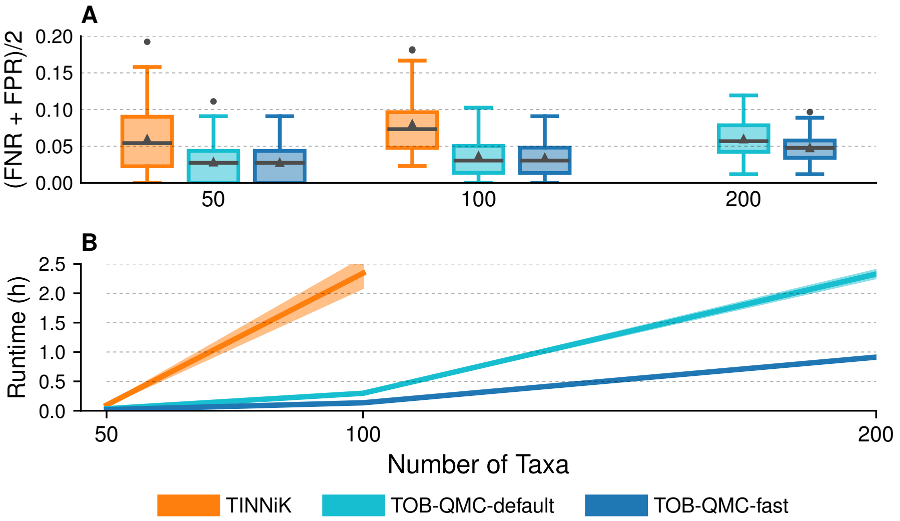

Figure S16: **Impact of number of taxa on level-2 network data sets with scale factor 2.0 (ILS: ∼45%) and estimated gene trees.** Note that TINNiK could not be run on 200-taxon data sets due to exceeding the wall clock time or memory.

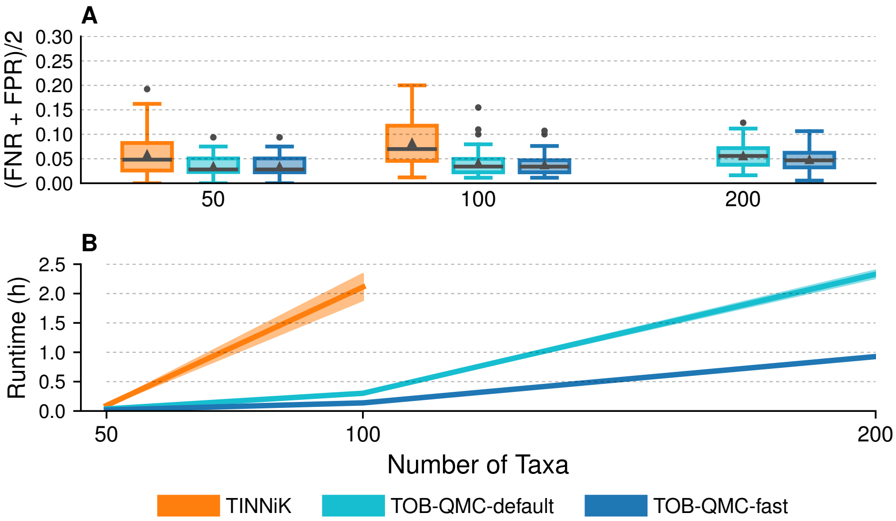

Figure S17: **Impact of number of taxa on level-1 network data sets with scale factor 1.0 (ILS: ∼65%) and estimated gene trees.** Note that TINNiK could not be run on 200-taxon data sets due to exceeding the wall clock time or memory.

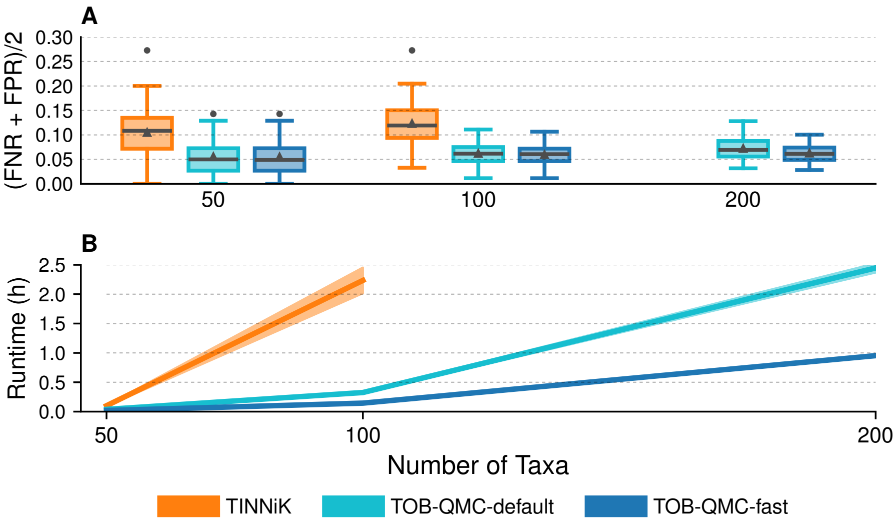

Figure S18: **Impact of number of taxa on level-1 network data sets with scale factor 0.5 (ILS: ∼80%) and estimated gene trees.** Note that TINNiK could not be run on 200-taxon data sets due to exceeding the wall clock time or memory.

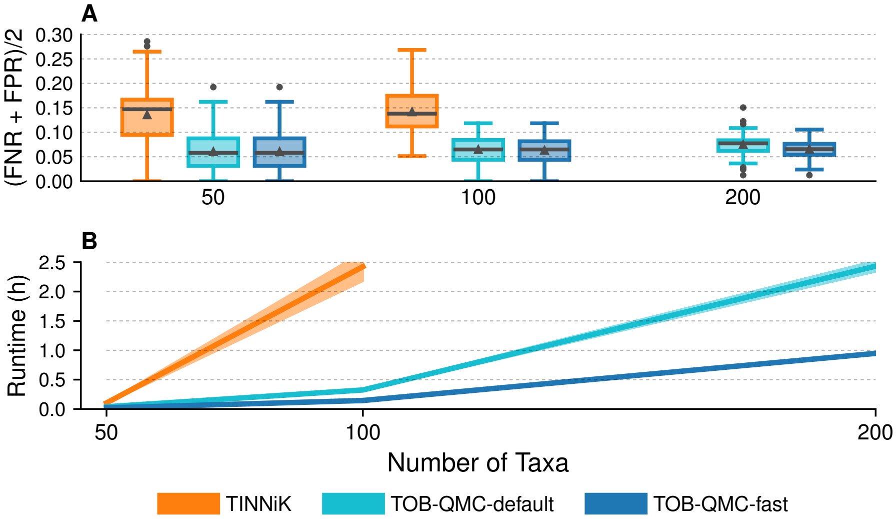

Figure S19: **Impact of number of taxa on level-2 network data sets with scale factor 0.5 (ILS: ∼80%) and estimated gene trees.** Note that TINNiK could not be run on 200-taxon data sets due to exceeding the wall clock time or memory.

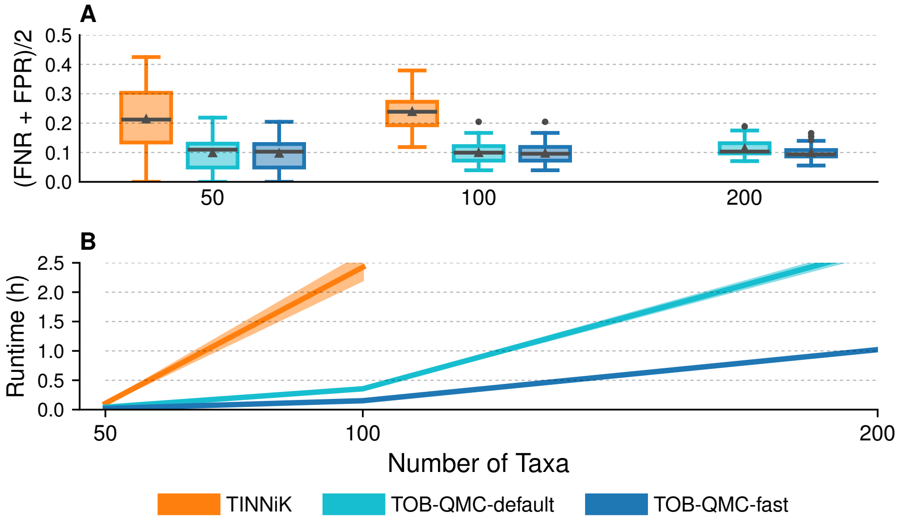

Figure S20: **Impact of number of taxa on level-1 network data sets with scale factor 0.25 (ILS: ∼90%) and estimated gene trees.** Note that TINNiK could not be run on 200-taxon data sets due to exceeding the wall clock time or memory.

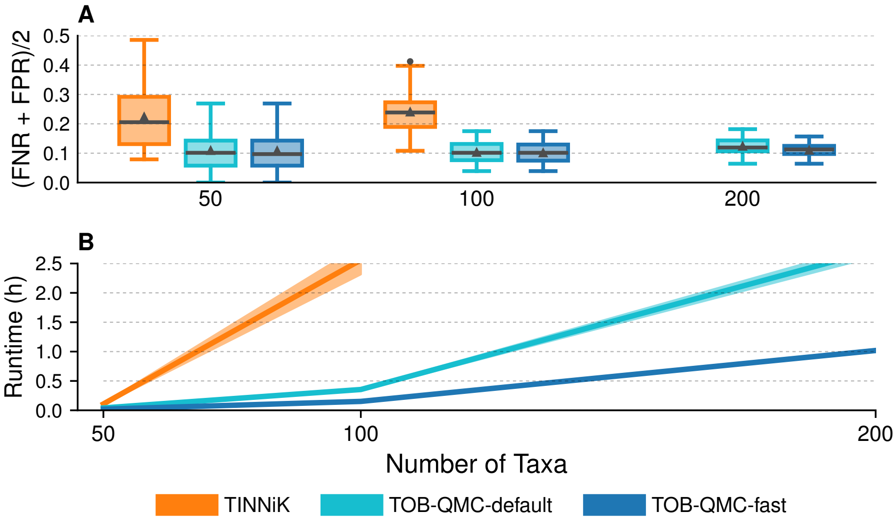

Figure S21: **Impact of number of taxa on level-2 network data sets with scale factor 0.25 (ILS: ~90%) and estimated gene trees.** Note that TINNiK could not be run on 200-taxon data sets due to exceeding the wall clock time or memory.

#### 2.3 Impact of violation of class 1 quartet-nonanomalous assumption

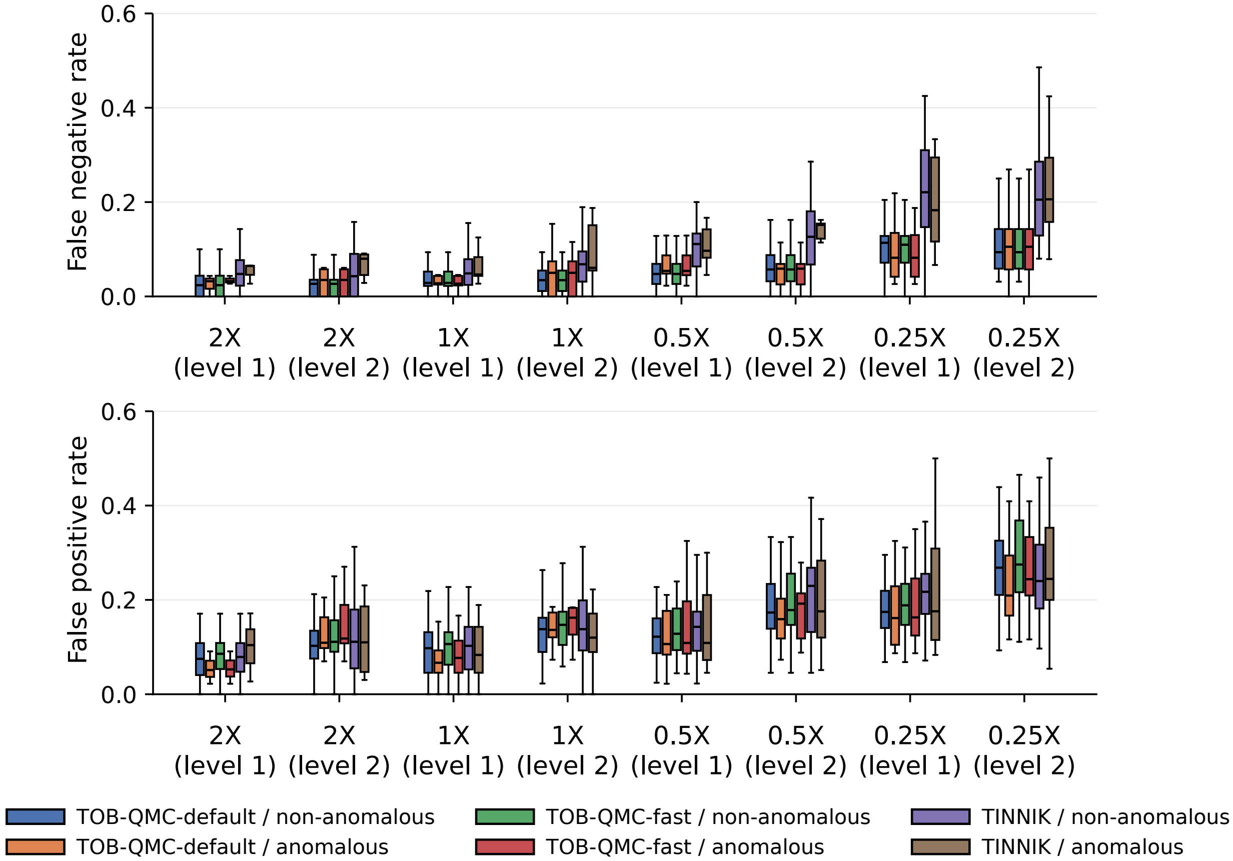

Figure S22: **Impact of violation of class 1 quartet-nonanomalous assumption on 50-taxon networks.** Model condition on $x$-axis indicates network level and branch scaling factor (lower factor is higher ILS). Boxplots indicate method used for TOB reconstruction given estimated gene trees and whether replicates violated the class 1 quartet-nonanomalous assumption (evaluated using expected qCFs). *Importantly, the number of replicates in each bin varies across the model conditions (e.g., just 6% of replicates were anomalous for model condition on the far left compared to 34% anomalous for the far right).*

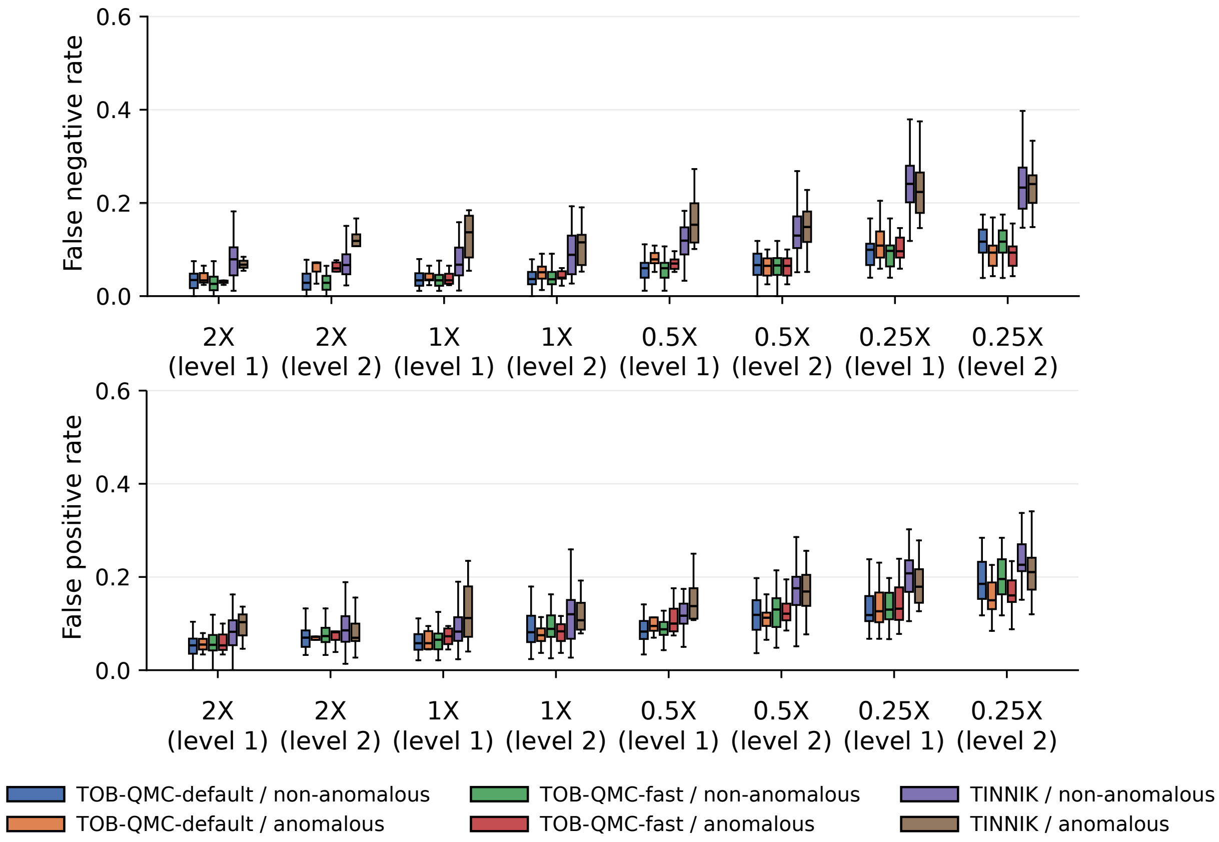

Figure S23: **Impact of violation of class 1 quartet-nonanomalous assumption on 100-taxon networks.** Model condition on $x$-axis indicates network level and branch scaling factor (lower factor is higher ILS). Boxplots indicate method used for TOB reconstruction given estimated gene trees and whether replicates violated the class 1 quartet-nonanomalous assumption (evaluated using expected qCFs). *Importantly, the number of replicates in each bin varies across the model conditions (e.g., just 6% of replicates were anomalous for model condition on the far left compared to 50% anomalous for the far right).*

#### 2.4 Analysis of bee subfamily *Nomiinae*

We re-analyzed a phylogenomic data set for bees, specifically, the subfamily *Nomiinae* [6]. The data set, available on Dryad[4], was recently analyzed for signals of hybridization by [4]. We downloaded the estimated gene trees file[5] from Github[6]. One of the two outgroups `Durfourea_novaeangliae` had been removed from each gene tree, leaving 31 taxa in total. TINNiK and TOB-QMC were run on these gene trees setting $\alpha \in \{0, 10^{-9}, 10^{-8}, 10^{-7}, 10^{-6}, 10^{-5}, 10^{-4}, 10^{-3}\}$ and $\beta \in \{0.9, 0.95, 1\}$. Since some of the taxa are missing in some gene trees and TINNiK will remove some of them, the output TOB produced by TINNiK contained only 25 taxon (see the below log message).

```
    Loading required package: ape
Loading required package: phangorn
Analyzing 25 taxa: Acunomia_melanderi, Afronomia_circumnitens, Austronomia_australica,
    Curvinomia_chalybeata, Dieunomia_heteropoda, Dieunomia_triangulifera, Hoplonomia_elliotii,
    Lasioglossum_albipes, Lipotriches_collaris, Lipotriches_justiciae, Macronomia_clavisetis,
    Nomiapis_bispinosa, Nomiapis_diversipes, Pachynomia_amoenula, Pachynomia_tshibindica,
    Pseudapis_cinerea, Pseudapis_kenyensis, Pseudapis_oxybeloides, Pseudapis_pandeana,
    Pseudapis_riftensis, Pseudapis_siamensis, Steganomus_junodi, Stictonomia_aliceae,
    Stictonomia_sangaensis, Stictonomia_schubotzi
Counting occurrences of displayed quartets for 12650 four-taxon subsets of 25 taxa across 852
    gene trees.
Applying hypothesis test for model T3 to 12650 quartets.
Applying hypothesis test for star tree model to 12650 quartets.
Warning message:
In quartetTable(genetrees, taxanames, epsilon = epsilon) :
  Some taxa missing from some trees.
```

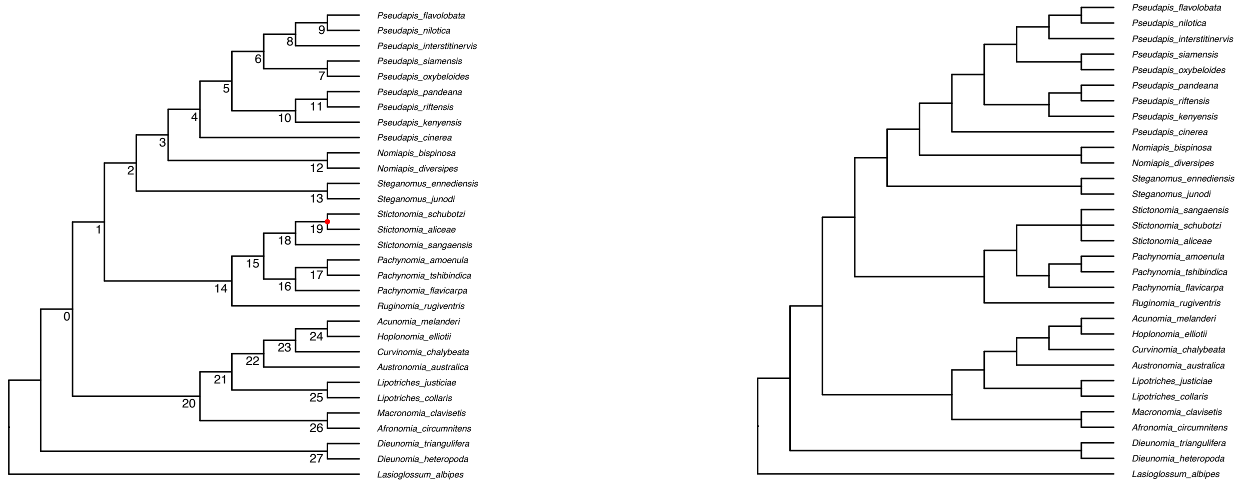

(a) Base tree

(b) Tree of blobs with $\alpha = 10^{-6}$ and $\beta = 0.95$.

Figure S24: **TOB-QMC-3f1a analysis of bee subfamily *Nomiinae*.** Red dots indicate contraction with $\alpha = 10^{-6}$. See meta-data in Table S3 by branch numbers.

[4] https://datadryad.org/dataset/doi:10.5061/dryad.z08kprrb6

[5] Iq2_GTRG_concatenated_gene_tree_files.tre_remove_Dufourea_novaeangliae.txt

[6] https://github.com/mbjorner/hybrid-detection-comparison/tree/main/scripts/Bees_real_dataset

Table S3: **Branch information for TOB-QMC-3f1a analysis of the bee subfamily *Nomiinae*.** Star indicate branch was contracted with $\alpha = 10^{-6}$.

| Branch ID # | T3-test $p$-value | qCFs | norm qCFs | # genes |
|---|---|---|---|---|
| 0 | 0.0617985 | 174/74/53 | 0.58/0.25/0.18 | 301 |
| 1 | 0.00104795 | 526/25/7 | 0.94/0.04/0.01 | 558 |
| 2 | 0.166407 | 232/75/59 | 0.63/0.20/0.16 | 366 |
| 3 | 0.00295547 | 510/52/26 | 0.87/0.09/0.04 | 588 |
| 4 | 0.000163783 | 374/38/12 | 0.88/0.09/0.03 | 424 |
| 5 | 0.0558659 | 143/26/14 | 0.78/0.14/0.08 | 183 |
| 6 | 0.0245771 | 293/50/30 | 0.79/0.13/0.08 | 373 |
| 7 | 0.129982 | 183/27/17 | 0.81/0.12/0.07 | 227 |
| 8 | 0.154292 | 155/89/71 | 0.49/0.28/0.23 | 315 |
| 9 | 2.95346e-05 | 145/82/37 | 0.55/0.31/0.14 | 264 |
| 10 | 0.0148028 | 208/24/10 | 0.86/0.10/0.04 | 242 |
| 11 | 0.0329004 | 139/70/47 | 0.54/0.27/0.18 | 256 |
| 12 | 0.0122178 | 176/18/6 | 0.88/0.09/0.03 | 200 |
| 13 | 0.0560181 | 219/19/9 | 0.89/0.08/0.04 | 247 |
| 14 | 0.0253405 | 73/68/46 | 0.39/0.36/0.25 | 187 |
| 15 | 0.0531949 | 209/22/11 | 0.86/0.09/0.05 | 242 |
| 16 | 0.000233489 | 266/24/5 | 0.90/0.08/0.02 | 295 |
| 17 | 0.00439101 | 240/67/38 | 0.70/0.19/0.11 | 345 |
| 18 | 0.00629214 | 89/55/30 | 0.51/0.32/0.17 | 174 |
| ***19** | **1.36381e-07** | **80/69/21** | **0.47/0.41/0.12** | **170** |
| 20 | 0.0149498 | 127/85/118 | 0.38/0.26/0.36 | 330 |
| 21 | 0.0059578 | 233/225/173 | 0.37/0.36/0.27 | 631 |
| 22 | 0.0251026 | 312/13/4 | 0.95/0.04/0.01 | 329 |
| 23 | 0.0024431 | 330/174/122 | 0.53/0.28/0.19 | 626 |
| 24 | 0.0956415 | 235/145/118 | 0.47/0.29/0.24 | 498 |
| 25 | 0.0407593 | 596/12/4 | 0.97/0.02/0.01 | 612 |
| 26 | 0.1042 | 246/174/145 | 0.44/0.31/0.26 | 565 |
| 27 | 0.0393 | 612/4/0 | 0.99/0.01/0.00 | 616 |

#### 2.5 Analysis of butterfly subfamily *Heliconiinae*

Gene trees for the butterfly subfamily *Heliconiinae* data set were shared with us by the corresponding authors of the original study [7]. In this analysis, the authors reconstructed a species tree with ASTRAL-III and then annotated it with introgression events, determined by testing triplets (rooted triplet trees) displayed by rooted gene trees, with the DCT and branch-length tests implemented in R from [31]. Like this prior analysis (Fig. 2b in [7]), TOB-QMC-3f1a detects introgression

- across the *Melpomene/Silvantiformis* clade (branches 26–46 in Fig. S25)
- impacting 3 species in *Doris* clade (branch 46)
- impacting 3 species in *Dione/Agraulis* clade (branch 54)
- towards root of *Erato* clade (branches 18 19, 24)
- between the *Sara/Sapho* clade and the *Erato* clade (branch 8)

These introgression events above survived the high significance threshold of $\alpha = 10^{-15}$. The original study also detected introgression

- impacting 3 species in *Wallacei* clade (branch 43)
  - min $p$ found for branch 43: $1.48751 \times 10^{-08}$; qCFs: 2710/205/106; normalized qCFs: 0.90/0.07/0.04
- in *Eueides* clade (e.g., branches 48 and 50)
  - min $p$ found for branch 48: 0.000118569; qCFs: 1533/103/55; normalized qCFs: 0.91/0.06/0.03
  - min $p$ found for branch 50: $2.26042e-07$; qCFs: 1003/297/184; normalized qCFs: 0.68/0.20/0.12

The events were significant in the TOB-QMC-3f1 analysis but did not survive the high significance threshold of $\alpha = 10^{-15}$. Importantly, they are are characterized by lower amounts of introgression. To confirm that this was not due to a failure of the 3f1a search, we exhaustively tested all taxa around the quadrapartition induced by branch 43 (i.e., we performed T3 tests on qCFs for *H. burneyi*, *H. wallacei*, *H. egeria*, plus every all other taxa, so 63 tests total). All tests had similar qCFs and $p$-values (Table S5), so the strength of introgression is accurately recovered by the 3f1a algorithm. Overall, this suggests that the $\alpha$ hyperparameter and amount of introgression (inheritance proportion) are important factors to consider in TOB reconstruction. It also highlights the utility of branch annotations, in contrast TINNiK.

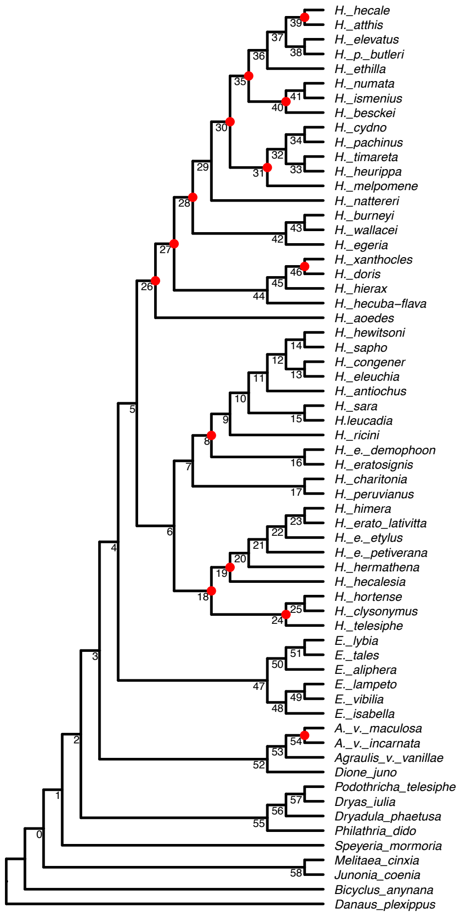

(a) Base tree estimated with TREE-QMC with branches numbered; see meta-data in Table S4.

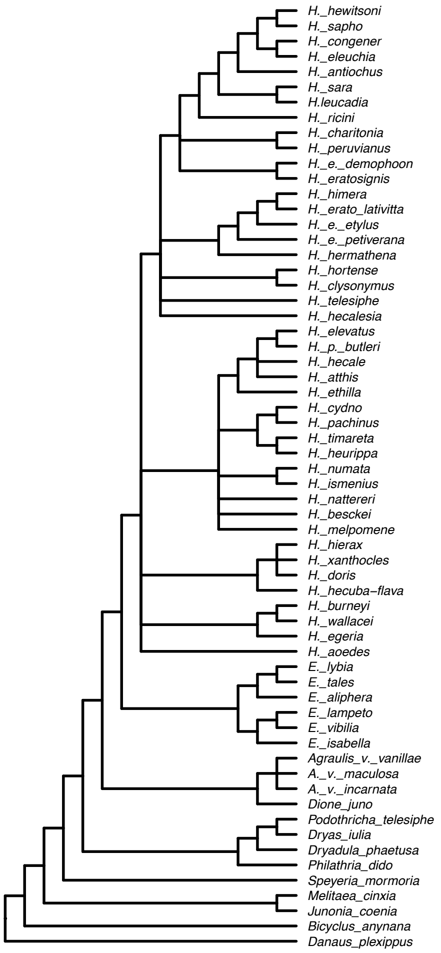

(b) Tree of blobs with $\alpha = 10^{-15}$ and $\beta = 0.95$.

Figure S25: TOB-QMC-3f1a analysis of butterfly subfamily *Heliconiinae*

Table S4: **Branch information for TOB-QMC-3f1a analysis of the butterfly subfamily *Heliconiinae.*** Star indicate branch was contracted with $\alpha = 10^{-15}$.

| Branch ID # | T3-test $p$-value | qCFs | norm qCFs | # genes | taxa |
|---|---|---|---|---|---|
| 0 | 0.0339333 | 1168/315/264 | 0.67/0.18/0.15 | 1747 | Heet/Mcin/Bany/Dple |
| 1 | 0.0881241 | 2431/31/19 | 0.98/0.01/0.01 | 2481 | Hper/Smor/Jcoe/Dple |
| 2 | 0.0171276 | 1969/31/15 | 0.98/0.02/0.01 | 2015 | Elyb/Diul/Smor/Dple |
| 3 | 0.00224253 | 1834/348/272 | 0.75/0.14/0.11 | 2454 | Hnat/Avpe/Ptel/Dple |
| 4 | 0.0127462 | 2159/68/42 | 0.95/0.03/0.02 | 2269 | Hsar/Evib/Avpe/Jcoe |
| 5 | 0.0746376 | 2542/21/11 | 0.99/0.01/0.00 | 2574 | Hleu/Hnat/Evib/Djun |
| 6 | 0.0062224 | 3035/67/39 | 0.97/0.02/0.01 | 3141 | Hert/Hher/Hege/Etal |
| 7 | 0.00453244 | 2118/349/278 | 0.77/0.13/0.10 | 2745 | Hric/Hper/Hhec/Etal |
| ***8** | **1.92268e-26** | **1178/841/460** | **0.48/0.34/0.19** | **2479** | **Hric/Hert/Hper/Hcly** |
| 9 | 1.12164e-07 | 900/882/676 | 0.37/0.36/0.28 | 2458 | Hsap/Hric/Hert/Heet |
| 10 | 0.000497697 | 2546/87/47 | 0.95/0.03/0.02 | 2680 | Hcon/Hsar/Hric/Avcr |
| 11 | 0.0422157 | 2042/341/290 | 0.76/0.13/0.11 | 2673 | Hsap/Hant/Hsar/Hric |
| 12 | 0.0727304 | 1969/382/334 | 0.73/0.14/0.12 | 2685 | Hele/Hhew/Hant/Hric |
| 13 | 0.0531927 | 1765/616/550 | 0.60/0.21/0.19 | 2931 | Hele/Hcon/Hsap/Hbes |
| 14 | 0.0432608 | 960/933/852 | 0.35/0.34/0.31 | 2745 | Hsap/Hhew/Hele/Hric |
| 15 | 0.00065808 | 1585/347/263 | 0.72/0.16/0.12 | 2195 | Hleu/Hsar/Hele/Mcin |
| 16 | 0.0405682 | 3133/27/14 | 0.99/0.01/0.00 | 3174 | Hert/Hdem/Hhew/Htel |
| 17 | 0.184483 | 2651/40/29 | 0.97/0.01/0.01 | 2720 | Hper/Hcha/Hric/Hcyd |
| ***18** | **3.43967e-31** | **1151/1010/554** | **0.42/0.37/0.20** | **2715** | **Hher/Hcly/Hcha/Djun** |
| ***19** | **5.0012e-124** | **1285/1273/355** | **0.44/0.44/0.12** | **2913** | **Hpet/Hhec/Hhor/Hxan** |
| 20 | 0.00185555 | 1577/407/323 | 0.68/0.18/0.14 | 2307 | Hpet/Hher/Hhec/Mcin |
| 21 | 0.0122194 | 1660/208/160 | 0.82/0.10/0.08 | 2028 | Heet/Hpet/Hher/Elyb |
| 22 | 0.00192941 | 1095/849/726 | 0.41/0.32/0.27 | 2670 | Hhim/Heet/Hpet/Hege |
| 23 | 0.350402 | 1231/641/608 | 0.50/0.26/0.25 | 2480 | Hlat/Hhim/Heet/Hpet |
| ***24** | **1.02149e-41** | **1962/685/273** | **0.67/0.23/0.09** | **2920** | **Hhor/Htel/Hhec/Djun** |
| 25 | 0.0288664 | 1429/321/268 | 0.71/0.16/0.13 | 2018 | Hcly/Hhor/Htel/Elam |
| ***26** | **2.6271e-41** | **1594/820/363** | **0.57/0.30/0.13** | **2777** | **Hwal/Haoe/Hhor/Ptel** |
| ***27** | **1.66991e-29** | **1505/914/494** | **0.52/0.31/0.17** | **2913** | **Hwal/Hxan/Haoe/Hsap** |
| ***28** | **1.35845e-18** | **1303/1049/684** | **0.43/0.35/0.23** | **3036** | **Hcyd/Hbur/Hxan/Hhim** |
| 29 | 0.00413092 | 2528/38/17 | 0.98/0.01/0.01 | 2583 | Heth/Hnat/Hege/Dple |
| ***30** | **1.25358e-50** | **1242/1198/577** | **0.41/0.40/0.19** | **3017** | **Hpac/Hbes/Hnat/Htel** |
| ***31** | **3.02167e-39** | **1783/598/228** | **0.68/0.23/0.09** | **2609** | **Hcyd/Hmel/Helv/Jcoe** |
| 32 | 1.1664e-06 | 1315/904/709 | 0.45/0.31/0.24 | 2928 | Htim/Hpac/Hmel/Avfl |
| 33 | 0.00149354 | 1903/589/485 | 0.64/0.20/0.16 | 2977 | Hheu/Htim/Hpac/Hmel |
| 34 | 6.76737e-07 | 1511/804/617 | 0.52/0.27/0.21 | 2932 | Hpac/Hcyd/Hheu/Hmel |
| ***35** | **1.53172e-51** | **1228/1194/570** | **0.41/0.40/0.19** | **2992** | **Hpar/Hism/Hheu/Djun** |
| 36 | 1.94429e-06 | 1506/782/605 | 0.52/0.27/0.21 | 2893 | Hhel/Heth/Hnum/Hnat |
| 37 | 0.00034128 | 1199/893/748 | 0.42/0.31/0.26 | 2840 | Hpar/Hatt/Heth/Hpac |
| 38 | 0.0401207 | 1408/443/384 | 0.63/0.20/0.17 | 2235 | Hpar/Helv/Hatt/Elyb |
| ***39** | **3.32537e-17** | **1415/999/657** | **0.46/0.33/0.21** | **3071** | **Hatt/Hhel/Hpar/Hhim** |
| ***40** | **1.20947e-27** | **1344/1058/615** | **0.45/0.35/0.20** | **3017** | **Hnum/Hbes/Hpar/Hcyd** |
| 41 | 8.15761e-09 | 1210/1060/811 | 0.39/0.34/0.26 | 3081 | Hism/Hnum/Hbes/Hpar |
| 42 | 0.0307561 | 2950/25/12 | 0.99/0.01/0.00 | 2987 | Hwal/Hege/Htim/Eisa |
| 43 | 1.48751e-08 | 2710/205/106 | 0.90/0.07/0.04 | 3021 | Hwal/Hbur/Hege/Dpha |
| 44 | 0.137532 | 2243/45/32 | 0.97/0.02/0.01 | 2320 | Hhie/Hheb/Hcyd/Dple |
| 45 | 1.53875e-05 | 2450/110/55 | 0.94/0.04/0.02 | 2615 | Hxan/Hhie/Hheb/Hbur |
| ***46** | **6.67898e-207** | **1696/941/58** | **0.63/0.35/0.02** | **2695** | **Hdor/Hxan/Hhie/Htel** |
| 47 | 0.00190763 | 1740/18/4 | 0.99/0.01/0.00 | 1762 | Evib/Elyb/Hcly/Dple |
| 48 | 0.000118569 | 1533/103/55 | 0.91/0.06/0.03 | 1691 | Evib/Eisa/Eali/Pdid |
| 49 | 0.334161 | 1586/48/39 | 0.95/0.03/0.02 | 1673 | Evib/Elam/Eisa/Bany |
| 50 | 2.26042e-07 | 1003/297/184 | 0.68/0.20/0.12 | 1484 | Elyb/Eali/Eisa/Hdor |
| 51 | 0.205965 | 1020/208/183 | 0.72/0.15/0.13 | 1411 | Etal/Elyb/Eali/Hlat |
| 52 | 0.00319535 | 2336/31/12 | 0.98/0.01/0.01 | 2379 | Avpe/Djun/Heth/Dple |
| 53 | 0.24 | 2055/4/1 | 1.00/0.00/0.00 | 2060 | Avpe/Avfl/Djun/Elyb |
| ***54** | **3.71573e-35** | **1872/577/231** | **0.70/0.22/0.09** | **2680** | **Avcr/Avpe/Avfl/Hism** |
| 55 | 0.0145563 | 1727/410/343 | 0.70/0.17/0.14 | 2480 | Dpha/Pdid/Hlat/Dple |
| 56 | 2.11402e-13 | 1398/889/606 | 0.48/0.31/0.21 | 2893 | Diul/Dpha/Pdid/Hhel |
| 57 | 8.05961e-12 | 1898/458/274 | 0.72/0.17/0.10 | 2630 | Diul/Ptel/Dpha/Jcoe |
| 58 | 0.0780876 | 1370/44/29 | 0.95/0.03/0.02 | 1443 | Jcoe/Mcin/Eali/Dple |

Table S5: **Branch 43 analysis of the butterfly subfamily *Heliconiinae*.** All 4-taxon samples around the quadrapartition induced by branch 43 are tested (Hbur, Hwal, Hege, and the fourth taxa listed).

| T3-test $p$-value | qCFs | norm qCFs | # genes | taxa |
|---|---|---|---|---|
| 7.573101e-08 | 2696/206/111 | 0.89/0.07/0.04 | 3013 | Hheu |
| 1.020258e-07 | 2758/213/117 | 0.89/0.07/0.04 | 3088 | Htim |
| 1.440484e-07 | 2714/209/115 | 0.89/0.07/0.04 | 3038 | Hpac |
| 1.090846e-07 | 2694/206/112 | 0.89/0.07/0.04 | 3012 | Hcyd |
| 3.740373e-07 | 2737/208/117 | 0.89/0.07/0.04 | 3062 | Hmel |
| 5.772771e-08 | 2718/207/111 | 0.90/0.07/0.04 | 3036 | Helv |
| 1.451381e-07 | 2772/213/118 | 0.89/0.07/0.04 | 3103 | Hpar |
| 5.897829e-08 | 2714/211/114 | 0.89/0.07/0.04 | 3039 | Hatt |
| 6.218247e-07 | 2740/206/117 | 0.89/0.07/0.04 | 3063 | Hhel |
| 7.1535e-06 | 2605/183/107 | 0.90/0.06/0.04 | 2895 | Heth |
| 1.117222e-07 | 2767/214/118 | 0.89/0.07/0.04 | 3099 | Hnum |
| 4.068414e-07 | 2762/209/118 | 0.89/0.07/0.04 | 3089 | Hism |
| 7.374688e-07 | 2707/204/116 | 0.89/0.07/0.04 | 3027 | Hbes |
| 4.420594e-07 | 2747/210/119 | 0.89/0.07/0.04 | 3076 | Hnat |
| 4.420594e-07 | 2748/210/119 | 0.89/0.07/0.04 | 3077 | Hdor |
| 6.679103e-07 | 2748/211/121 | 0.89/0.07/0.04 | 3080 | Hxan |
| 1.726718e-06 | 2392/170/93 | 0.90/0.06/0.04 | 2655 | Hhie |
| 6.819638e-07 | 2681/199/112 | 0.90/0.07/0.04 | 2992 | Hheb |
| 2.271409e-06 | 2685/206/121 | 0.89/0.07/0.04 | 3012 | Haoe |
| 1.399025e-06 | 2756/208/121 | 0.89/0.07/0.04 | 3085 | Hcon |
| 4.133021e-06 | 2713/197/116 | 0.90/0.07/0.04 | 3026 | Hele |
| 3.429197e-07 | 2752/211/119 | 0.89/0.07/0.04 | 3082 | Hhew |
| 1.693279e-07 | 2712/203/111 | 0.90/0.07/0.04 | 3026 | Hsap |
| 1.442384e-06 | 2665/200/115 | 0.89/0.07/0.04 | 2980 | Hant |
| 1.297033e-07 | 2731/204/111 | 0.90/0.07/0.04 | 3046 | Hsar |
| 7.526025e-06 | 2604/188/111 | 0.90/0.06/0.04 | 2903 | Hleu |
| 5.019669e-07 | 2519/180/97 | 0.90/0.06/0.03 | 2796 | Hric |
| 3.142306e-07 | 2769/214/121 | 0.89/0.07/0.04 | 3104 | Hdem |
| 5.203097e-07 | 2768/212/121 | 0.89/0.07/0.04 | 3101 | Hert |
| 1.316402e-06 | 2706/203/117 | 0.89/0.07/0.04 | 3026 | Hper |
| 1.885672e-07 | 2786/216/121 | 0.89/0.07/0.04 | 3123 | Hcha |
| 1.071472e-07 | 2655/202/109 | 0.90/0.07/0.04 | 2966 | Hhor |
| 3.206422e-06 | 2447/177/100 | 0.90/0.06/0.04 | 2724 | Hcly |
| 3.727091e-07 | 2780/212/120 | 0.89/0.07/0.04 | 3112 | Htel |
| 1.72996e-07 | 2761/215/120 | 0.89/0.07/0.04 | 3096 | Hhim |
| 1.179613e-07 | 2533/203/110 | 0.89/0.07/0.04 | 2846 | Hlat |
| 3.610585e-08 | 2401/181/91 | 0.90/0.07/0.03 | 2673 | Heet |
| 3.142306e-07 | 2769/214/121 | 0.89/0.07/0.04 | 3104 | Hdem |
| 3.429321e-08 | 2702/213/114 | 0.89/0.07/0.04 | 3029 | Hpet |
| 3.727091e-07 | 2765/212/120 | 0.89/0.07/0.04 | 3097 | Hher |
| 3.429197e-07 | 2741/211/119 | 0.89/0.07/0.04 | 3071 | Hhec |
| 1.72377e-07 | 2746/211/117 | 0.89/0.07/0.04 | 3074 | Etal |
| 7.454296e-06 | 2043/147/80 | 0.90/0.06/0.04 | 2270 | Elyb |
| 2.797886e-05 | 1703/108/55 | 0.91/0.06/0.03 | 1866 | Eali |
| 2.70459e-06 | 2473/175/98 | 0.90/0.06/0.04 | 2746 | Evib |
| 1.661133e-05 | 2044/144/80 | 0.90/0.06/0.04 | 2268 | Elam |
| 4.07334e-07 | 2701/201/112 | 0.90/0.07/0.04 | 3014 | Eisa |
| 4.727144e-08 | 2666/205/109 | 0.89/0.07/0.04 | 2980 | Avfl |
| 2.276022e-06 | 2459/177/99 | 0.90/0.06/0.04 | 2735 | Avpe |
| 3.467681e-08 | 2661/202/106 | 0.90/0.07/0.04 | 2969 | Avcr |
| 1.297033e-07 | 2716/204/111 | 0.90/0.07/0.04 | 3031 | Djun |
| 5.275008e-07 | 2677/200/112 | 0.90/0.07/0.04 | 2989 | Ptel |
| 6.263127e-07 | 2675/198/111 | 0.90/0.07/0.04 | 2984 | Diul |
| 1.487508e-08 | 2710/205/106 | 0.90/0.07/0.04 | 3021 | Dpha |
| 7.417329e-07 | 2679/200/113 | 0.90/0.07/0.04 | 2992 | Pdid |
| 1.239242e-06 | 2549/194/110 | 0.89/0.07/0.04 | 2853 | Smor |
| 1.789127e-07 | 2410/181/95 | 0.90/0.07/0.04 | 2686 | Jcoe |
| 4.477547e-07 | 2095/157/80 | 0.90/0.07/0.03 | 2332 | Mcin |
| 7.48532e-07 | 2128/158/82 | 0.90/0.07/0.03 | 2368 | Bany |
| 7.289211e-06 | 2430/179/104 | 0.90/0.07/0.04 | 2713 | Dple |

#### 2.6 Analysis of seed plants

Recently, Kolbow *et al.* reanalyzed a seed plant phylogenomic data set to examine a conflict involving four major gymnosperm clades as well as four major fern clades [16]. They used TINNiK to estimate a TOB and found the resulting tree to be highly unresolved with $\alpha = 0.001$ (Fig. 5 in Supplementary Materials of [16]). To reanalyze the data, we downloaded the estimated gene trees from the OneKP project[7] and then restricted to the same set of 96 taxa as [16]. At $\alpha = 0.001$, TOB-QMC contracts 25 branches. Some of the events branches 6 and 7 are deeper in the tree and do not seem very well supported based on the qCFs are 220/27/3 and 179/26/2 (although the min $p$ values found are less than $10^{-5}$). Only three branches are contracted at the $\alpha = 10^{-15}$

- Subclade *Polypodium*, genus of ferns (branch 25)
  - min $p$ found for branch 25: $4.67094 \times 10^{-20}$; qCFs: 149/134/25; normalized qCFs: 0.48/0.44/0.08
- Root of *Pinales*, order of extant conifers (branch 76)
  - min $p$ found for branch 76: $1.25474 \times 10^{-16}$; qCFs: 145/124/27; normalized qCFs: 0.49/0.42/0.09
- Subclade of *Lycophytes*, a group of vasucular plants, including clubmosses (branch 88)
  - min $p$ found for branch 88: $2.89384 \times 10^{-16}$; qCFs: 127/101/17; normalized qCFs: 0.52/0.41/0.07

It is also worth noting that other factors impacting plant evolution, e.g. differences in ploidy, may also challenge accurate TOB reconstruction. Future work should explore this further.

[7] https://de.cyverse.org/anon-files/iplant/home/shared/commons_repo/curated/oneKP_capstone_2019/alignments_and_trees/genetrees/Best.FAA.tre

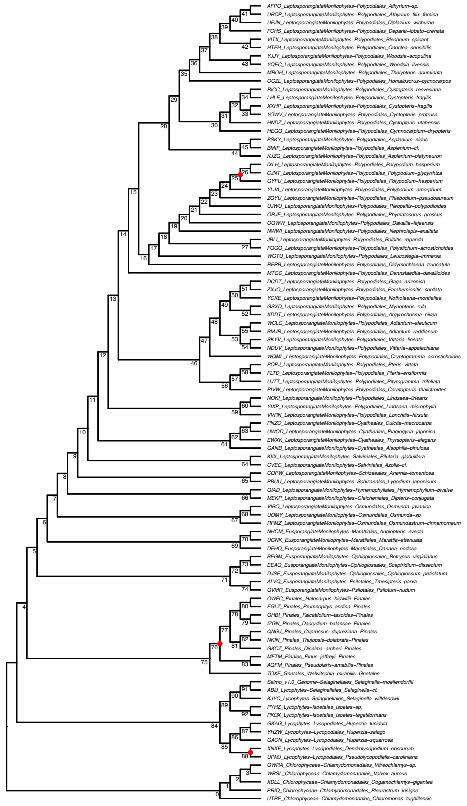

(a) Base tree via TREE-QMC with branches numbered; see meta-data in Table S6–S7.

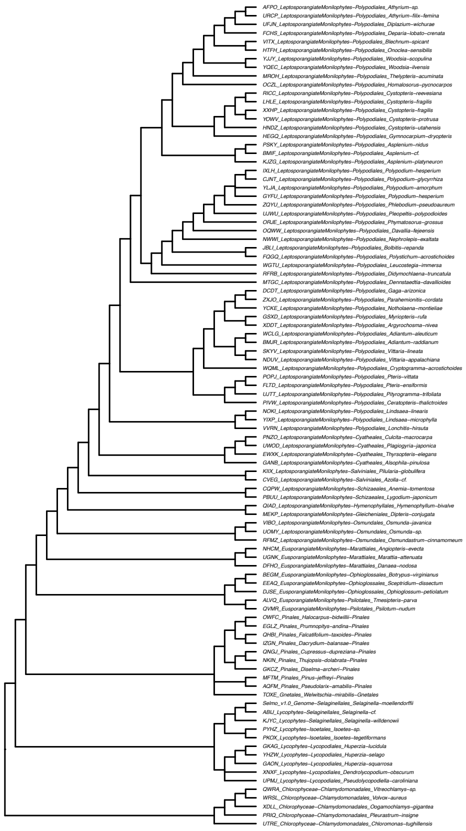

(b) Tree of blobs with $\alpha = 10^{-15}$ and $\beta = 0.95$.

Figure S26: TOB-QMC Analysis of seed plants

Table S6: **Branch information for TOB-QMC-3f1a analysis of the seed plants.** Star and dagger indicate branch is contracted with $\alpha = 10^{-15}$ and $\alpha = 0.001$, respectively.

| Branch ID # | T3-test $p$-value | qCFs | norm qCFs | # genes |
|---|---|---|---|---|
| 0 | 1 | 190/0/0 | 1.00/0.00/0.00 | 190 |
| 1 | 0.0174576 | 40/39/23 | 0.39/0.38/0.23 | 102 |
| 2 | 0.0426941 | 46/37/22 | 0.44/0.35/0.21 | 105 |
| 3 | 0.0385924 | 63/20/9 | 0.68/0.22/0.10 | 92 |
| 4 | 0.0146863 | 97/30/14 | 0.69/0.21/0.10 | 141 |
| 5 | 0.00881743 | 181/11/2 | 0.93/0.06/0.01 | 194 |
| †6 | 9.22048e-06 | 81/72/29 | 0.45/0.40/0.16 | 182 |
| †7 | 2.60994e-06 | 220/27/3 | 0.88/0.11/0.01 | 250 |
| †8 | 7.80087e-07 | 179/26/2 | 0.86/0.13/0.01 | 207 |
| 9 | 0.0385924 | 132/20/9 | 0.82/0.12/0.06 | 161 |
| †10 | 0.000644052 | 184/20/4 | 0.88/0.10/0.02 | 208 |
| †11 | 7.12951e-06 | 105/90/40 | 0.45/0.38/0.17 | 235 |
| †12 | 3.93147e-09 | 128/25/0 | 0.84/0.16/0.00 | 153 |
| 13 | 0.0272956 | 112/31/16 | 0.70/0.19/0.10 | 159 |
| 14 | 0.0015797 | 121/88/51 | 0.47/0.34/0.20 | 260 |
| †15 | 0.000394211 | 146/27/7 | 0.81/0.15/0.04 | 180 |
| †16 | 7.66417e-08 | 87/69/20 | 0.49/0.39/0.11 | 176 |
| 17 | 0.00620988 | 56/54/31 | 0.40/0.38/0.22 | 141 |
| 18 | 0.00611406 | 81/81/53 | 0.38/0.38/0.25 | 215 |
| 19 | 0.0110568 | 98/38/19 | 0.63/0.25/0.12 | 155 |
| 20 | 0.091631 | 128/27/16 | 0.75/0.16/0.09 | 171 |
| 21 | 0.00679837 | 172/21/7 | 0.86/0.10/0.04 | 200 |
| 22 | 0.0103728 | 108/65/39 | 0.51/0.31/0.18 | 212 |
| 23 | 0.147458 | 70/57/43 | 0.41/0.34/0.25 | 170 |
| 24 | 0.021165 | 142/20/8 | 0.84/0.12/0.05 | 170 |
| *25 | 4.67094e-20 | 149/134/25 | 0.48/0.44/0.08 | 308 |
| †26 | 2.98261e-06 | 144/108/50 | 0.48/0.36/0.17 | 302 |
| 27 | 0.213092 | 52/39/29 | 0.43/0.33/0.24 | 120 |
| †28 | 0.000164111 | 137/95/50 | 0.49/0.34/0.18 | 282 |
| 29 | 0.0100091 | 87/56/32 | 0.50/0.32/0.18 | 175 |
| 30 | 0.102718 | 130/24/14 | 0.77/0.14/0.08 | 168 |
| 31 | 0.0848273 | 195/12/5 | 0.92/0.06/0.02 | 212 |
| †32 | 1.44799e-07 | 103/75/24 | 0.51/0.37/0.12 | 202 |
| 33 | 0.284224 | 180/32/24 | 0.76/0.14/0.10 | 236 |
| 34 | 0.138549 | 100/39/27 | 0.60/0.23/0.16 | 166 |
| 35 | 0.0684428 | 69/66/50 | 0.37/0.36/0.27 | 185 |
| 36 | 0.0499697 | 78/65/45 | 0.41/0.35/0.24 | 188 |
| 37 | 0.0108879 | 85/83/56 | 0.38/0.37/0.25 | 224 |
| 38 | 0.00125639 | 60/51/24 | 0.44/0.38/0.18 | 135 |
| †39 | 2.62339e-05 | 85/74/32 | 0.45/0.39/0.17 | 191 |
| 40 | 0.0287487 | 123/62/40 | 0.55/0.28/0.18 | 225 |
| 41 | 0.158712 | 234/16/9 | 0.90/0.06/0.03 | 259 |
| †42 | 3.44333e-05 | 85/76/34 | 0.44/0.39/0.17 | 195 |
| 43 | 0.113789 | 213/35/23 | 0.79/0.13/0.08 | 271 |
| 44 | 0.00871 | 250/6/0 | 0.98/0.02/0.00 | 256 |
| 45 | 0.103491 | 172/20/11 | 0.85/0.10/0.05 | 203 |
| †46 | 9.42142e-05 | 243/11/0 | 0.96/0.04/0.00 | 254 |

Table S7: **Branch information for TOB-QMC-3f1a analysis of the seed plants, cont.** Star and dagger indicate branch is contracted with $\alpha = 10^{-15}$ and $\alpha = 0.001$, respectively.

| Branch ID # | T3-test $p$-value | qCFs | norm qCFs | # genes |
|---|---|---|---|---|
| †47 | 0.000961723 | 107/87/49 | 0.43/0.35/0.20 | 243 |
| †48 | 1.85638e-06 | 174/74/27 | 0.63/0.27/0.10 | 275 |
| 49 | 0.0021481 | 244/27/9 | 0.87/0.10/0.03 | 280 |
| 50 | 0.0904991 | 185/47/32 | 0.77/0.19/0.13 | 264 |
| 51 | 0.0311502 | 207/40/23 | 0.78/0.15/0.09 | 270 |
| 52 | 0.0486603 | 128/46/29 | 0.60/0.22/0.14 | 203 |
| †53 | 1.55462e-06 | 92/87/36 | 0.44/0.42/0.17 | 215 |
| 54 | 0.478 | 279/1/0 | 0.99/0.01/0.00 | 280 |
| 55 | 0.00194314 | 124/78/44 | 0.45/0.28/0.16 | 246 |
| †56 | 6.75293e-08 | 101/98/38 | 0.41/0.40/0.15 | 237 |
| 57 | 0.0317025 | 133/19/8 | 0.83/0.12/0.05 | 160 |
| 58 | 0.0271723 | 117/61/39 | 0.51/0.27/0.17 | 217 |
| 59 | 0.00430026 | 56/54/30 | 0.40/0.39/0.21 | 140 |
| 60 | 0.086 | 163/3/0 | 0.98/0.02/0.00 | 166 |
| †61 | 0.000250813 | 126/47/18 | 0.63/0.24/0.09 | 191 |
| †62 | 0.000461037 | 77/73/38 | 0.41/0.39/0.20 | 188 |
| 63 | 0.14122 | 144/15/8 | 0.83/0.09/0.05 | 167 |
| 64 | 0.0226258 | 155/58/36 | 0.64/0.24/0.15 | 249 |
| 65 | 0.121027 | 144/26/16 | 0.80/0.14/0.09 | 186 |
| †66 | 5.36294e-07 | 77/72/25 | 0.44/0.41/0.14 | 174 |
| 67 | 0.086 | 232/3/0 | 0.99/0.01/0.00 | 235 |
| 68 | 0.132347 | 136/44/31 | 0.62/0.20/0.14 | 211 |
| 69 | 0.086 | 155/3/0 | 0.98/0.02/0.00 | 158 |
| 70 | 0.00416 | 141/7/0 | 0.95/0.05/0.00 | 148 |
| 71 | 0.0038105 | 153/24/8 | 0.83/0.13/0.04 | 185 |
| 72 | 0.0164505 | 206/19/7 | 0.89/0.08/0.03 | 232 |
| 73 | 0.0461386 | 179/10/3 | 0.93/0.05/0.02 | 192 |
| 74 | 0.197 | 216/2/0 | 0.99/0.01/0.00 | 218 |
| 75 | 0.086 | 233/3/0 | 0.99/0.01/0.00 | 236 |
| *76 | 1.25474e-16 | 145/124/27 | 0.48/0.41/0.09 | 296 |
| 77 | 0.0764175 | 207/9/3 | 0.94/0.04/0.01 | 219 |
| 78 | 0.197 | 278/2/0 | 0.99/0.01/0.00 | 280 |
| 79 | 0.00416 | 165/7/0 | 0.96/0.04/0.00 | 172 |
| 80 | 0.0130181 | 81/55/32 | 0.47/0.32/0.19 | 168 |
| 81 | 0.478 | 219/1/0 | 0.99/0.00/0.00 | 220 |
| 82 | 0.0169614 | 187/63/39 | 0.62/0.21/0.13 | 289 |
| 83 | 0.197 | 227/2/0 | 0.99/0.01/0.00 | 229 |
| 84 | 2.42278e-07 | 91/41/7 | 0.68/0.31/0.05 | 139 |
| 85 | 0.0393 | 209/4/0 | 0.98/0.02/0.00 | 213 |
| 86 | 0.197 | 203/2/0 | 0.99/0.01/0.00 | 205 |
| 87 | 0.123008 | 141/14/7 | 0.86/0.09/0.04 | 162 |
| *88 | 2.89384e-16 | 127/101/17 | 0.51/0.41/0.07 | 245 |
| †89 | 4.44927e-07 | 141/69/22 | 0.63/0.31/0.10 | 232 |
| 90 | 0.197 | 148/2/0 | 0.99/0.01/0.00 | 150 |
| 91 | 0.0393 | 207/4/0 | 0.98/0.02/0.00 | 211 |
| 92 | 0.086 | 191/3/0 | 0.98/0.02/0.00 | 194 |

[16] Nathan Kolbow, Sungsik Kong, and Claudia Solís-Lemus. A method for massively scalable phylogenetic network inference. *bioRxiv*, 2025.

[33] Yufeng Wu. Inference of population admixture network from local gene genealogies: a coalescent-based maximum likelihood approach. *Bioinformatics*, 36(Supplement_1):i326–i334, 2020.

[34] Yun Yu, Jianrong Dong, Kevin J. Liu, and Luay Nakhleh. Maximum likelihood inference of reticulate evolutionary histories. *Proceedings of the National Academy of Sciences*, 111(46):16448–16453, 2014.
